# UNICORN: Towards Universal Cellular Expression Prediction with a Multi-Task Learning Framework

**DOI:** 10.1101/2025.01.22.634371

**Authors:** Tianyu Liu, Tinglin Huang, Lijun Wang, Yingxin Lin, Rex Ying, Hongyu Zhao

## Abstract

Sequence-to-function analysis is a challenging task in human genetics, especially in predicting cell-type-specific multi-omic phenotypes from biological sequences such as individualized gene expression. Here, we present UNICORN, a new method with improved prediction performances than the existing methods. UNICORN takes the embeddings from biological sequences as well as external knowledge from pre-trained foundation models as inputs and optimizes the predictor with carefully-designed loss functions. We demonstrate that UNICORN outperforms the existing methods in both gene expression prediction and multiomic phenotype prediction at the cellular level and the cell-type level, and it can also generate uncertainty scores of the predictions. Moreover, UNICORN is able to link personalized gene expression profiles with corresponding genome information. Finally, we show that UNICORN is capable of characterizing complex biological systems for different disease states or perturbations. Overall, embeddings from foundation models can facilitate the understanding of the role of biological sequences in the prediction task, and incorporating multi-omic information can enhance prediction performances.

## 1 Introduction

Modelling DNA and Central Dogma is a challenging but essential task to biological research [1–7]. One key question is how to predict the expression of products from sequences (e.g., DNA sequences [8] and amino acid sequences [9, 10]). A good predictor can help us uncover the relationship between genetic information and cellular functions, which cannot be directly achieved by using previous sequence-to-expression models due to the difference of training data [11]. With the accumulation of single-cell sequencing data [12–14], it is possible to impute and analyze cell-specific or cell-type-specific functional information based on sequence models, an exciting direction to link genomics and transcriptomics/proteomics with a unified framework to advance genetic research.

Single-cell sequencing (including scRNA-seq [12, 13], scATAC-seq [15], and CITE-seq [16]) generates different types of measurement at the single-cell resolution. The knowledge gained from downstream analysis, such as cell-type or cell-cycle information, may raise biological questions to explore. Although single-cell data allow us to capture the omic information at the single-cell resolution, there is large noise in these data, which presents a major obstacle for data analysis and interpretation [17]. To tackle this problem, [18] considers aggregating single-cell data into pseudo-bulk data using metadata information of cells. Such approach is widely used in inferring cell-type-specific expression quantitative trait loci (eQTLs) [19] or evaluations for deconvolution [20]. However, the effectiveness of pseudo-bulk settings in sequence-to-expression tasks has not yet been discussed in detail. Meanwhile, these predictors may also have biases for certain cell types or diseases [21, 22], which motivates us to explore the reasons behind such performance differences.

Existing methods for gene expression prediction based on DNA sequences relied on either CAGE-seq results (e.g., Basenji2 [23], Enformer [21]) or bulkRNA-seq results (e.g., Borzoi [24]) as targets for training. Most of them were developed before significant progress was made in single-cell atlas, and the cost of bulk-level data is less than sequencing single-cell data [25]. Nvwa [26] attempted to predict whether a gene will express in cells using single-cell transcriptomics, but non-human data were considered, and classification is not a proper problem setting for gene expression prediction because it is important to evaluate the expression levels. Recently, seq2cells [11] built on the pre-trained Enformer model to extract embeddings from DNA sequences to predict gene expression across all the cells, whereas scooby [27] used a similar idea for single-cell multi-omic data. Moreover, researchers are also interested in exploring the modeling of DNA language based on self-supervised learning [28] (SSL) approach (e.g., HyenaDNA [2], DNABERT [29], Evo [30]). However, these methods have specific limitations. First, models similar to Enformer did not consider enough cell-type specific information, and thus, the heterogeneity of gene expression at the cell level cannot be fully modeled. Second, seq2cells did not discuss the choices of models for generating sequence embeddings, nor did it discuss the selection process of targets for prediction. The prediction of single-cell gene expression is also not practical because of the experimental noise. Third, SSL-based DNA language models did not always show better performance in the evaluations of downstream tasks [31], and few are applied for cell-type-level gene expression prediction. Finally, there are some technical limitations in constructing a sequence-to-expression model, such as how to model meaningful sequence context and how to reflect the individual-level effects in the prediction results. Therefore, we need a flexible and transparent framework to answer these questions.

In this manuscript, we introduce a novel transfer-learning-based framework for predicting multi-omic expression levels from their corresponding sequence information, called a **UNI**versal **C**ell expressi**O**n p**R**edictio**N** framework (UNICORN). We utilize a general genetic language model [32] plus a general large language model (LLM) [33] as an option to generate the embeddings of sequences and predict the corresponding expression levels (also defined as phenotypes in our context [34]) of such sequences (genes, peaks, or proteins) in different cell types at the pseudo-bulk level. Our framework is explainable with the help of uncertainty prediction for the expression levels of different sequences and multi-task-learnable with the help of different types of omic data used for training and testing. UNICORN is a powerful framework to understand the relationships between 1. biological sequence and cell-type-specific effects, 2. biological sequence and individual genetic effects, and 3. biological sequence and disease/perturbation-specific effects.

## 2 Results

### Overview of UNICORN

Our model predicts different types of omic data based on their corresponding biological sequence information. We first infer the embeddings of target biological sequences with corresponding language models (genomic language models (gLMs), protein language models (PLMs), or LLMs) and save the embeddings **e** in a separate file. We then construct the training and testing datasets based on the target omic type and train a non-linear predictor 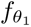 to model the relationship between sequence and expression information by predicting the expression-level information from the sequence-level information. Meanwhile, we have an uncertainty estimator to quantify the uncertainty of each prediction. The difference between UNICORN and other baseline models is summarized in Supplementary Figure 1, which demonstrates the unique contributions of our model design. Our metrics include the correlation and distance between the observed and the predicted expression levels, and our model is capable of multiple downstream applications, including the analysis of genetic effects at the cell-type levels, individual levels, disease-specific levels, and perturbation-specific levels. The landscape of UNICORN is summarized in Figures 1 (a)-(c). Details of UNICORN are presented in the Methods section.

**Fig. 1:**
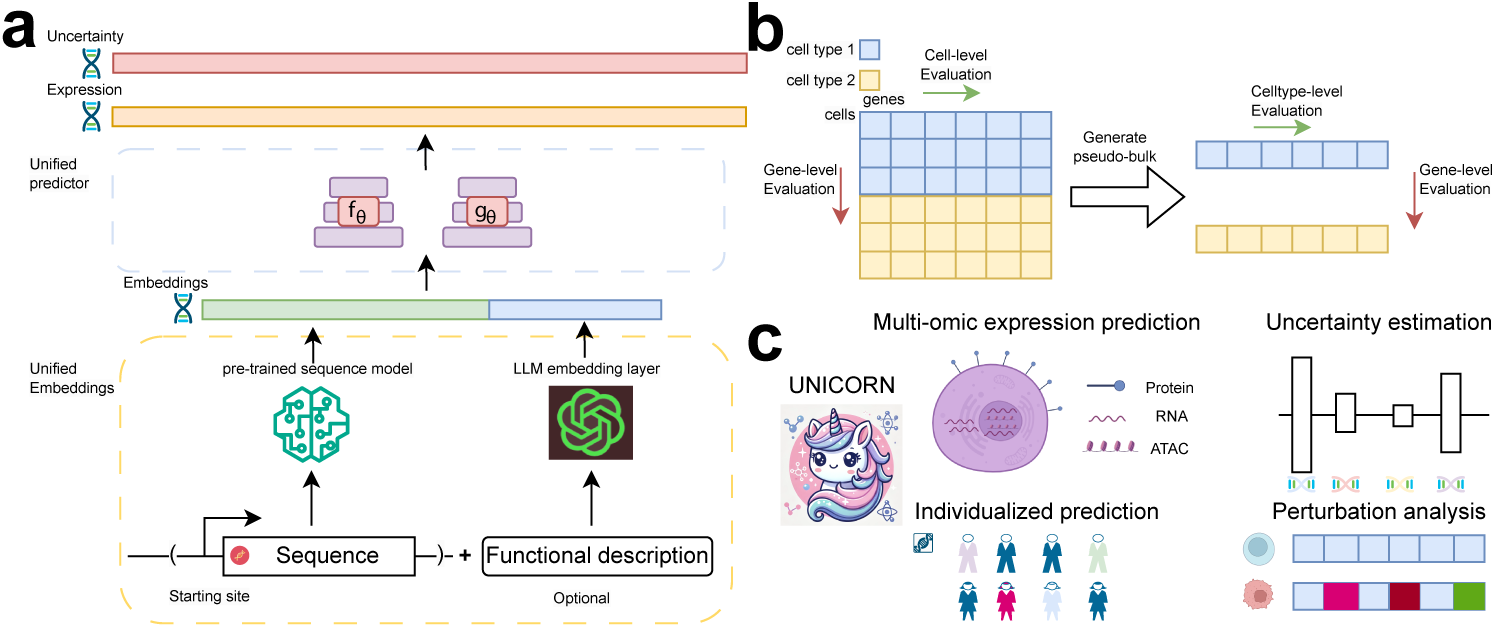
Workflow and functions of UNICORN. (a) The architecture and components of UNICORN. Our method accepts embeddings from multi-modal sources, predicts expression levels at the single-cell or cell-type resolutions, and estimates the uncertainty of model outputs. (b) Explanations of our evaluation framework. We consider evaluations from two perspectives (per-cell and per-gene) and two modes (single-cell level and pseudo-bulk level). (c) A summary of UNICORN’s unique functions, including multi-omic expression prediction, uncertainty estimation, individualized prediction, and perturbation prediction. The logo is generated with DALLE [35].

### Exploring the capacity of UNICORN for expression prediction

Here we explore the performance of UNICORN towards predicting gene expression levels at the single-cell or cell-type level. This task is the basic component of our expression prediction problem and is helpful for us in analyzing the capacity of sequence-based predictors. To predict the expression levels of each cell, we referred to the original design of Enformer to collect sequence information of ∼200 kb length centered at the transcription start site (TSS) for each gene. We used the centered sequence to generate the gene embeddings as inputs and single-cell gene expression levels as outputs. We split a single-cell dataset into a training set (70%), a validation set (20%), and a testing set (10%) using gene names and chose mean squared error (MSE) and Pearson correlation coefficients (corr) as metrics for evaluations [11, 36]. Our evaluations considered both genes and cells as dimensions and observed expression levels in the pseudo-bulk mode. For the models we evaluated, we chose baselines including Enformer (enformer), Borzoi (borzoi), seq2cells, UNICORN with gene embeddings from Enformer (UNICORN), gene embeddings from HyenaDNA (UNICORN hyenadna), gene embeddings from Borzoi (UNICORN borzoi), gene embeddings from Nucleotide Transformers [37] (UNICORN nt), gene embeddings from DNABERT (UNICORN dnabert), gene embeddings from Gene2vec [38] (UNICORN gene2vec), gene embeddings from LLMs [39](UNICORN genept), and gene embeddings from both Enformer and LLMs (UNICORN comb). We considered two datasets for testing, including scRNA-seq data from Thymus (thymus) and scRNA-seq data from PBMC (pbmc). Details of our training settings, methods, and metrics can be found in the Methods section.

We present the prediction results at the single-cell resolution in Figures 2 (a)-(d). All the methods are ranked by the mean values of their evaluation results in ascending order from left to right. Different figures correspond to different evaluation approaches. For the thymus dataset, UNICORN comb had the highest correlations and lowest MSE. Figure 2 (a) shows the comparisons based on correlations at the gene level, and we found that using more advanced models to encode sequence information (for example, HyenaDNA) did not yield better results compared with the default mode of UNICORN. Moreover, the other methods (UNICORN gene2vec, UNICORN comb, seq2cells, and UNICORN) did not show obvious differences by considering the distribution of correlations. However, the UNICORN and its combination mode had smaller averaged MSE according to Figure 2 (b), and thus a good approach to integrating gene embeddings from different modalities effectively can enhance prediction performances. Furthermore, we visualize the results with gene-level evaluations of the PBMC dataset in Figures 2 (c) and (d). For this dataset, UNICORN and UNICORN combine had high correlation scores and UNICORN had the lowest MSE. We found that UNICORN with gene2vec, genept, and combined information as sources of gene embeddings had larger averaged MSE, while full-parameter fine-tuning based on Enformer and Borzoi did not perform well for both datasets. The results of cell-level metrics of these two datasets are shown in Supplementary Figures 2 (a)-(d), and UNICORN was always among the top two performers across different cell-level metrics. To further support the strength of our model design, we compared UNICORN with Decima [22], a model pre-trained with large-scale single-cell transcriptomic data for expression prediction, and found that our method outperformed Decima in the evaluation of average MSE and gene-level correlation, shown in Supplementary Figure 3. Therefore, gene embeddings from sequences worked as a better input for predicting gene expression levels, compared with information directly from co-expression relationships or text mining. One possible reason to explain the contribution of gene embeddings for prediction is the positive effect of decoupling training [40]. In conclusion, our experimental results demonstrate that sequence information can serve as a better starting point for analyzing gene expression at the cellular level.

**Fig. 2:**
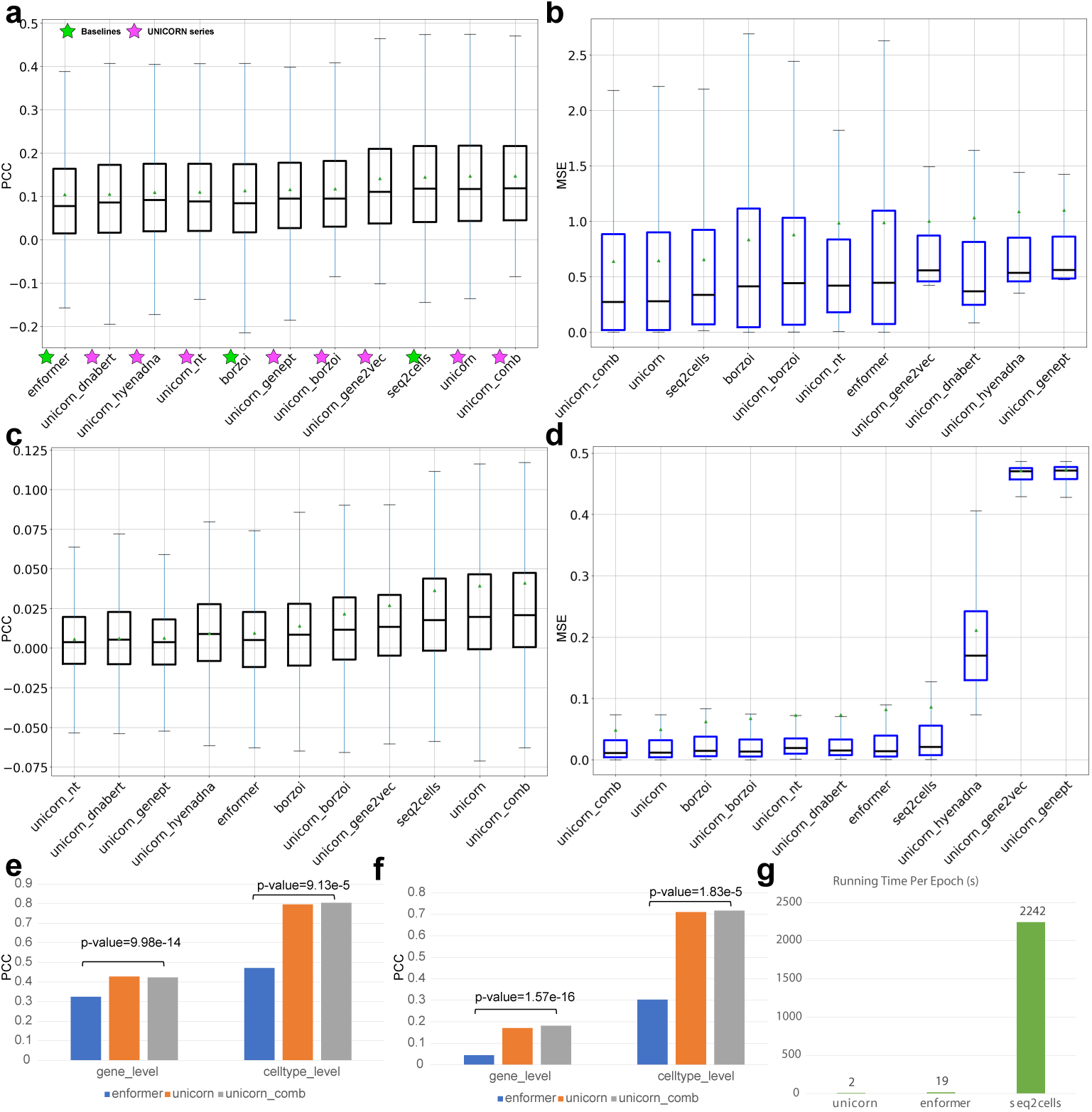
Comprehensive evaluations for gene expression predictions from sequences. The triangle shape represents the mean value, and the black dashed line represents the median value. We report the results based on box plots. (a) Results of gene-level correlations across different methods based on the thymus dataset. (b) Results of MSE across different methods based on the thymus dataset. (c) Results of gene-level correlations across different methods based on the PBMC dataset. (d) Results of MSE across different methods based on the PBMC dataset. (e) Results of gene-level correlations with pseudo-bulk evaluation based on the thymus dataset. (f) Results of gene-level correlations with pseudo-bulk evaluation based on the PBMC dataset. (g) Training time per epoch of different models. The unit is second (s).

We also performed biology-oriented evaluation to demonstrate the superiority of UNICORN for predicting meaningful gene expression levels. According to Supplementary Figures 4 (a) and (b), UNICORN ranked in the top models for predicting the expression levels of cell-type marker genes in Thymus dataset. Within the same dataset, we also computed clustering scores with predicted transcriptomic information at the cell level to examine the capacity of UNICORN and other models in distinguishing different cell types and generating meaningful cell clusters. According to our results in Supplementary Figure 5, UNICORN still shows strong performance in representing cells from different cell types, supported by the high clustering scores by either metric-specific or average evaluations. Finally, we tested the proportion of extracted Gene Ontology (GO) pathways with the genes from different approaches with good prediction. To select important genes, we first removed the genes whose PCC is lower than the mean PCC or MSE is higher than the mean MSE from different methods, and run GO pathway analysis based on these gene sets. We then computed the proportion of significant pathways discovered in these gene sets. According to Supplementary Figure 6, methods from the series of UNICORN still present leading performances in predicting gene expression levels with biological information.

We improved the training efficiency compared with seq2cells and Enformer, shown in Figure 2 (g). We controlled the batch size as 1024 for different models to make a fair comparison and UNICORN had the fastest training speed.

However, we note that the correlations measured at the single-cell level were generally low, especially for the PBMC dataset. Most methods did not have prediction results with an average correlation value higher than 0.05. This is largely due to the high noise in single-cell data measurements. Therefore, we aggregated single-cell data into pseudo-bulk data by cell types and visualized the results in Figures 2 (e) and (f). Based on these two figures, correlations for both gene and cell levels increased and UNICORN could better predict gene expressions at the cell type level with a significant improvement over Enformer based on the two-sided Mann–Whitney U test. Furthermore, UNICORN with combination embeddings as input data outperformed the default mode in general. Therefore, our conclusions for the single-cell resolution also hold for the cell-type resolution.

We further evaluated the capacity of predicting cell-type-level expression profiles with UNICORN based on two atlas-level scRNA-seq datasets, including the ROSMAP dataset from human brain tissue and the OneK1K dataset from human PBMC. We updated the baseline methods with dofferemt models, including SCOVER (scover) [41] and DanQ [42], which are based on the motif-captured design; LassoNet [43], which works as a feature-selection-driven method for multi-target regression; and GET [44], which is a foundation model for predicting gene expression levels from chromatin accessibility across different cell types. We retrained the previous three baseline methods with selected datasets for comparison. To compare GET, as it provided predicted gene expression profiles for different cell types, we directly took the predicted results for comparison. We utilized the same PCC and MSE as metrics to perform gene-level evaluation and cell-type-level evaluation. The results are shown in Supplementary Figures 7 (a)-(d) based on the OneK1K [45] dataset and Supplementary Figures 8 (a)-(d) based on the ROSMAP [46] dataset. According to these figures, UNICORN outperforms all selected baseline methods except GET for cell-level correlations. Our method also captured gene-level signals better than GET for both the ROSMAP and OneK1K datasets and was close to GET in capturing cell-type-level signals. Since GET was pretrained with data from ATAC-seq, it had a high correlation with certain transcriptomic profiles [47, 48].

We also found that more cells for training can improve the predictors, even if our target is to predict gene expressions at the cell-type level. As shown in Supplementary Figure 9, increasing the number of cells for training also increased the gene-level correlations. We also compared the prediction results between the model input as pseudo-bulk information and the model input as single-cell information to further support our conclusions, shown in Supplementary Figures 10 (a) and (b), for these two datasets. We investigated the selected sequence length’s effect on prediction performances, summarized in Appendix A. Our model can also predict gene expression levels for atlas-level datasets, which is further discussed in Appendix B.

### Uncovering the complexity of model training

Explainability is an important part of understanding how neural networks work. In this section, we utilize function-based explainability to understand the factors that affect the performance of predictions using the model’s output based on different scenarios [49]. Here, we discuss the first component of explainability, which helps us understand the behavior of gene expression prediction. In Figure 3 (a) with stages ① -③, we illustrate the differences in model performance under different settings. We can adopt many ideas to improve model performance, and the basic approach is to tune hyper-parameters and test a better loss function design. Such design can explain the performance improvement in ①. Secondly, we consider addressing the sparsity problem existing in single-cell datasets. In the past, seq2cells utilized scVI [50] to impute the scRNA-seq dataset as a denoising step. However, such design is biased to the imputation method because we found that using different denoising methods led to different results, shown in Supplementary Figure 11 including the comparison between scVI and DeepImpute [51]. Therefore, handling sparsity might not fully address this problem, so we standardized the single-cell dataset using z-norm for each gene to reduce the existence of zeros as the neural network is hard to learn and predict zeros in the regression problem [52]. Such design can explain the performance improvement in ②. Finally, the samples in the single-cell dataset can be treated as replicates of specific cell types, and predicting single-cell gene expression is still extremely difficult since our previous improvement is insufficient. Therefore, we evaluate the outputs of UNICORN by generating the pseudo-bulk dataset aggregated by cell types and visualize the result in Figure 3 (a). We found that this method led to noticeable improvement, and the performance gap in ③ can be explained by data complexity. Furthermore, we also tried to explore the linkage between gene functions and expression prediction performance, which are further discussed in Appendix C.

**Fig. 3:**
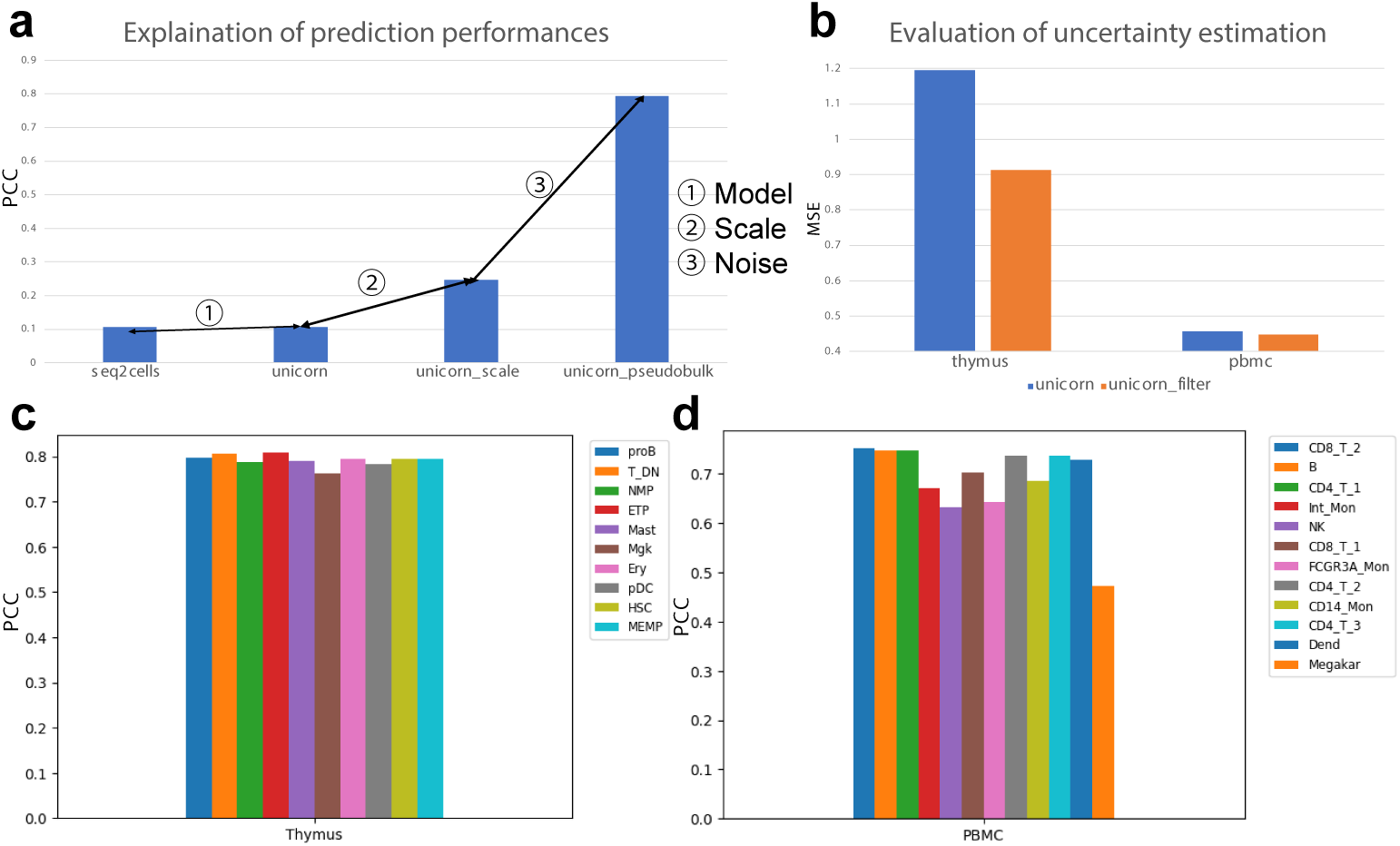
Explainability of UNICORN helps understand the principles of expression prediction. (a) Explainability of UNICORN from the differences of pre-processing steps and post-processing steps. (b) Explainability of UNICORN from uncertainty estimation. (c) Average correlation coefficients across different cell types for the thymus dataset. (d) Average correlation coefficients across different cell types for the PBMC dataset.

In conclusion, sequence-based models are promising for predicting cell-type-specific gene expression levels powered by methods from deep learning.

### Improving prediction performance by uncertainty estimation

Here, we discuss the second component of explainability, which facilitates the selection of more informative genes inspired by the uncertainty estimation framework [53]. Our idea is to estimate the uncertainty levels of each sample to identify the reliability of model prediction. We computed the uncertainty of each gene and each cell based on the loss prediction method (LossPred) [54]. As shown in Figure 3 (b), by filtering the genes whose uncertainty scores were higher than the median (denoted as a high-uncertainty group), we achieved lower MSE for both the thymus and PBMC datasets. Furthermore, to validate the relationship between the uncertainty levels of each gene as well as its statistics, we computed the variance across cells for genes in the high-uncertainty group and low-uncertainty group, and the average values are shown in Supplementary Figure 12. We found that genes with low uncertainty also had low variance on average, and thus our estimation of uncertainty is also supported by the features of tested datasets. Moreover, we explored the possibility of explaining the uncertainty scores from biological perspectives. Here we selected the remaining genes and ran Gene Ontology Enrichment Analysis (GOEA) [55, 56], recorded in Supplementary File 1. The adjusted p-value threshold of significance is set as 0.05. For both datasets, the gene sets with lower uncertainty always had more significantly enriched pathways (37 pathways from the low uncertainty group, 28 pathways from the high uncertainty group in PBMC; 148 pathways from the low uncertainty group, 112 pathways from the high uncertainty group in thymus.), which supports our findings that these genes play a more important role in biological activity. Moreover, genes with different uncertainty levels still have functional similarity reflected by the overlaps of pathways, including regulation of cellular metabolic process (GO:0031323), RNA metabolic process (GO:0016070), and others, while genes with lower uncertainty demonstrate more enrichment related to tissue-specific functions, supported by the tissue-specific pathway discovery shown in Supplementary Figure 13. For example, the discovered pathways related to myeloid development represents one of the most important thymus functions. We also analyzed the relationship between the coefficient of variation (CV) based on expression profiles for two gene sets, shown in Supplementary Figures 14 (a) and (b) for different datasets. We found that genes with lower uncertainty also have higher CV, which implies that genes with larger CV values may carry biological significance. In conclusion, our uncertainty estimation component can help us identify difficult components of expression prediction.

### Analyzing effects from cell types and individuals for expression prediction

Previous work suggests that the expressions and functions of the same gene vary across different cell types, which derived specific functions of cell types and tissues [57–59]. Therefore, we investigate if our model has different performances for different cell types. To conduct this experiment, we trained different models for different cell types to compute the MSE and correlation across genes and cells, and visualize our metrics in Figure 3 (c) for thymus and in Figure 3 (d) for PBMC. For the thymus dataset, our model achieved the best performance in predicting gene expressions for the ETP cells, and the worst for the Mgk cells, with no significant differences across different cell types. However, for the PBMC dataset, our model performed best for the CD8 T 2 cells, and worst for the Megakar cells, with significant differences between these two cell types. To explore factors associated with prediction performance, we calculated the PCCs between the number of cells for different cell types and the corresponding MSE metric for both datasets, and they both showed significant negative correlations (corr=-0.83, p-value=0.003 for the thymus dataset; and corr=-0.63, p-value=0.03 for the PBMC dataset). Therefore, UNICORN performs better in predicting gene expressions for major cell types, and increasing the number of sequenced cells can potentially improve the performance of UNICORN.

Although UNICORN showed promising performance in predicting gene expression for certain cell types, it might be more reliable to replace the sequences from the reference genome with sequences measured directly from the individuals. Therefore, we investigated whether UNICORN can perform individual-level gene expression predictions based on paired whole-genome sequencing (WGS) [60] data for individuals. Here, we collected all the scRNA-seq datasets as well as corresponding WGS data from the Genotype-Tissue Expression project (GTEx) v9 [61]. The database from GTEx v9 contains 16 individuals with their corresponding WGS and scRNA-seq datasets. First, we selected the gene expression profiles from the overlapped cell type (endothelial cell) across individuals and trained one model for each person. We also controlled the same training/validation/testing sets of genes across individuals and incorporated Enformer without fine-tuning as a baseline by selecting the output track of the endothelial cell as its output. The predicted values are evaluated based on the pseudo-bulk mode. In Supplementary Figure 16, we showed the performances of UNICORN in predicting the unseen genes for each person. UNICORN showed more similar expression patterns as well as consistency across individuals compared with observed expression levels, while Enformer failed to reflect the personal expression levels, shown in Figure 4 (a). Our findings also matched previous conclusions on the drawbacks of using Enformer for individualized gene expression prediction [62, 63]. To strengthen our conclusion, we also fine-tuned Enformer based on the selected sample 12BJ1 (which is the sample with the highest averaged correlation predicted by UNICORN) and visualize the comparison between fine-tuned Enformer and UNICORN in Supplementary Figure 15 (a). According to this figure, UNICORN also had lower MSE and higher celltype correlation. Examples of genes with good (high correlation) and poor predictions (low correlation) are shown in Supplementary Figures 17 (a) and (b). Therefore, the prediction of individualized gene expression at the cell-type level is practical, and UNICORN is a better model for learning the linkage between sequence and expression. Furthermore, we investigated the relationship between data quality and prediction performance to study the influence of technical noise on biological-related conclusions. We collected the cell type information of 16 different scRNA-seq datasets and computed the clustering scores by averaging Normalized Mutual Information (NMI) and Adjusted Rand Index (ARI) to evaluate the quality of given datasets, shown in Supplementary Figure 15 (b). According to this figure, we identify four samples (15EOM, 1MCC2, 1CAMS, and 1CAMR) whose clustering scores are lower than other samples’ scores. We then trained UNICORN based on the whole scRNA-seq dataset of each person and computed the correlations and MSE between observed and predicted gene expression profiles across individuals, shown in Supplementary Figures 15 (c) and (d). According to these two figures, we identified that samples with low correlations on average were also those with low data quality. These four samples also have a low cell-type-level correlation of the endothelial cell, shown in Figure 4 (a). After filtering these samples, we tested if the model trained by one sample could be used to predict the gene expression on endothelial cells of other samples, and the results are shown in Figure 4 (b) and Supplementary Figure 17 (c). According to these figures, the performance of different models is stable for a given testing sample, which also shows the potential of UNICORN in learning the shared genomic features across individuals. Therefore, UNICORN’s poor performances in certain samples are explained by their low data quality, and thus collecting scRNA-seq data with good cell-type stratification and paired WGS information should enhance the performance of UNICORN in predicting gene expression across different dimensions.

**Fig. 4:**
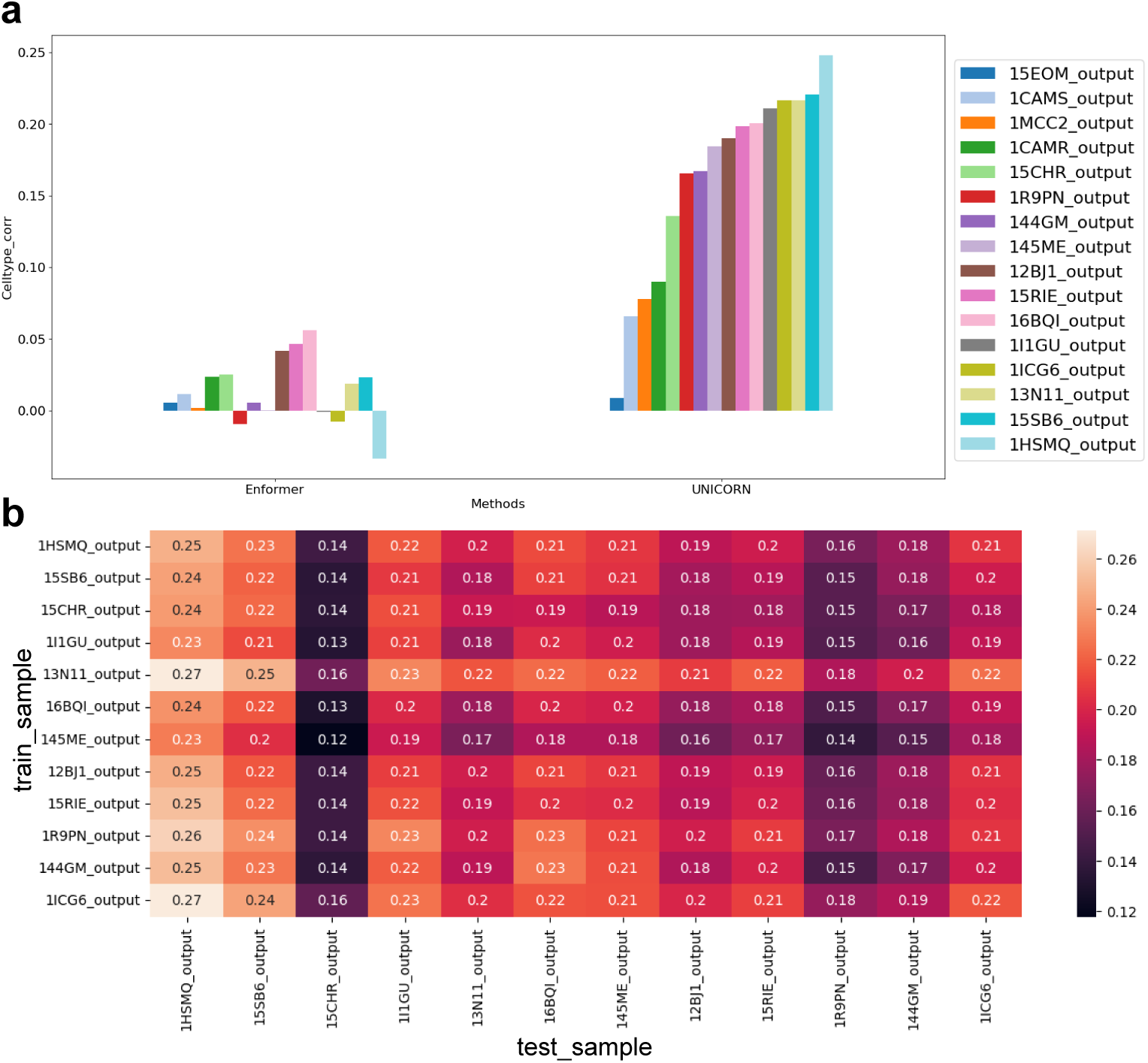
Results of gene expression predictions for individualized scRNA-seq datasets. The shared cell type is endothelial cell. (a) The results of cell-type correlations computed based on the given cell type across individuals for outputs from Enformer and UNICORN. Samples are ranked based on the cell-type correlation from UNICORN in ascending order. (b) The cell-type correlation of cross-individual prediction result based on UNICORN, after filtering the low-quality samples. The rows represent training samples and the columns represent the testing samples.

Since our current model accepts individualized genome information as inputs and individualized gene expression levels as outputs, our model can also be used to evaluate the effects of specific variants [64]. Our discussion can be found in Appendix F. Although UNICORN is a good model for predicting gene expression levels, this method still cannot explain the variant effects well due to the group-level effects of variants as well as the pre-training setting of Enformer.

### Predicting multi-omic phenotypes at cell-type resolution simultaneously

The capacity of UNICORN allows us to jointly predict multi-omic phenotypes from one cell type, which could help identify regulatory relationships between different biological features. In this section, we utilized one 10X Multi-omic dataset containing gene expression and ATAC peak profiles, and one CITE-seq dataset containing gene expression and surface protein expression profiles for experiments. For the 10X Multiomic dataset, the input of UNICORN came from the genomic sequence embeddings generated by Enformer. For the CITE-seq dataset, the input of UNICORN comes from the genomic sequence embeddings generated by Enformer and amino acid sequence from ESM2. We hypothesized that including multi-omic information can help the prediction of each modality, relative to the single-modality case. Therefore, we compared the prediction performances with only one modality with joint modalities, shown in Figure 5 (a) for the 10x Multi-omic dataset and Figure 5 (b) for the CITE-seq dataset. According to these two figures, having more modalities can help predict different modalities, especially for predicting the surface protein expression levels across cell types. We also tested various settings of embeddings and did not find significant differences, shown in Supplementary Figure 18 (a). Therefore, our current setting is capable of predicting the expression levels in the multi-omic setting.

**Fig. 5:**
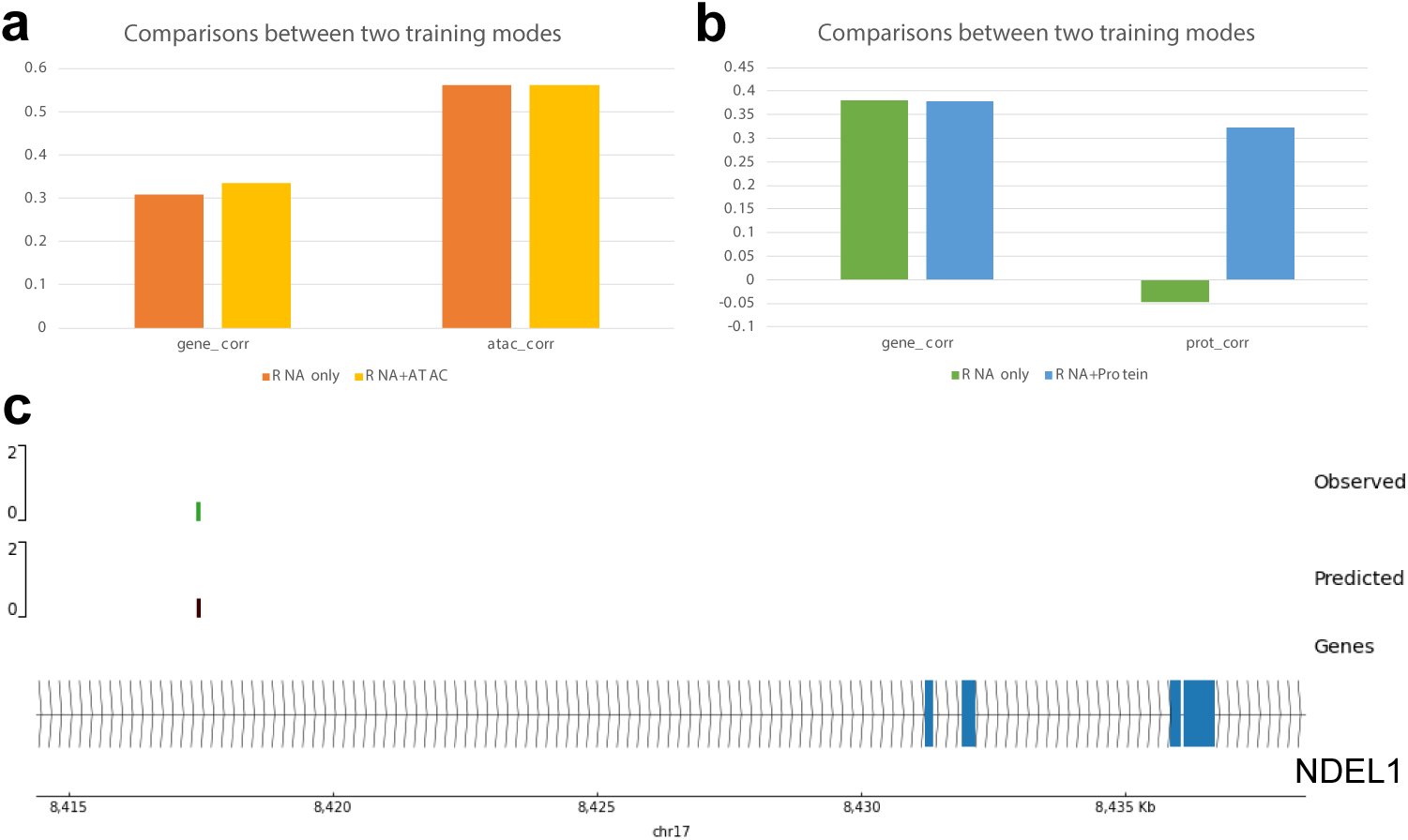
UNICORN demonstrates its ability in predicting multi-omic information at the cell-type level. Here RNA represents the gene expression profiles. The evaluation is performed based on the pseudo-bulk data. We reported the averaged correlation scores in (a) and (b). (a) The comparisons between RNA only mode or ATAC only mode with the joint training mode (RNA+ATAC). (B) The comparisons between RNA only mode or Protein only mode with the joint training mode (RNA+Protein). (c) Visualization of observed and predicted peak information for CD 14 Mono cells. We selected the peak with the highest correlation for analysis.

We also hypothesized that UNICORN can offer insights into selecting important features for downstream analysis. Here we selected the peak measured between 8417393 and 8418413 on chromosome 1, as its predicted value is highly correlated with the observed value (corr*>*0.99). We visualized the relationship between the predicted and observed values for this peak in Supplementary Figure 18 (b). Furthermore, this gene overlaps with NDEL1, as shown in Figure 5 (c). NDEL1 expressed in monocytes has proved to be able to regulate osteoclasts and bone homeostasis [65], and thus UNICORN can accurately predict the important peak associated with biological functions.

### Predicting disease effects and perturbation effects uncovers the biological variation

By performing prediction for gene expression levels under different conditions (including diseased/non-diseased and perturbed/non-perturbed), UNICORN can also help us understand the biological differences of cells and genes with respect to observed conditions. Following our previous design for analyzing the differences in prediction results across different cell types, we predicted the expression levels of testing genes based on three datasets from different conditions. The datasets Aorta [66] and Heart [67] are used to investigate the effects of diseases, and the dataset Openproblem [68] is used to investigate the effects of perturbations. The Aorta dataset has four disease states, including Ascending only, Ascending to descending, Ascending w/root, and Control. The Heart dataset has three disease states, including hypertrophic (HCM), dilated cardiomyopathy (DCM), and non-failing (NF). We utilized the default hyper-parameters to fine-tune different models for these datasets and compared the performances for the prediction results conditioned by disease states and perturbations. Our prediction results at the pseudo-bulk level are aggregated based on conditions and evaluated based on both cell-level and gene-level metrics.

First, we analyzed the differences in prediction results across different diseased and non-disease conditions. Figure 6 (a) shows the prediction results based on the Aorta dataset, and the labeled results such as “Subset: Control” represents that our trained model is only based on cells under the condition Control. According to this figure, UNICORN had the worst performance in predicting cells under the Ascending to descending condition, which is an intermediate state between the rest of the states. Such a finding implies that cells in the intermediate state might be hard to predict because cells under this condition are not stable. Figure 6 (b) shows the prediction results based on the Heart dataset, and the labeled results like “Subset: NF” represents that our trained model is only based on cells under the condition NF. According to this figure, UNICORN had the worst performance in predicting cells under the NF condition, which is unexpected. Such findings might be explained by the differences in data quality. Meanwhile, we assume that incorporating cells with different phenotypes can also enhance the overall prediction performance, and to validate if training with diseased cells can facilitate the prediction for normal cells or not, we include the prediction results labeled with “Subset: Control” and “Subset: NF” in our discussion. We found that for both datasets, including diseased cells can help predict gene expressions from normal cells and vice versa. Therefore, including heterogeneity in the training process can improve the performance of UNICORN. There is a similar conclusion from [69] related to cell-type transferring. Furthermore, UNICORN can preserve the expression levels of most differential expression genes (DEGs) under different disease states based on the testing split, validated by the strong correlation. For example, we observed accurate prediction results for genes RPL27A (corr=0.96), RPL28 (corr=0.76), and SON (corr=0.96) from the Aorta dataset, and gene NT5C2 (corr=0.99) for the Heart dataset. Notably, the variants in NT5C2 have been shown to have high risk in the association with heart diseases, and thus UNICORN also shows strength in capturing the correct expression levels for these disease-associated genes [70]. Therefore, UNICORN can help us analyze the differences of various conditions and suggests the importance of collecting cells from different states to build atlas-level databases.

**Fig. 6:**
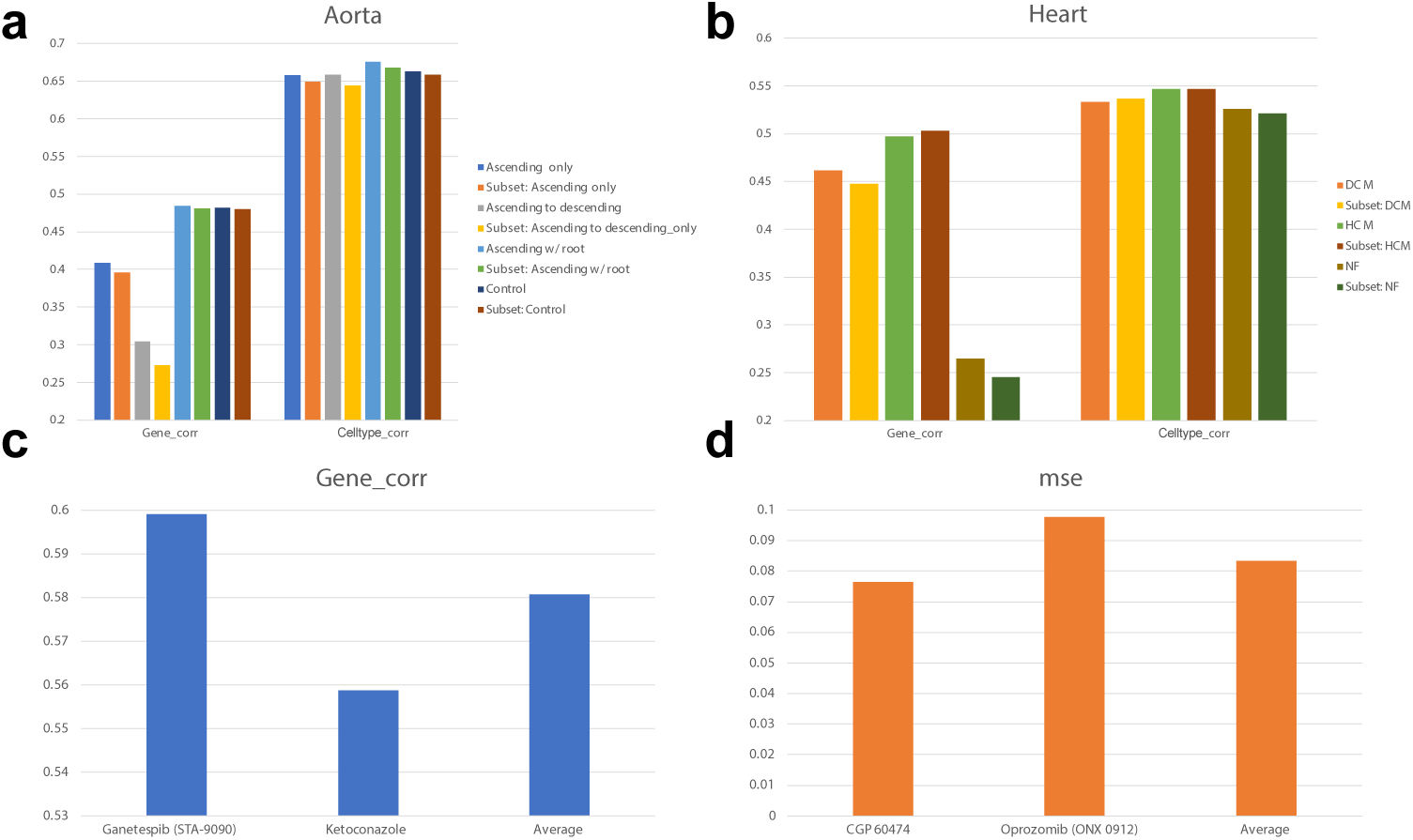
UNICORN uncovers the complexity of diseases and perturbations. (a) The correlation coefficients computed based on genes and cells across different conditions from the Aorta dataset. (b) The correlation coefficients computed based on genes and cells across different conditions from the Heart dataset. (c) Visualization of selected perturbation conditions by correlation coefficients computed based on genes. We only include perturbations with the highest and lowest correlation coefficients, as well as the averaged value across all perturbations. (d) Visualization of selected perturbation conditions by MSE computed based on genes. We only include perturbations with the highest and lowest MSE, as well as the averaged value across all perturbations.

We then investigated the performances of UNICORN in predicting gene expression levels under different perturbations. Figure 6 (c) shows the gene-level correlation results under three different cases, known as Ganetespib (STA-9090) (a perturbation with the highest corr score), Ketoconazole (a perturbation with the lowest corr score) and Average (the average corr score across all the perturbations). Such difference is also reflected in the results of the predicted DEG values under the perturbation cases, as the DEG SRSF7 of Ganetespib (STA-9090) had high correlation score (gene corr=0.70) and the DEG MX1 of Ketoconazole had low correlation score (gene corr=-0.77). Figure 6 (d) shows the MSE results under three different cases, known as CGP 60474 (a perturbation with the lowest MSE score), Oprozomib (ONX 0912) (a perturbation with the highest MSE score), and Average (the averaged MSE score across all the perturbations). The predicted value of DEG SRSF7 from CGP 60474 had a high correlation (gene corr=0.99), and DEG SNHG6 from Oprozomib (ONX 0912) also had a low correlation score (gene corr=-0.83). The ranks of the same perturbations under different metrics can also be different because of the existing intrinsic biases in evaluations. The value of metrics represents the difficulties of predicting these perturbations’ effects on gene expression levels based on the testing split, which can also be used to describe the heterogeneity of these perturbations. Therefore, the result of UNICORN can also offer a new direction to model the perturbation effects from a computational perspective.

Perturbation or disease effect can lead to the change of gene expression levels in both direction and magnitude. Therefore, it is also important to test if UNICORN can capture such difference in the predicted expression profiles across different conditions. We have provided the comparison of condition-level gene expression differences between predicted and observed levels. According to Supplementary Figures 19 (a) and (b), the difference computed based on predicted and observed data is similar and has a significant correlation for selected disease datasets. For the perturbation dataset, since we have collected over 100 perturbations, we display the correlation between the difference computed based on the predicted and observed levels of each perturbation in Supplementary Figure 19 (c). This figure also shows that all the correlations are positive, and the averaged PCC is also high (*>*0.3). Therefore, we can conclude that our model can correctly reflect differences in conditional gene expression profiles.

## 3 Discussion

Predicting gene expression from DNA sequence directly is a long-standing and challenging problem in biology. An effective model can help researchers better understand the relationship between regulatory elements and gene expressions at the cell-type or cell-state levels. In this work, we present a novel framework, called UNICORN, to address this research question with the help of prior biological information as well as an advanced machine learning framework. UNICORN utilizes pre-trained sequence-to-function models based on non-single-cell-level data as well as LLMs and is fine-tuned based on single-cell multi-omic datasets to perform predictions under different resolutions. Furthermore, UNICORN is capable of explaining the prediction process from different aspects. UNICORN not only can predict the multi-omic information but can also give individualized outputs based on DNA sequences from cohort studies and help in analyzing the perturbation effects based on diseased or perturbed data. Overall, UNICORN can improve prediction performances as well as gain biological insights on expression regulation.

Regarding the contributions of UNICORN for single-cell multi-omic data prediction, we demonstrated the feasibility of prediction of molecular expressions at the single-cell and cell-type resolutions, but the performance depends on data quality and requires attention to avoid overfitting. Although we demonstrated UNICORN’s better performances by running a comprehensive benchmarking analysis, it will always be helpful to explore novel pre-trained sequence-based models and possibly use them to enhance prediction performance. It is also interesting to note that pre-trained models focusing on gene expression prediction in other types of data performed better than other foundation models trained directly with DNA sequences in this task. Meanwhile, as expected, predicting gene expression at the cell-type level is easier than that at the single-cell level, especially for major cell types, which are further validated in our experiments for reducing the sparsity of datasets and increasing the number of cells used for training. By estimating the uncertainty levels of predicted genes, we can further offer more reliable prediction results. Therefore, UNICORN can help us uncover the heterogeneity of (gene) expression prediction at different resolutions.

Previous sequence-to-expression models utilize the reference genome as input data and thus unpaired expression profiles are used for training. Here we investigated the capacity of UNICORN in predicting individualized gene expression profiles with the help of paired WGS data from the GTEx cohort. The paired data also incorporate the variant information of each individual and thus it is a better resource than those models trained with unpaired information. Based on our analysis, the performance of UNICORN varies from person to person. Moreover, the pre-trained model (Enformer) used in this section is still from the reference genome and we lack such model pretrained with individual information, and thus it will be helpful to inherit the structure of UNICORN with an individual-aware sequence-based model for this task.

Inspired by our observations of the heterogeneity of performance across different cell types, we consider utilizing UNICORN to uncover disease effects and perturbation effects by analyzing the predicted gene expression across different conditions. Based on our experiments, including cells from more disease states can enhance the prediction performances under different conditions, even for the control case. Therefore, UNICORN can also be used to demonstrate the necessity of sequencing and collecting large-scale single-cell datasets under different disease states, which are valuable resources for model improvement. Regarding the analysis of perturbation effects, UNICORN showed different performances for different genes, which could be further used to investigate the strength of perturbation effects on cellular activity. Therefore, UNICORN can be used to explore the unique signals existing in different perturbations.

Despite the promising results from UNICORN, there is still a big gap between the DNA sequence and the corresponding expression levels. Firstly, the stochastic nature of gene expression regulation at the single-cell level coupled with high noise in expression measurements pose a major challenge to accurately construct the cell-type-specific gene expression, so there is a need to collect more high quality cells to improve prediction accuracy. Secondly, there are various approaches to pre-train a base model from sequence information for this task, and we are still looking for the best framework which can capture both the context of sequence information, the gene functional information, as well as the regulators for gene expression changes. Considering the inclusion of cohort-level data for enriching the corpus of the training set may be an approach. Moreover, UNICORN does not accurately capture the relative changes for certain diseases, which implies the necessity of dataset-specific fine-tuning with the base model. Finally, unraveling the contributions of variant effects at the single-cell or cell-type levels is still an important and challenging task, and the current framework may not fundamentally solve it due to the limitation of design. For example, we did not have models pre-trained with gene expression levels from individuals for different cell states, and we might need a better model architecture other than transformer or a novel loss function to capture the contribution of variants in controlling the expression levels or other related phenotypes. Therefore, exploring the contribution of a pre-training+fine-tuning framework for explaining such effects is still welcoming for further exploration.

Overall, UNICORN is a useful tool for unifying the expression prediction tasks from the sequence-based information and understanding the complex biology system created by the effects of genetic variation.

## 4 Methods

### Problem statement

Considering we have a single-cell dataset 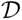 with modalities 1*…m*, we then can denote the single-cell dataset as 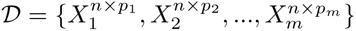. Each expression matrix *X* has the same number of sequenced cells *n* but a different number of biological features *p*_1_*…p_m_*. These expression matrices represent our prediction targets. Considering we also have biological feature embeddings for all modalities from pre-training models, which are denoted as 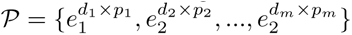. Our target is to fine-tune two models 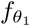 to predict the targeted expression levels and 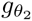 to estimate the uncertainty of prediction, where *θ*_1_ and *θ*_2_ represent two sets of parameters, that is:

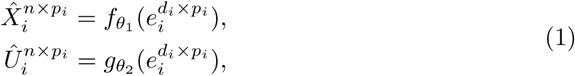

where *X̂* represents the predicted gene expression and *Û* represents the estimated uncertainty. Our equation utilizes the *i^th^* modality as an example. The output dimension of 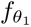 is equal to the number of cells, as we are predicting gene expression levels from sequence given the context of a specific dataset.

### Explanations of model architecture

There are two components of UNICORN. The first component is a pre-trained model to extract the embeddings from biological features as inputs. Using gene as an example, we utilize Enformer to generate DNA embeddings based on DNA sequences of the given genes and the GPT 3.5 embedding layer of the text description of these genes from NCBI to generate text embeddings. The source species of target datasets determine the genome of DNA sequences. We then combine DNA embeddings and text embeddings to finalize our dataset. We note that the generator of these two types of embeddings can be changed, so it should be updated with current state-of-the-art (SOTA) methods for each subproblem. Moreover, we use the same model to generate the embeddings of peaks for scATAC-seq data from DNA sequences and use ESM2 [71] to generate the embeddings of proteins for CITE-seq data from amino acid sequences. ESM2 was demonstrated as a top performer in protein-related tasks [72]. After having the group of embeddings as input, we assign separate predictors for each modality, and each predictor is constructed based on a two-layer MLP. We utilize ReLU [36] as the activation function and Softplus [36] as the output layer. The former design ensures that our model can learn non-linear relationships, and the latter design ensures that our model always has positive prediction results.

Meanwhile, for each predictor, we have the corresponding uncertainty estimator 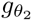. Our estimator and predictor have the same architectures but do not share weights because they work for different predictions. The method we used to estimate the uncertainty for prediction output is LossPred, which means we directly predict the MSE of the given example during the training process. This uncertainty estimation component will combine with the default loss function when we are interested in exploring the uncertainty quantification of different features. Such a method, supported by [53], outperforms other methods in their evaluation results across various datasets. Our current design is more efficient than full-parameter fine-tuning based on Enformer or other large-scale prediction models for genomic data, and we also demonstrate the superiority of our current design in the Results section.

### Model Explainability

We utilize function-based explainability to understand the factors that affect the performance of predictions and select critical genes. The idea of function-based explainability is designed to understand the contribution of sample-level information for making decisions based on deep-learning models, which has been widely discussed in several method-driven papers or systemic review papers [73–75]. This idea is also critical in biology research for us to understand hidden mechanisms in a transparent approach [76]. In our experiment, each sample is one gene so our target is to understand the role of gene-level signals in making the prediction. To achieve this target, we have designed several experiments for exploration. First, we consider processing the training data with different pre-processing methods or aggregating methods to investigate the progress of making more accurate predictions. This approach can be interpreted as an approach to explain the effect of distribution-level assumption for gene expression measurement. Second, we perform various data ablation studies to investigate the introduction of sparsity and cell-type proportion for model prediction. This approach can be interpreted as an approach to explain the effect of the modeling assumption for gene expression measurement. Third, we utilize the estimated uncertainty of biological features to select genes with low uncertainty and conduct biological analysis as well as training dynamics analysis. Our uncertainty estimation approach has a comprehensive theoretical foundation supported by experiments with simulation and real transcriptomic data. This approach allows us to provide a computational-level explanation for each gene and uncovers the biological signals from different pathway-associated genes selected by UNICORN.

### Details of model training

We first discuss our choices of loss functions. We utilize different loss functions for single-modality prediction and multi-modality prediction. For the single-modality prediction, we combine mutual Pearson correlation (MPC), Poisson negative log-likelihood loss (PNLL), and mean squared error (MSE) losses together to finalize our loss functions. Considering we have prediction *Ŷ_k_* and observed values *Y_k_* for modality k, our loss function is defined as:

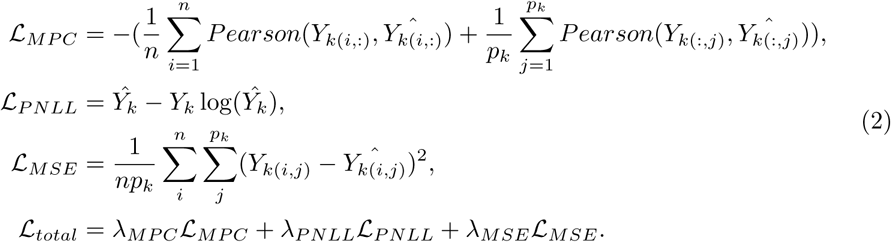

Here, the function *Pearson*(*a, b*) computes the Pearson correlation coefficients between two items, and our method is based on cosine similarity, and thus, it is differentiable. the notation (*i, j*) represents the *i^th^* cell and *j^th^* feature in the predicted matrix. The weights of these loss components, denoted by *λ_X_*, are the same after tuning with different options. To elaborate on the necessity of including all three loss components to formalize our final loss function, we perform ablation tests shown in Supplementary Figures 20 (a)-(c) under different metrics.

For the foundation support of our loss function design, we referred from the theory of empirical risk [77] and multi-task learning [78] to provide further support. Our loss function considers three different tasks and thus it covers multiple objectives when performing optimization. To predict high-dimensional biological data, relative patterns are more robust and biologically meaningful. Therefore, maximizing the correlation between predicted values and observed values can better align model prediction with underlying biological structure. Furthermore, Pearson correlation is scale-invariant, so it can also help our method learn invariant representations that preserve structure rather than magnitude. Based on the observation of data sparsity and distribution, we introduce Poisson Negative Log-Likelihood loss (PNLL) for modeling data sparsity explicitly and model gene expression profiles. Finally, we include mean squared error loss to capture the magnitude-level similarity between predicted value and observed value, which further captures the value-level fidelity globally. Therefore, in our design, different components act as regularizers, guiding the model to learn representations that are simultaneously expressive, interpretable, and robust to noise and data scale. For multi-modal prediction, instead of applying separate prediction heads for each modality, we predict the multi-modal information jointly for a single cell or a single cell type. We design the loss functions for multi-modal prediction based on a distribution-aware strategy. Since scATAC-seq is very sparse, we use the same loss function designed for regressing gene expression profiles. Moreover, after normalization, the proteomic part of CITE-seq data is dense, so we only keep the PCC loss. This principle of loss design is also supported by our ablation studies for multi-omic expression prediction.

To elaborate on the necessity of including distribution-aware loss function design to formalize our final loss function, we perform ablation tests shown in Supplementary Figures 20 (d)-(f) under different metrics. Here we found that including all three components is helpful for predicting the combination of scRNA-seq and scATAC-seq, while the prediction for CITE-seq data cannot be benefited by loss function modification. This observation aligns well with our discussion of the distribution-aware loss function design strategy. We also demonstrate the contributions of multi-modal prediction by comparing it with prediction based on a single modality.

We also discuss the hyper-parameter selection for UNICORN. Most of the hyper-parameters are referred from seq2cells and tuned based on model’s performance computed on the validation dataset. Details of our discussion can be found in Appendix D.

Regarding the motivation and foundation of uncertainty estimation based on LossPred, we have provided theoretical support as well as validation based on simulation data in Appendix G.

### Methods for performing biological analysis

To map the biological features (including genes and peaks) with their relevant sequence window, we utilize the construction function from [79] to retrieve the location of such features in the chromosome as well as the starting site and the ending site.

To retrieve the functions of genes, we downloaded the gene functional information of labels from [80]. These genes are classified into protein-encoding genes, non-coding RNA, etc. Details of gene functional list can be found in Supplementary Figure 24 (a). To infer the functions of gene sets we are interested in, we utilize Gene Ontology Enrichment Analysis (GOEA) [55, 56] to analyze genes of different sets. GOEA is based on Fisher’s exact test to compare the difference between the selected gene sets and all background genes, with outputs including significance level and overlap ratios.

### Information of metrics

To evaluate different models for expression predictions, we consider two metrics for two dimensions. The metrics include Pearson correlation coefficients and mean squared errors. These metrics are widely used in evaluating tools for gene expression prediction [11, 21, 24, 27, 81–83]. Since the sequencing data do not contain auto-correlated patterns, correlation can capture the similarity of tendency between predicted and observed expression levels, while MSE can capture the absolute similarity between predicted and observed expression levels. Therefore, the combination of metrics we choose provides a comprehensive measure of the difference between predicted and observed profiles. Our metrics are computed using the scikit-learn package [84] and based on testing genes. Our dimensions include gene-level evaluation and cell-level evaluation. The former setting represents that we compute the correlation coefficients across genes for each cell/cell type/phenotype, and the latter setting represents that we compute the correlation coefficients across cells/cell types/phenotypes for each gene. We report the mean, median, and quartiles as statistics shown in boxplots. The value of correlation is in (−1, 1), and a higher value represents better model performances. The value of MSE is in (0, ∞), and a lower value represents better model performances.

To evaluate different gene embeddings as well as cells for clustering, we consider Normalized Mutual Information (NMI), Adjusted Rand Index (ARI), and Averaged silhouette Width as metrics. All of them are in [0, 1], and higher values represent better performances.

### Information of baselines

We consider different baselines for different tasks. For the task of gene expression prediction, we include full-feature-based methods including seq2cells [11], Enformer [21], Borzoi [24], Decima [22], GET [44], and variations of UNICORN as baselines. seq2cells is a model based on the sequence embeddings from pre-trained Enformer to predict gene expression levels at the single-cell resolution. Enformer and Borzoi are designed to predict gene expression from the corresponding sequence information, which allows us to fine-tune the full model to predict gene expression levels in single-cell resolution. Decima is pre-trained with the pseudo-bulk gene expression of large-scale single-cell transcriptomic data, and it can predict the gene expression of specific cell types under specific tissues. GET is pretrained with (sc)ATAC-seq and it can predict gene expressions by fine-tuning this model with cell-type-specific information from scATAC-seq and scRNA-seq data. Our pipeline for fine-tuning Enformer and Borzoi follows the default design of gReLU. The variations of UNICORN come from different types of inputs. We also consider using embeddings from text descriptions of gene functions (genept) [39], embeddings from the DNA foundation model (HyenaDNA [2], DNABERT [29], NT [37]), and embeddings from gene-gene interaction (gene2vec) [38], as inputs. We also consider feature-selection-based methods, including SCOVER [41], DanQ [42], and LassoNet [43]. SCOVER is a convolutional neural network (CNN) based methods with motif-scoring design for expression prediction. DanQ integrates both CNN and recurrent neural network (RNN) to model the functional information of genomic regions. LassoNet is a feature-selection-based method by penalizing the sparsity of features during the training process.

For the task of model explainability, we consider scVI [50] and DeepImpute [51] as baselines for imputing single-cell datasets. scVI is a model based on a variational autoencoder to learn the cell embeddings and the distribution of gene expression levels in single-cell data. DeepImpute is a deep-learning-based method to impute missing gene expression in single-cell data by denoising auto-encoder.

For the task of multi-omic data prediction, we consider UNICORN with a shared predictor as the baseline model.

### Data preprocessing

For the gene expression prediction task, we performed the log-normalized approach for scRNA-seq data based on scanpy [85]. We compared the performances of UNICORN between the raw mode and normalized mode in Appendix E. For the multi-omic data prediction task, we used the raw count data from scATACseq and scaled gene expression data from scRNA-seq. Such a setting is relevant to our design of loss functions.

## 5 Data Availability

We do not generate new data in this research and all data used in this manuscript are publicly available without restricted access. The thymus dataset is available under accession code [https://zenodo.org/records/8314644]. The PBMC dataset is available under access code [https://zenodo.org/records/8314644]. The thymus atlas dataset is available under access code [https://zenodo.org/records/8314644]. The Aorta dataset is available under access code [https://www.ncbi.nlm.nih.gov/geo/query/acc.cgi?acc=GSE155468]. The Heart dataset is available under access code [https://singlecell.broadinstitute.org/singlecell/study/SCP1303/single-nuclei-profiling-of-human-dilated-and-hypertrophic-cardiomyopathy]. The Openproblems dataset is available under access code [https://www.kaggle.com/competitions/open-problems-single-cell-perturbations/data]. The Onek1k dataset is available under access code [https://onek1k.org/]. For the datasets from GTEx, users need to submit application for access under access code [https://www.gtexportal.org/home/]. For the datasets from ROSMAP, users need to submit application for access under access code [https://adknowledgeportal.synapse.org/Explore/Studies/DetailsPage/StudyDetails?Study=syn3219045]. We include the links to access our experimental datasets in Supplementary File 2. Source Data are provided with this paper.

## 6 Code Availability

To run UNICORN, we relied on Yale High-performance Computing Center (YCRC) and utilized one NVIDIA A40 GPU with up to 40 GB RAM. The upper bound of running time for analysis is 24 hours.

The codes of UNICORN can be found in https://github.com/HelloWorldLTY/UNICORN. The license is MIT license.

## 8 Acknowledgments

We appreciate the comments, feedback, and suggestions from Dr. Gokcen Eraslan for model training.

## 9 Author contributions

T.L. and Y.L. designed this study. T.L. and T.H. designed the model. T.L., T.H., and L.W. ran all the experiments. T.L., T.H., L.W., Y.L., R.Y., and H.Z. wrote the manuscript. R.Y. and H.Z. provided the computation resources. H.Z. supervised this work.

## A How to describe the effect of input sequence length to gene expression prediction performances?

In this section, we intend to investigate the contributions of increasing window size of sequence length for gene expression prediction. Previous methods claimed that sequence-to-function models should capture long context to enhance more performances, so we hypothesized that such a conclusion should also fit this problem. To validate our assumption without being biased by the design of pre-trained models, we paired the sequence information centered by TSS but with different context lengths with single-cell gene expression profiles, and trained a linear regression model to evaluate our results. According to Supplementary Figure 22, the MSE increased as we increased the length of the context window, while the other two correlation-based scores also increased. Therefore, expanding the context window helps in learning the correlation between prediction outputs and observed gene expression levels, which might not help in learning the specific magnitude of gene expression levels.

## B Can we predict gene expression levels in atlas-level single-cell datasets?

In this section, we demonstrate that UNICORN is capable of predicting gene expression levels in the atlas dataset from human thymus (thymus atlas). Our dataset contains 255,901 cells and 18,204 genes. We used the same training setting and generated the gene embeddings using PCA from the observed gene expression and predicted gene expression for training genes shown in Supplementary Figure 23 (a) and testing genes shown in Supplementary Figure 23 (b). This figure shows that our predicted gene embeddings overlap with observed gene embeddings well for most of genes. Meanwhile, for the genes whose embeddings are not aligned well under the observed case and the predicted case, the observed gene embeddings show clustering patterns which are less informative under both training and testing cases. Therefore, the unaligned patterns of gene embeddings might come from the quality of data or genes rather than the capacity of UNICORN. Moreover, we also visualize the distribution of cells based on the predicted gene expression levels in Supplementary Figure 23 (c), which preserves the separation of different cell types in a good manner. Therefore, our method also supports atlas-level prediction without facing memory issues.

## C Can the functional annotations of genes fully explain our prediction performances?

In this section, we collect functional labels (e.g., protein-encoding and lincRNA) of genes and use different gene embeddings to quantify them. Our evaluation framework is inspired by the difference of their functionalities and the clustering performance based on gene embeddings can reflect the ability to represent gene functions. Understanding the relationship between gene functions and gene embeddings may help us explore the contributions of gene embeddings on expression prediction. In total we consider gene embeddings from GenePT, Gene2vec, Enformer, Enformer+GenePT (Combine), HyenaDNA and DNABERT and benchmark their performances on gene clustering. The UMAP plots of different gene embeddings are shown in Supplementary Figure 24 (a), with all of them being similar because the dominant genes are protein-encoding genes for scRNA-seq datasets. Furthermore, we compute three different metrics for evaluating clustering performances, including Normalized Mutual Information (NMI), Adjusted Rand Index (ARI) and Averaged silhouette Width, and average them to have the final score. We rank these method based on this score, shown in Supplementary Figure 24 (b). This figure shows that the best model for gene clustering is GenePT, followed by Gene2vec and Enformer. This rank is not highly correlated to the method’s rank for gene expression prediction. Therefore, our discovery might explain why direct inference based on gene embeddings is difficult for expression prediction, and further supports the utilization of our adapter design.

We also explore the relationship between gene-level similarity and prediction performances. Since embeddings from different foundation models have different dimensions, the gene-level similarity computed based on cosine similarity can unify the representations across different embeddings and expression profiles to explore if the embeddings can correctly capture the gene-gene similarity reflected in the expression level. We first make a comparison between the gene-similarity computed by Enformer and GenePT (LLM), shown in Supplementary Figures 25 (a) and (b). Based on these figures, we found that the gene-level similarity computed based on embeddings from Enformer and LLM is not exactly the same, which implies that they may capture different aspects of functional information. These embeddings also align well with the gene-level similarity from expression profiles, and thus LLM can provide unique contributions when building the model for predicting gene expression.

Moreover, we also make a similar comparison across different DNA Foundation models and sequence-to-function models, shown in Supplementary Figures 26 (a) and (b). These figures show that the similarity captured by the Enformer-based approach always aligns with the similarity captured by gene expression levels better than other models, and thus Enformer is still the best choice if we intend to predict gene expression levels directly from DNA sequence embeddings.

## D Do we need to tune hyper-parameters for prediction?

In this section, we discuss the hyper-parameter tuning process for UNICORN based on the validation dataset. Our best epoch is determined by early-stopping with the help of a validation dataset. Therefore, we only considered the effects of 1. Learning Rate (LR), and 2. Dropout Rate (DR) and 3. batch size. Our results are summarized in Supplementary Figure 27 (a) for LR, Supplementary Figure 27 (b) for DR and Supplementary Figure 27 (c) for batch size. We found that LR has larger effects for model performances and we finally set LR as 1e-4. Furthermore, DR does not affect model performances obviously and we finally set DR as 0.4, and adjusting the weight or batch size does not affect prediction results too much. Such setting is kept the same across all of the experiments in this manuscript.

We also explore the contribution of adjusting loss function weights for the three different components. The analysis based on gene expression prediction is shown in Supplementary Figures 27 (d)-(f). According to these figures, changing the weight does not obviously affect the prediction performance (less than 0.001). Therefore, we keep the same weight for all three components.

## E Do we need normalized single-cell data for prediction?

The normalization approach for pre-processing single-cell data has been well-discussed [86, 87], and the raw single-cell data are biased to the sequence depth [88]. To validate the necessity of utilization log-norm data for prediction, we utilize the PBMC dataset as an example and perform prediction for raw count data, log-normalized data, and sctransform [89] data as targets. Our results are shown in Supplementary Figures 28 (a)-(d). According to these figures, prediction outcomes based on log-normalized data have lower variance and MSE, and its correlation coefficients based on cells are also higher than the raw data mode, while comparable with the sctransform mode. Prediction results based on sctransform data also have higher MSE than the log-normalized mode result, which suggest that the log-normalized approach is more suitable in this case. Therefore, the utilization of log-normalized data can help us generate more stable and reliable outputs.

## F Can we use sequence-to-function models to validate variant effects in cell-type levels?

Here, we collected the fine-mapping variants from CausalDB [90] and utilized the effective alleles and non-effective alleles to generate different groups of prediction results for cells, which correspond to reference output and variant output. We ranked the p-values of different fine-mapping variants in ascending and selected the first 100 variants as well as the last 100 variants to validate the contributions of variants for gene expression levels. Our results are shown in Supplementary Figures 29 (a) and (b). Based on these two figures, we found that the correlation of gene expression levels between reference outputs and variant outputs is consistently high (≥0.99) (while the MSE is consistently low (∼1e-7)) for both groups. We further computed the Rank-sum test statistics for both the correlation coefficients and mean squared errors between reference outputs and variant outputs using these two groups, and both of them did not show significance (p-value=0.33 for correlation coefficients, and p-value=0.50 for mean squared errors). Therefore, we did not consider further analyses for the effect of a single variant on gene expression prediction. One possible reason to explain the performance is that the variants in the genome always work as a group [11, 91, 92]. Another reason is, the model Enformer was pre-trained based on the reference genome rather than the individual genome, which could also introduce a potential bias for individualized prediction.

## G Theoretical Discussions on Uncertainty Quantification

Regarding the foundation of our uncertainty quantification approach, the methodology employed in the manuscript has been extensively discussed in prior studies, including [93] and [94]. [93] provides a comprehensive review on the uncertainty quantification, and proposes directly estimating the uncertainty by learning a secondary predictor. Similar strategies have also been explored in recent studies by [95] and [96].

Specifically, consider *X* ∈ ℝ*^p^* and *Y* ∈ ℝ, where their relationship can be expressed as:

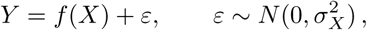

where *σ_X_* denotes the heterogeneous errors that might depend on the features *X* = (*X*_1_*,…, X_p_*). For example, 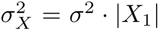. The *primary* predictor *f̂* learns the relationship between *X* and *Y*, and the *secondary* predictor *ĝ* learns the uncertainty *Y* − *f̂*(*X*), as a function of *X*.

Note that the true uncertainty is characterized by the (unknown) *σ_i_*, whereas we only observe the empirical error 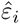. Given the powerful representation capabilities of neural networks, the predictor *f̂* often closely approximates *f*. Specifically, if *f̂* = *f*, then the empirical uncertainty simplifies to

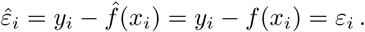

However, *ε_i_* represents just one realization from the distribution governed by *σ_i_*. While a larger *σ_i_* typically results in larger observed values of *ε_i_*, a small *σ_i_* can also occasionally produce large values of *ε_i_*. Therefore, it is essential to examine the consistency between *ε_i_* and *σ_i_*. Particularly, we investigate whether our procedure proposed which divides the *empirical* uncertainty into two groups can effectively reflect the *true* high and low uncertainty groups. Proposition 1 addresses this question by establishing the asymptotic accuracy of assigning empirical uncertainties to their correct underlying uncertainty groups.

### Proposition 1.

*Let* 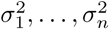 *be i.i.d. from a distribution with density function h*(*t*) *supported on* [*a, b*] ⊂ [0, ∞)*, a < b. Suppose* 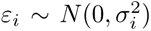*, i* = 1*,…, n. Let* 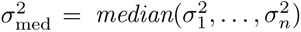 *and* 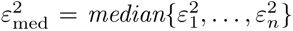*. Denote* 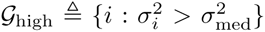*and* 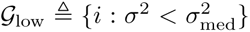 *as the*true *high uncertainty and low uncertainty groups, respectively. Similarly, denote* 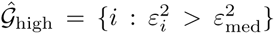, 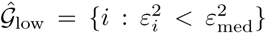 *as the* empirical *high uncertainty group and low uncertainty group, respectively. Then for another σ*^2^ ∼ *h and ε* ∼ *N* (0*, σ*^2^)*, we have, when n goes to infinity, the accuracy of assigning ε to the true low group is given by*

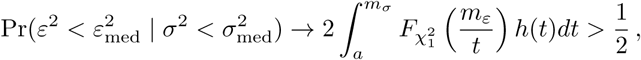

*where* 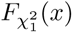 *is the CDF of* 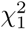 *distribution, m_ε_ satisfies*

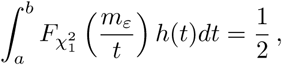

*and m_σ_ satisfies*

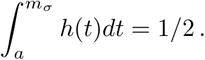

Proposition 1 shows that the probability of correctly assigning empirical uncertainties to their true groups is strictly greater than random guessing (i.e., greater than 1/2). To illustrate the results, we consider *h* as the distribution of *U ^k^*, i.e., the density function 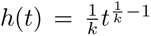, *t* ∈ [0, 1]. We then compute the theoretical asymptotic probability and verify this probability numerically using Monte Carlo approximations.

Supplementary Figure 30 validates the theoretical probabilities by the Monte Carlo experiments, clearly demonstrating that larger discrepancies between high- and low-uncertainty groups (e.g., the discrepancy in *U* ^2^ is larger than the one in *U*) result in higher classification accuracy. Furthermore, we conducted additional experiments in the real gene-expression scenarios studied in our manuscript. Here we first fit a model *f̂*(*X*) based on the thymus dataset. We further adding different noise level to simulate different outcomes *Y* from two cases with *ɛ* ∼ *N* (0*, σ*^2^). The *σ* ranges from 0.01 to 1.0 uniformly in the first case, which corresponds the condition *U* ^1^. Moreover, we take the square of *σ* in the first case and generate the variance used in the second case, which corresponds the condition *U* ^2^. We then measure the PCC and SCC between predicted uncertainty and real variance level, as well as the accuracy of capturing the correct noise-uncertainty relationship with median as threshold, to evaluate the performance of uncertainty estimation. To reduce the effect of randomness, we repeated our experiments by varying five different random seeds and report the group-level result. According to Supplementary Figure 31, our experiment results (ACC) match the theoretical conclusions (probability) well, as the averaged accuracy from experiments is close to the theoretical results from the two conditions, and the averaged accuracy in the second case is larger than the average accuracy in the first case.

The resulting accuracies closely align with our theoretical results, providing strong evidence that the proposed approach effectively identifies and excludes low-quality samples.

### Proof.

Consider the asymptotic distribution of 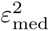 and 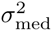.

By the central limit theorem for medians, we have

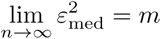

where *m* satisfies

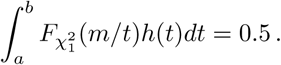

The empirical median of 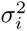 converges to the median of the distribution of *H*, where *H* ∼ *h*.

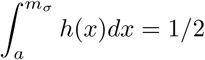

Particularly, if *H* = *U* [0, 1], then *m_σ_* = 1*/*2. Then the accuracy is given by

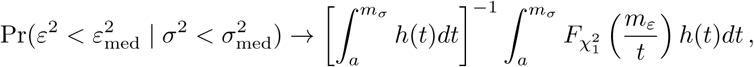

where *m_ε_*satisfies

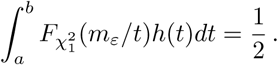

For the lower bound 1*/*2, note that

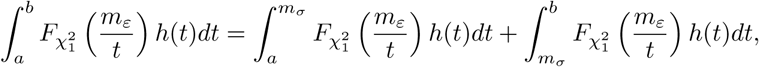

since 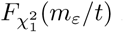 is a strictly decreasing function of *t* and

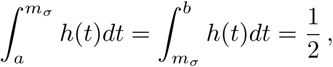

it follows that

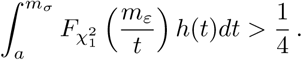

## H Supplementary figures

**Supplementary Fig. 1:**
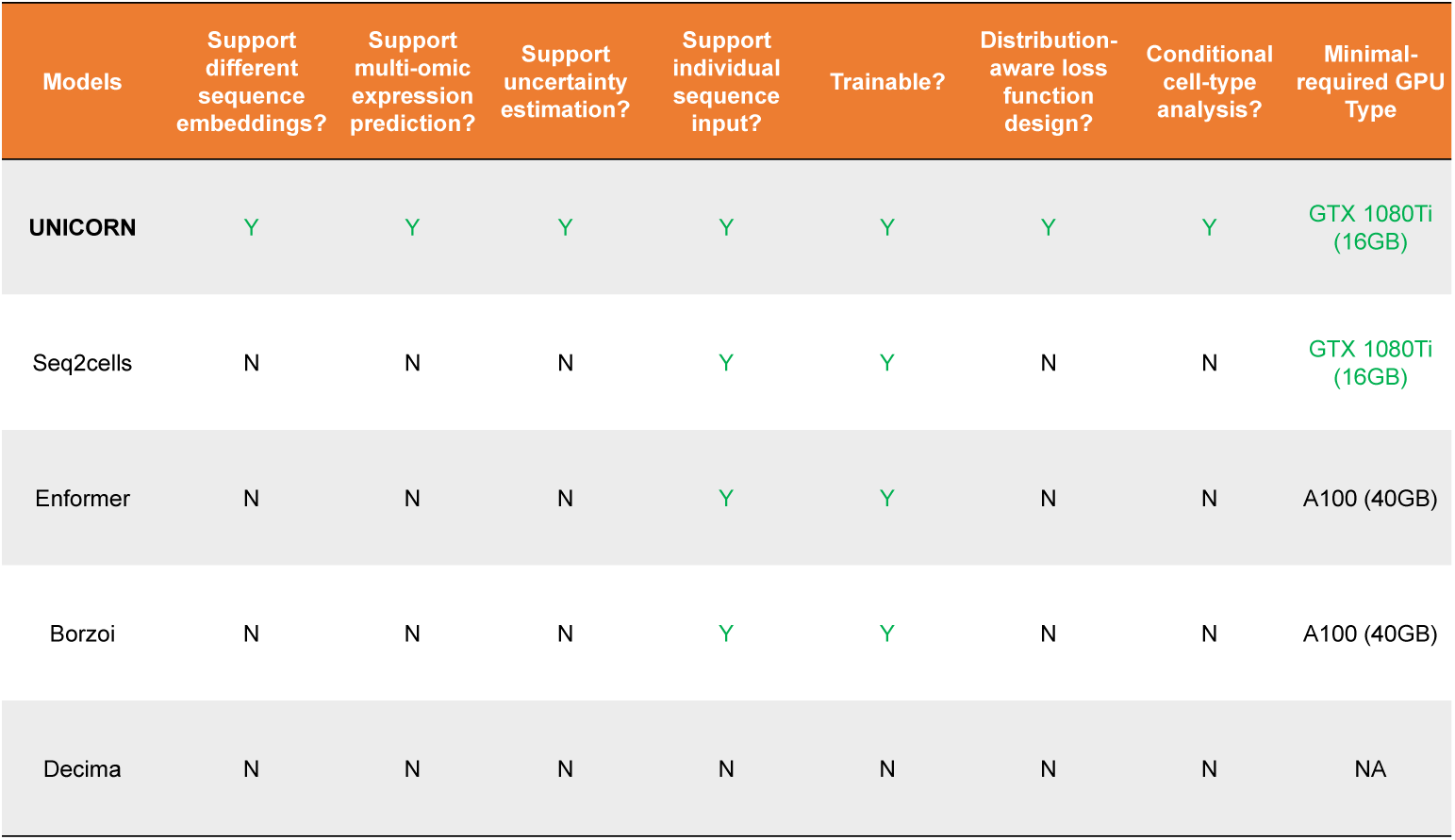
Comparison of different models based on their advantages in design and implementation.

**Supplementary Fig. 2:**
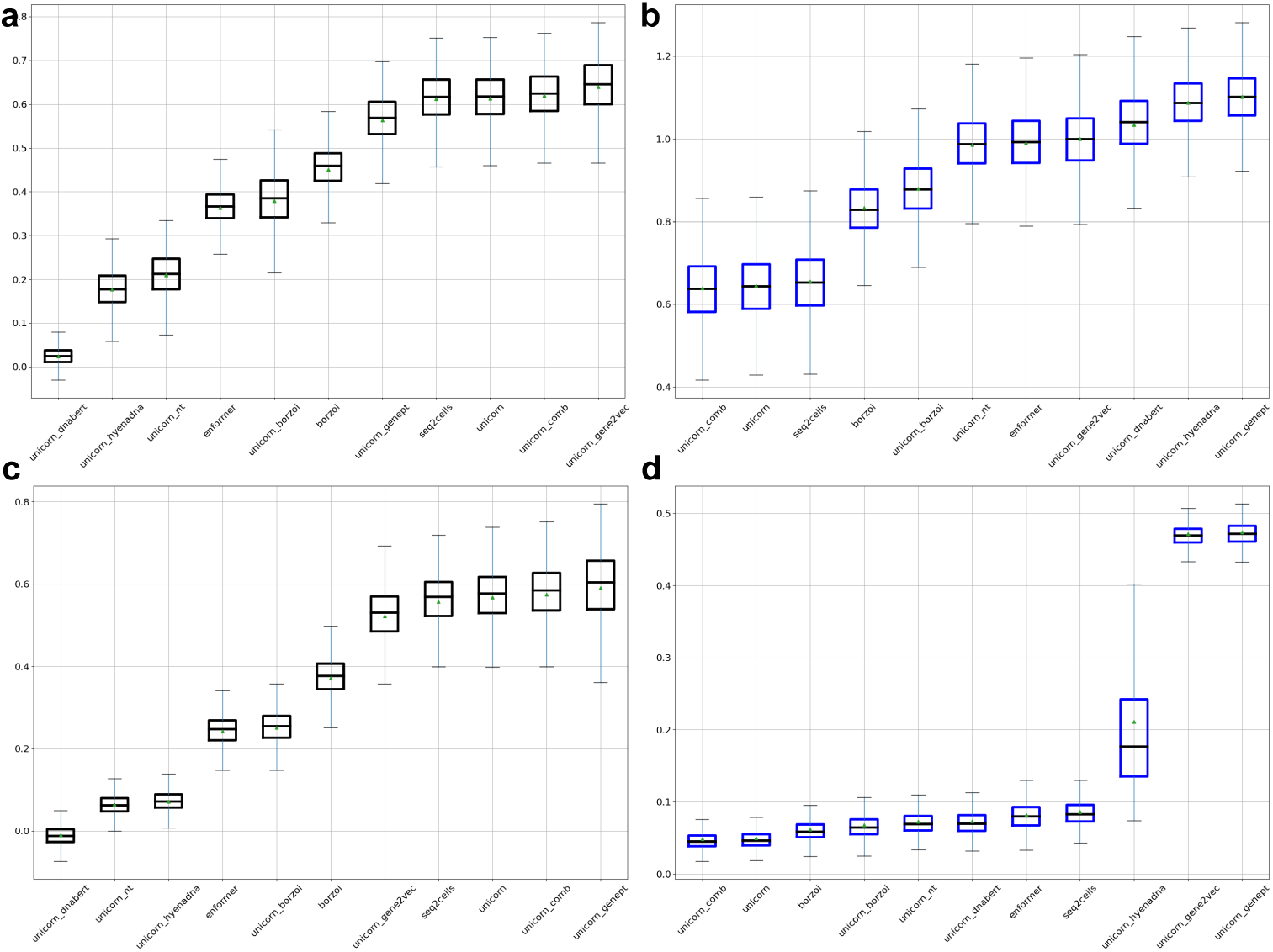
Comprehensive evaluations for gene expression predictions from sequences. The triangle shape represents the mean value, and the black dashed line represents the median value. We report the results based on box plots. (a) Results of cell-level correlations across different methods based on the thymus dataset. (b) Results of MSE across different methods based on the thymus dataset. (c) Results of cell-level correlations across different methods based on the PBMC dataset. (d) Results of MSE across different methods based on the PBMC dataset.

**Supplementary Fig. 3:**
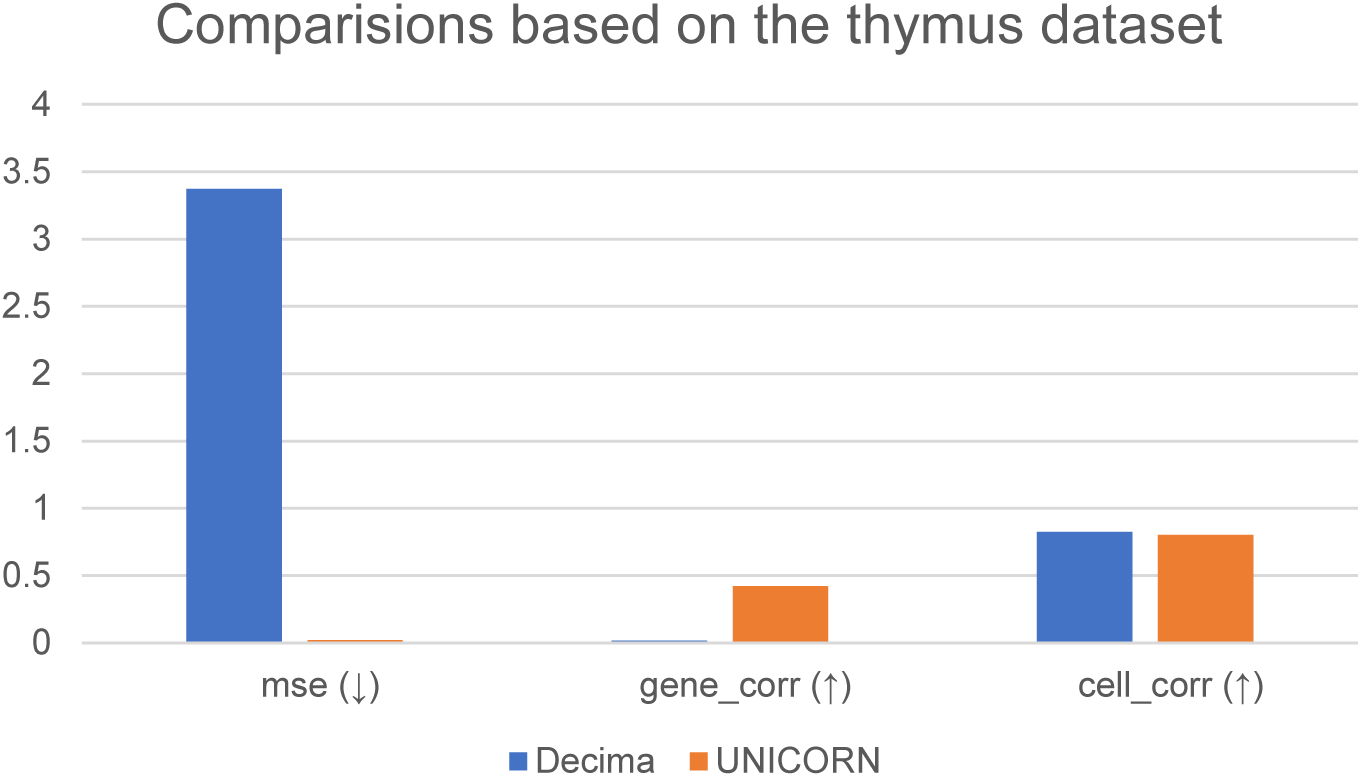
Comparison between UNICORN and Decima based on the thymus dataset. To make a fair benchmark, we select the overlapped cell types and tissue from Decima’s prediction results.

**Supplementary Fig. 4:**
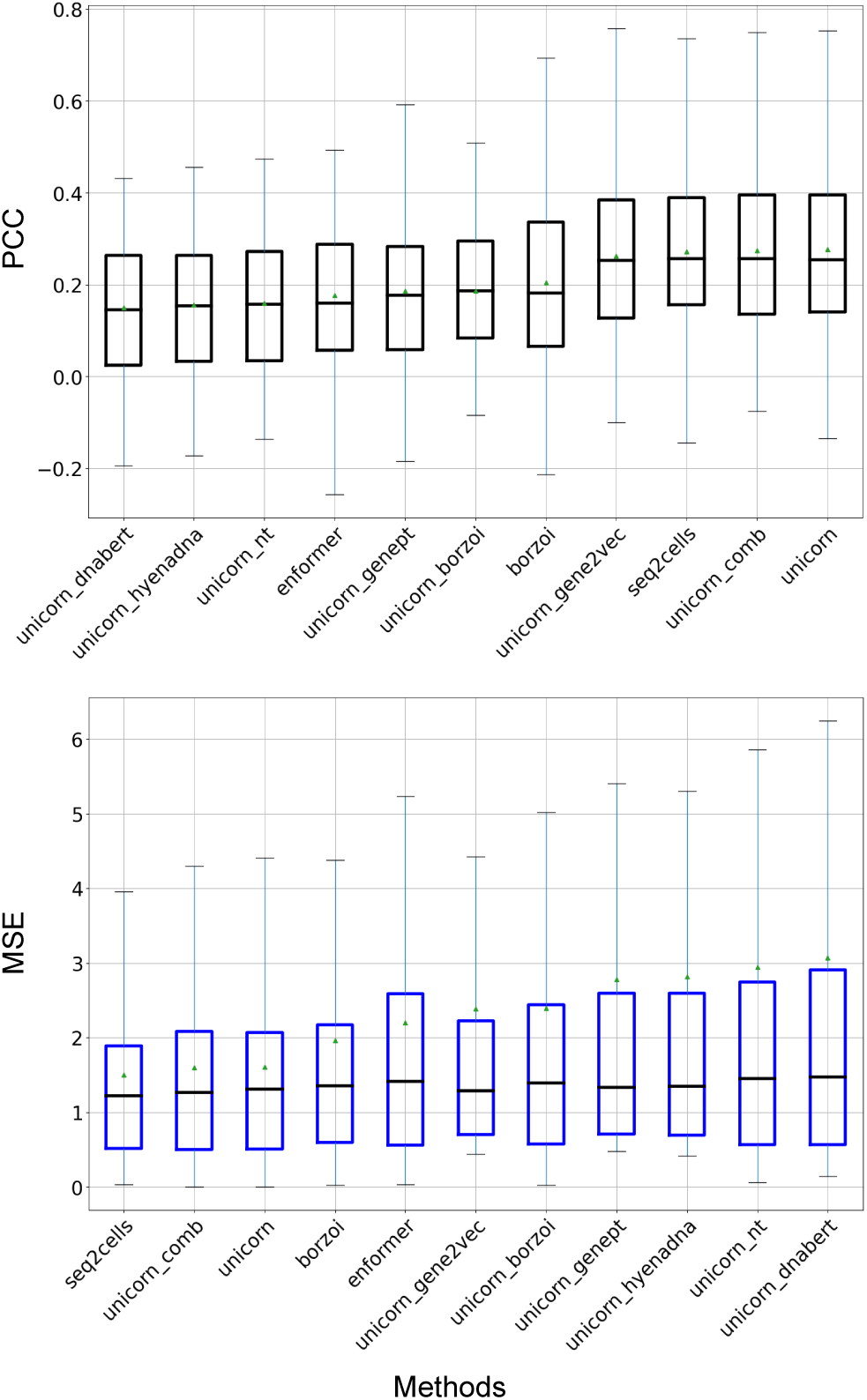
Comparison between UNICORN and other methods for predicting marker genes’ expression levels from different cell types. Here we present the results from top 50 marker genes. (a) Results of updated gene-level correlations across different methods based on the thymus dataset. (b) Results of updated MSE across different methods based on the thymus dataset.

**Supplementary Fig. 5:**
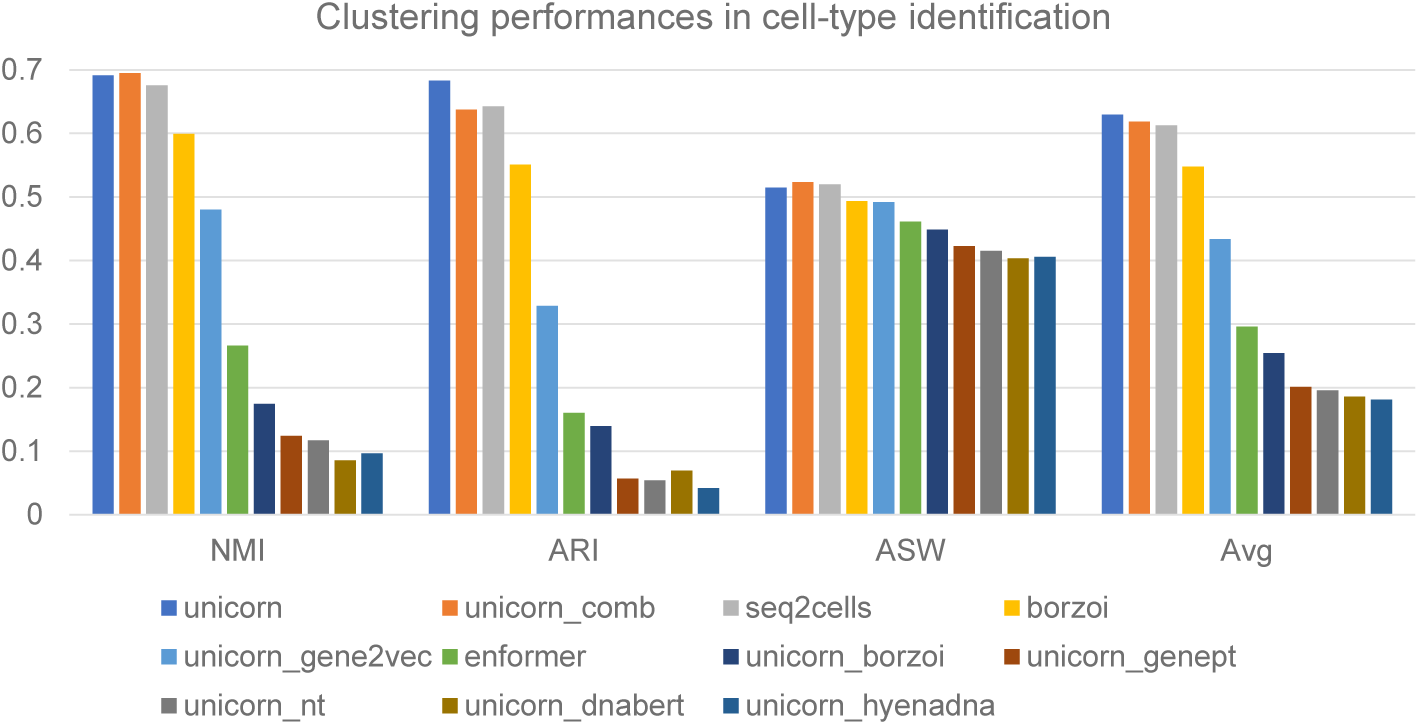
Comparison between UNICORN and other methods for cell clustering/cell-type identification. We have listed values from three metrics as well as averaged scores in this plot.

**Supplementary Fig. 6:**
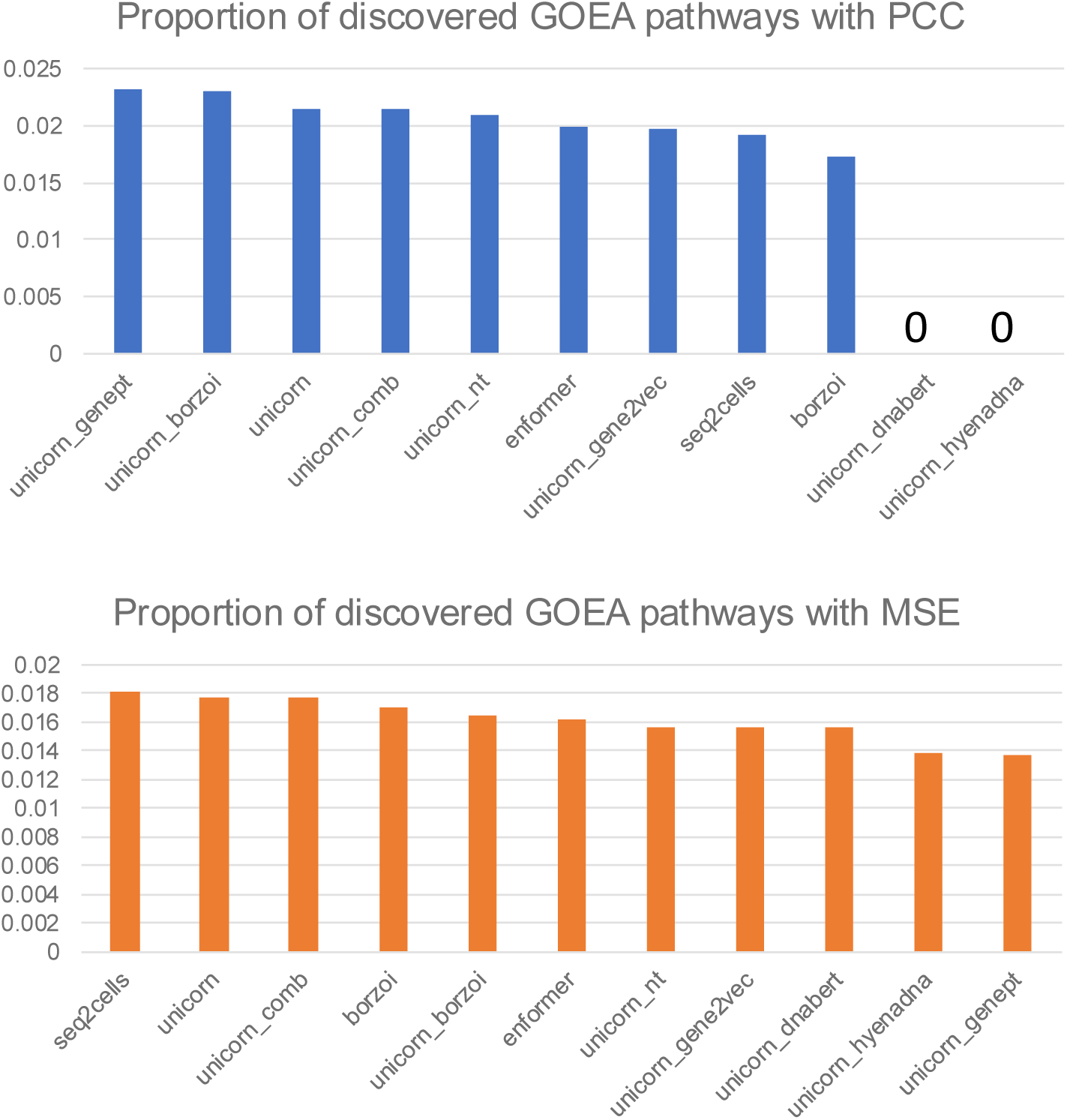
Comparison between UNICORN and other methods for identifying biological pathways. The upper panel represents the proportion of significant pathways versus all pathways by using PCC as threshold. The bottom panel represents the proportion of significant pathways versus all pathways by using MSE as threshold.

**Supplementary Fig. 7:**
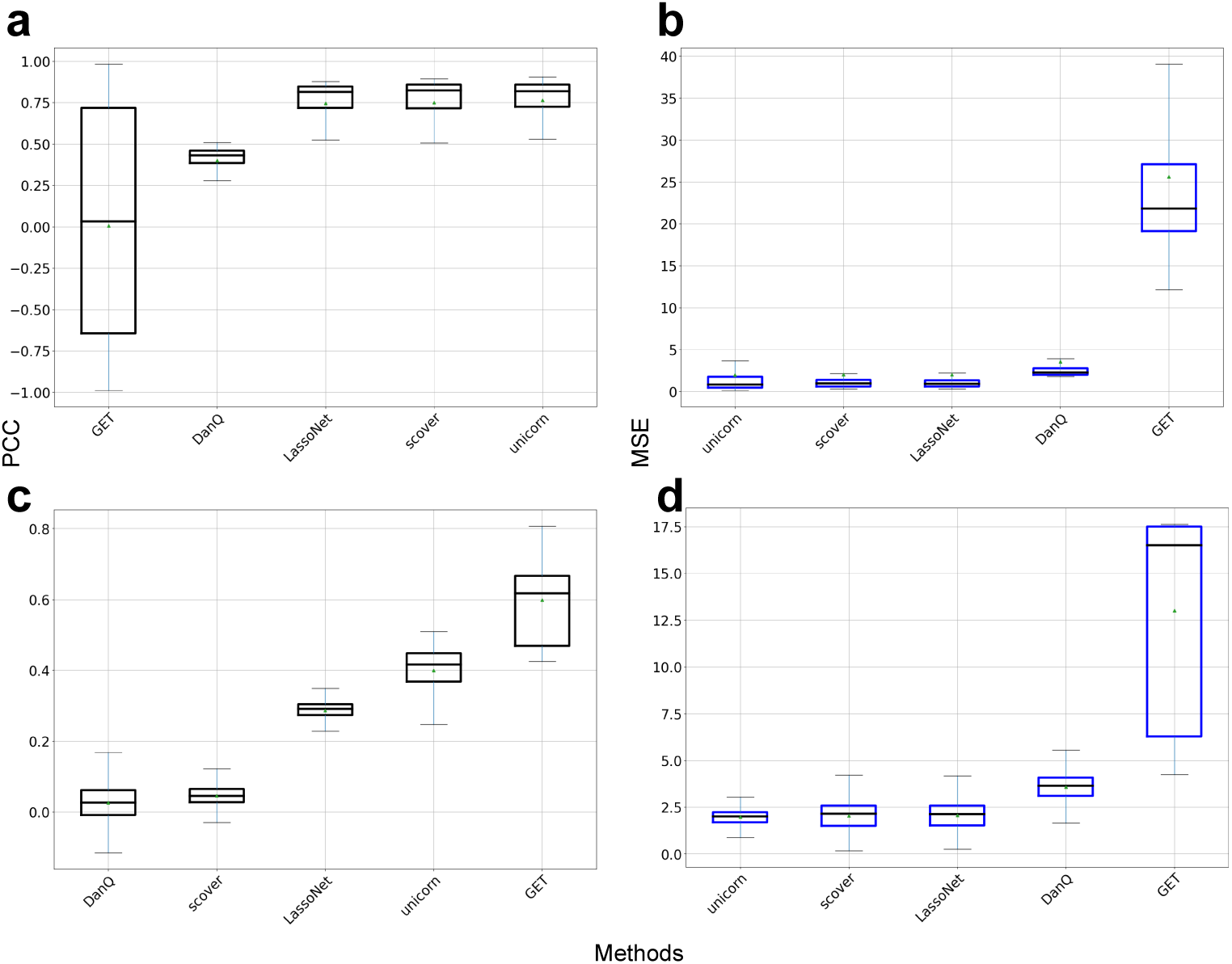
Comprehensive evaluations for gene expression predictions from sequences based on the Onek1k dataset. The triangle shape represents the mean value, and the black dashed line represents the median value. We report the results based on box plots. (a) Results of gene-level correlations across different methods. (b) Results of gene-level MSE across different methods. (c) Results of cell-level correlations across different methods. (d) Results of gene-level MSE across different methods.

**Supplementary Fig. 8:**
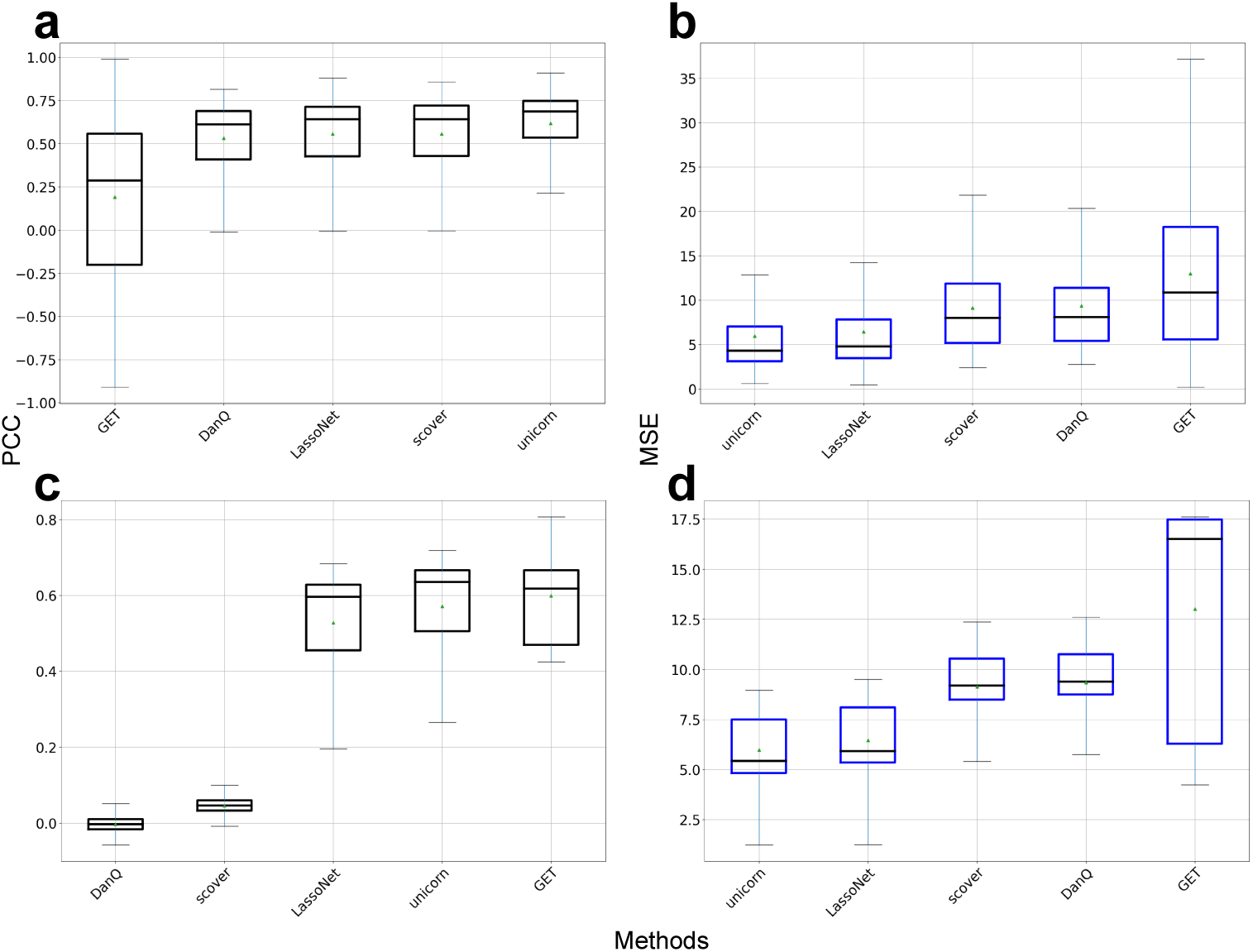
Comprehensive evaluations for gene expression predictions from sequences based on the ROSMAP dataset. The triangle shape represents the mean value, and the black dashed line represents the median value. We report the results based on box plots. (a) Results of gene-level correlations across different methods. (b) Results of gene-level MSE across different methods. (c) Results of cell-level correlations across different methods. (d) Results of gene-level MSE across different methods.

**Supplementary Fig. 9:**
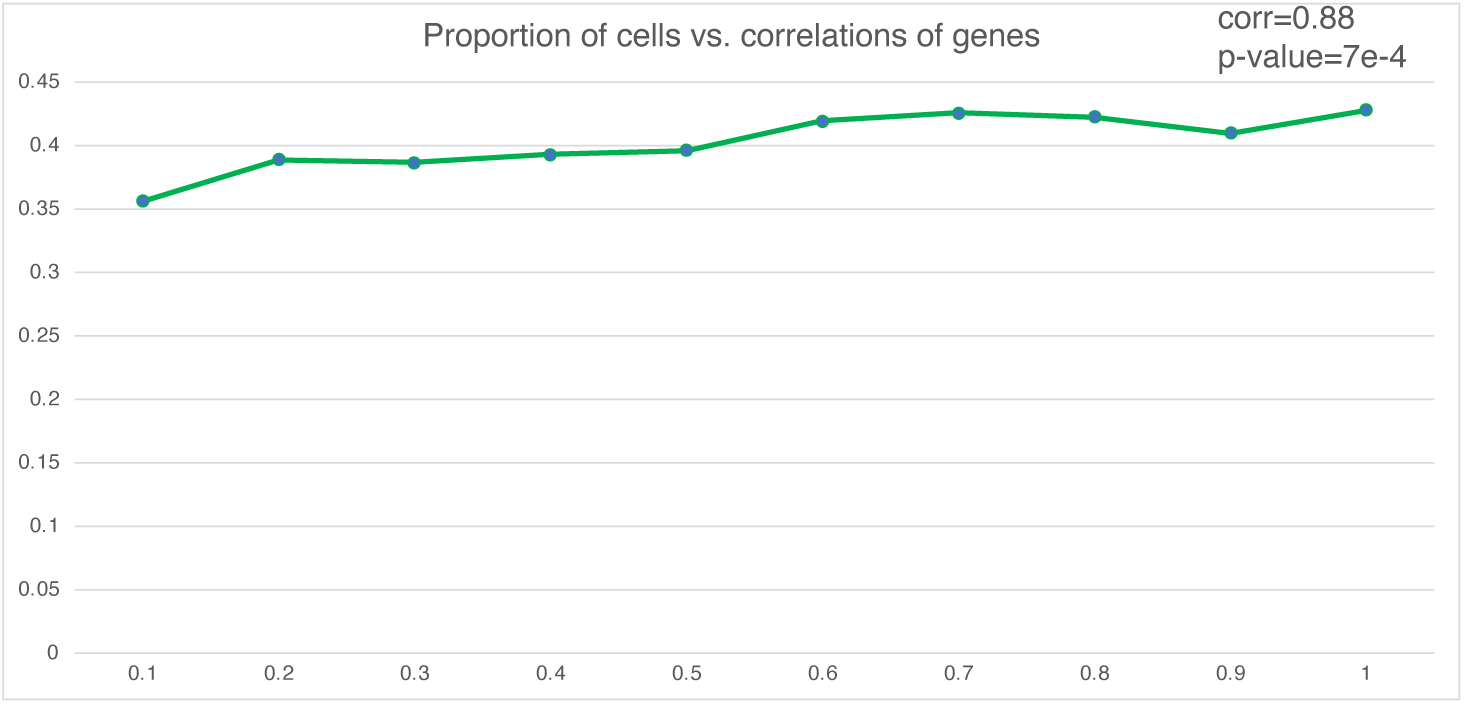
The relationship between the proportion of cells used for training and the correlation coefficients computed based on genes. The correlation is computed based on Pearson correlation.

**Supplementary Fig. 10:**
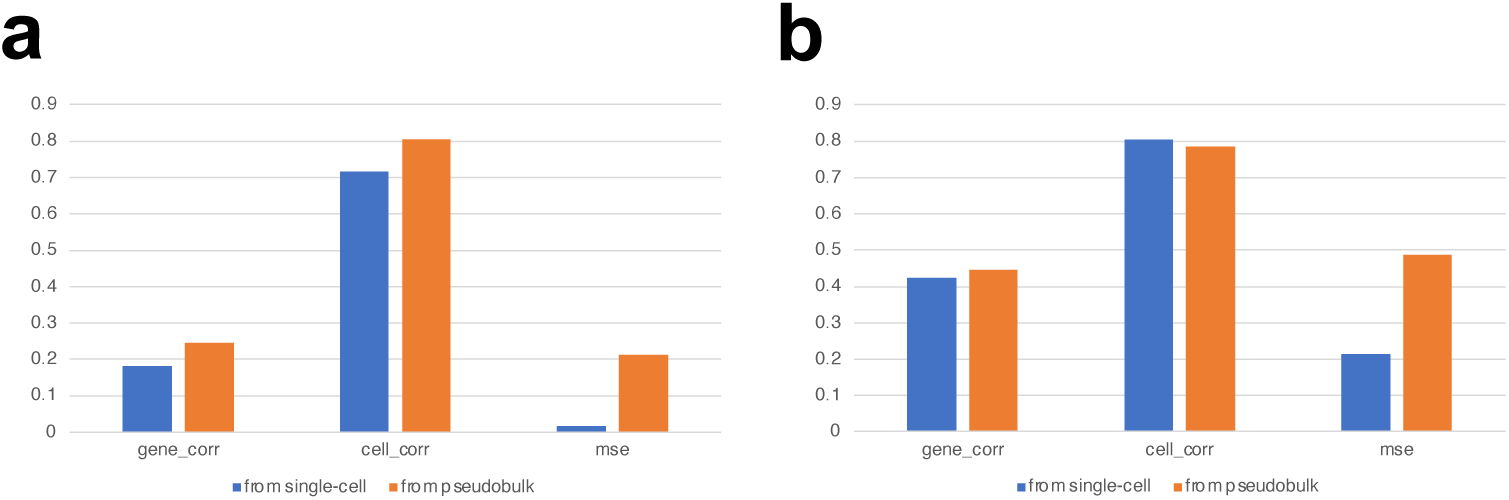
Comparisons between gene expression prediction based on single-cell data as input and gene expression prediction based on pseudo-bulk data as input. Both results are evaluated based on the pseudo-bulk level. (a): Comparisons of different metrics between two settings based on the thymus dataset. (b): Comparisons of different metrics between two settings based on the PBMC dataset.

**Supplementary Fig. 11:**
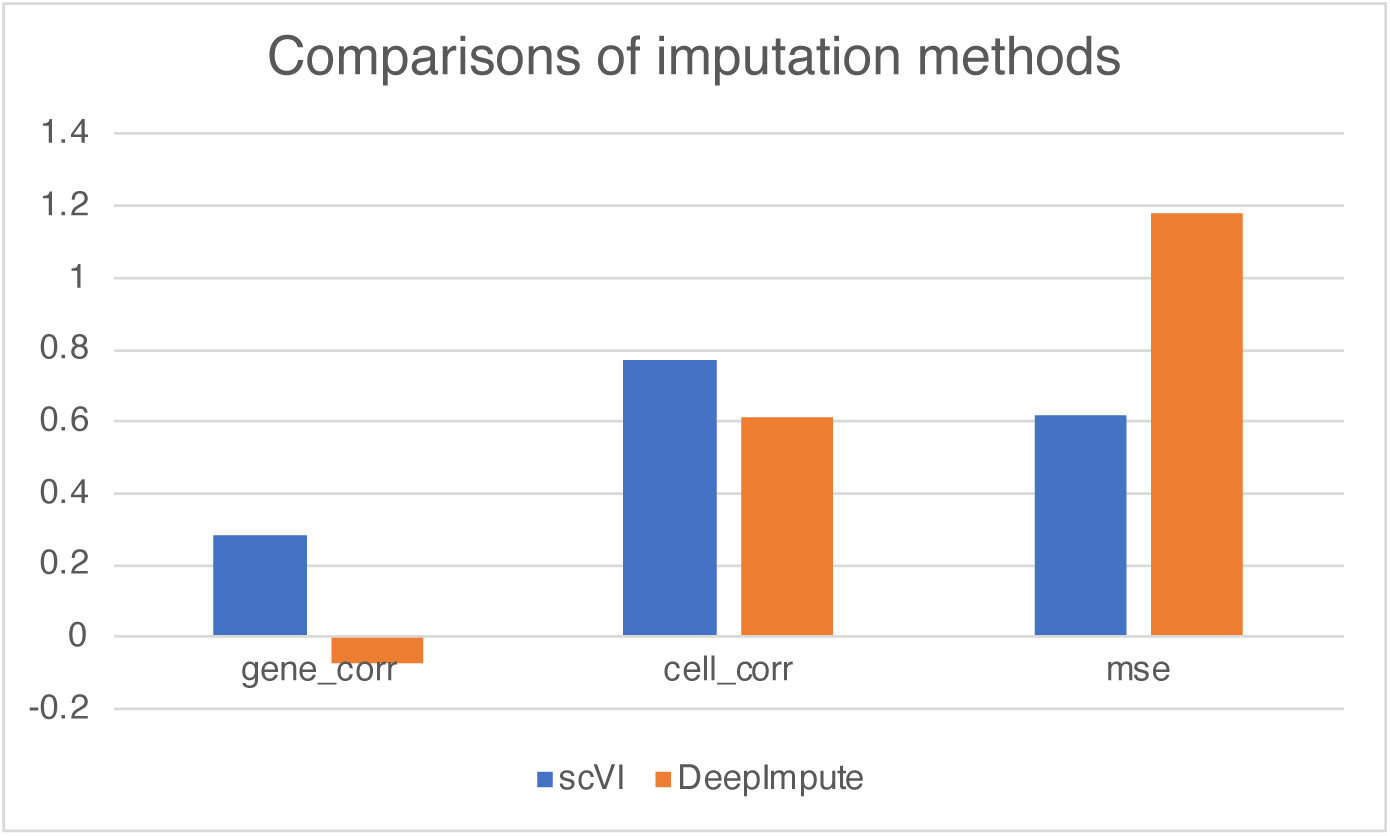
MSE under different imputation methods.

**Supplementary Fig. 12:**
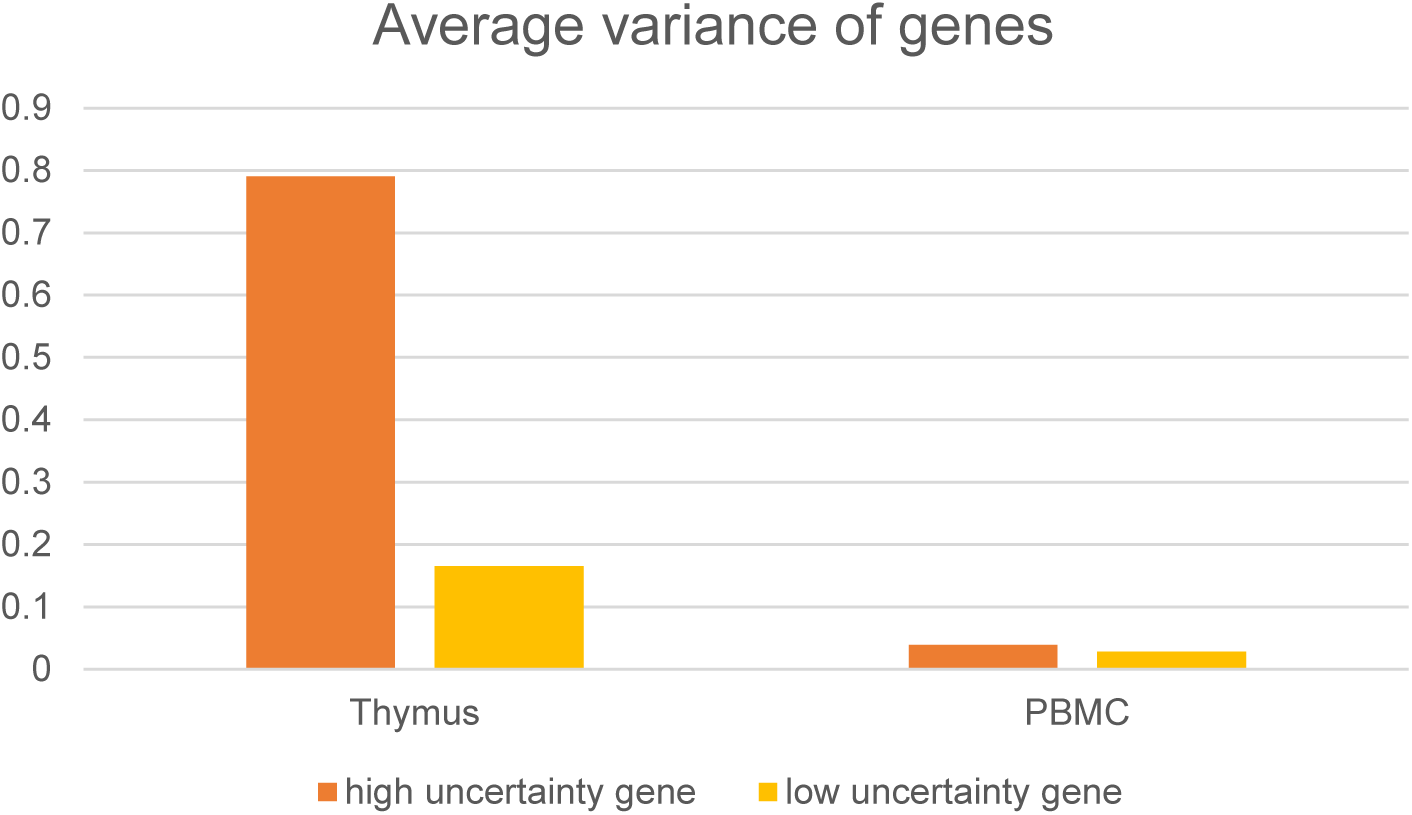
Average variance of genes for the two different datasets.

**Supplementary Fig. 13:**
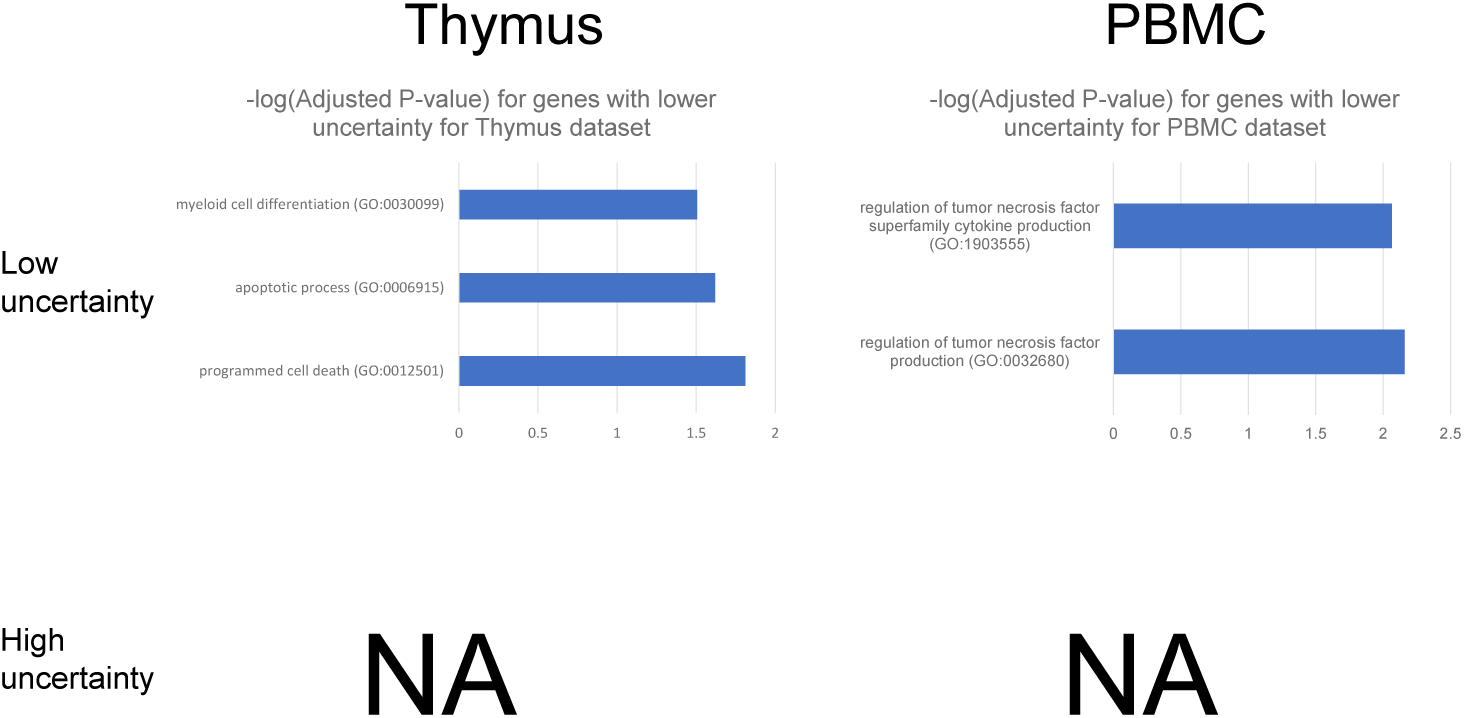
Tissue-specific GO Enrichment Analysis for genes with different uncertainty levels. NA means we cannot detect clear and research-supported tissue-specific pathways in this gene group.

**Supplementary Fig. 14:**
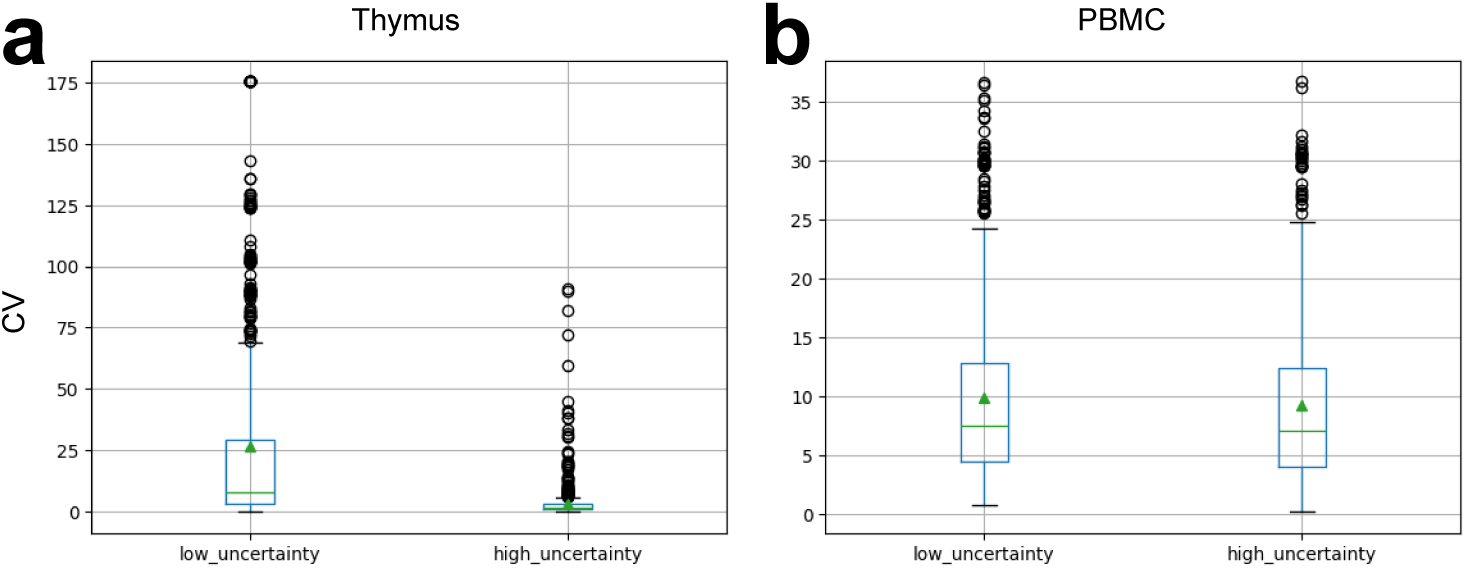
Comparisons of coefficients of variation (CV) for genes with different uncertainty levels for (a) thymus dataset and (b) PBMC dataset.

**Supplementary Fig. 15:**
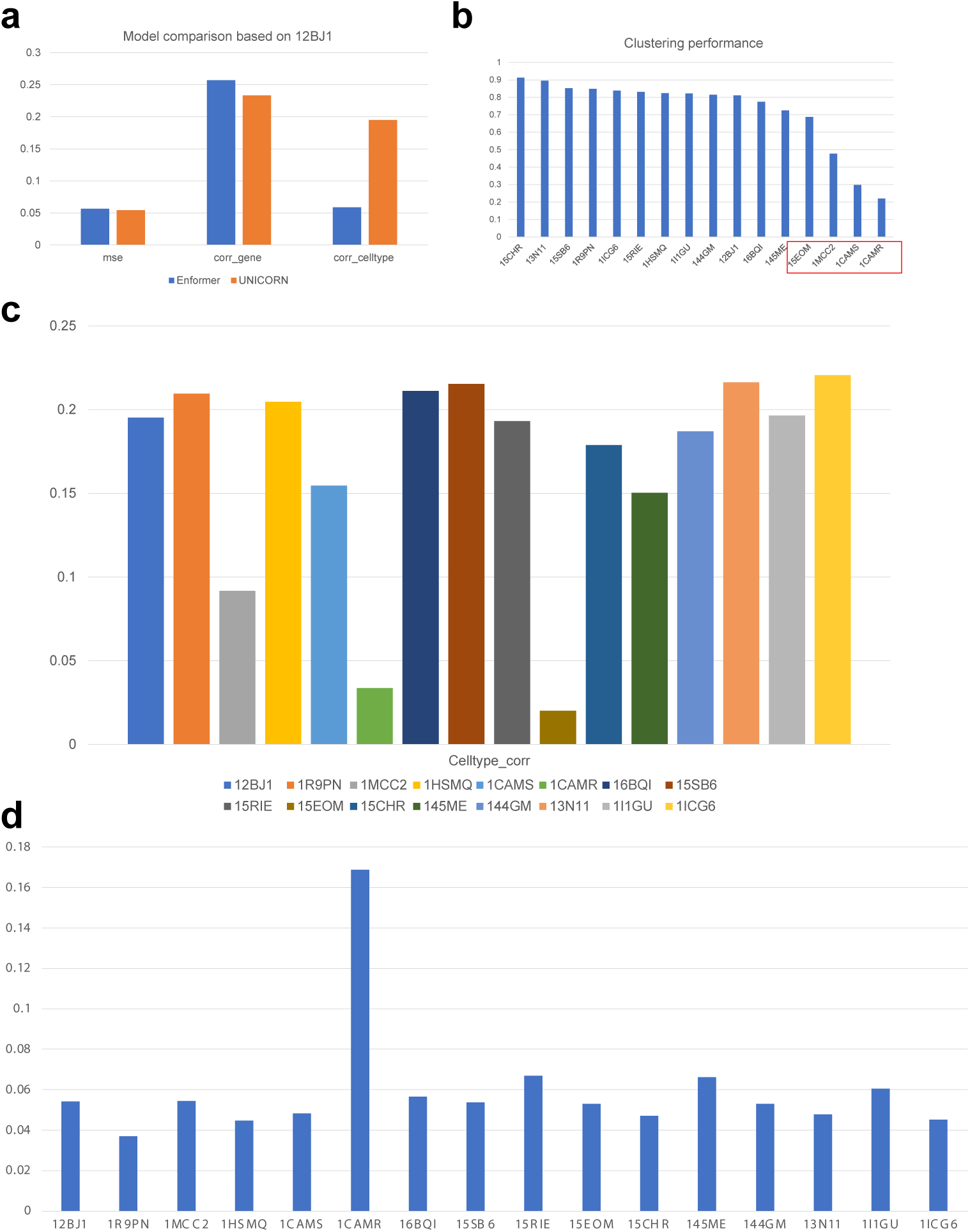
Analysis of gene expression predictions for individualized scRNA-seq datasets. (a) The performance comparison between Enformer and UNICORN based on the sample 12BJ1. (b) The average clustering scores across different individuals based on the corresponding scRNA-seq datasets. (c) The results of correlation coefficients computed based on cell types across different samples. (d) The results of MSE computed based on genes and cells across different samples.

**Supplementary Fig. 16:**
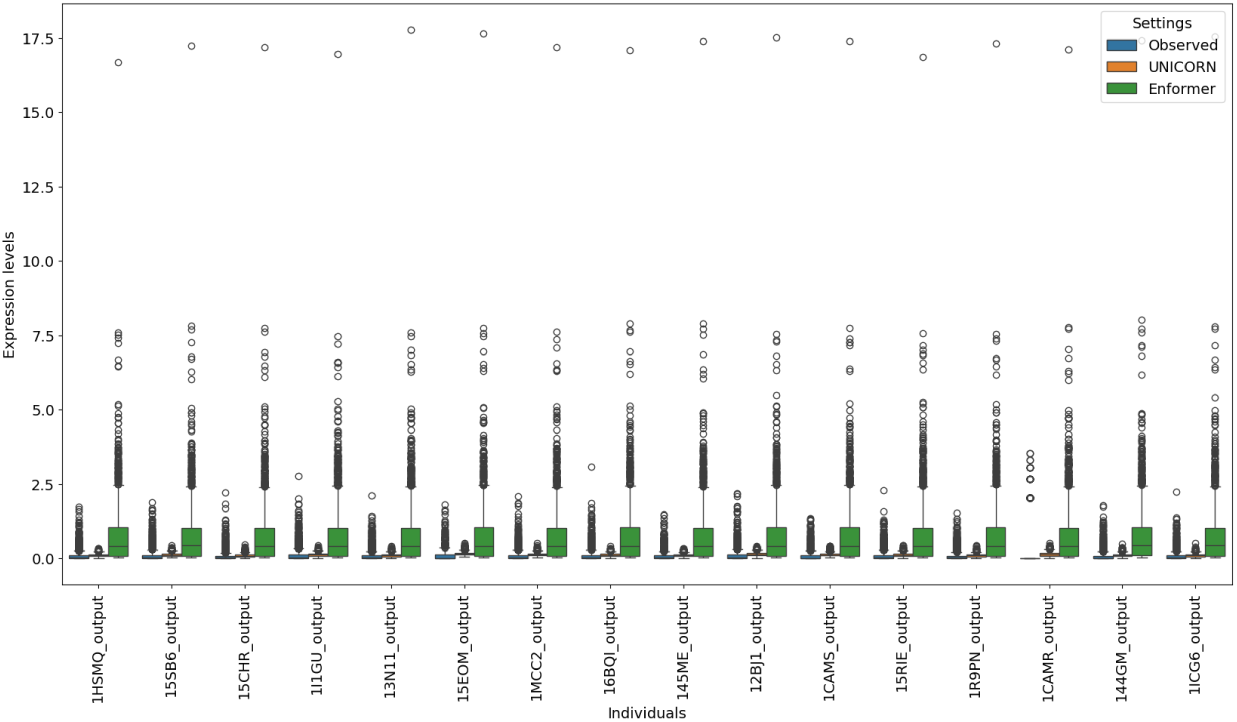
Comparisons of gene expression levels across individuals. We assign different colors for the observed values and predicted values.

**Supplementary Fig. 17:**
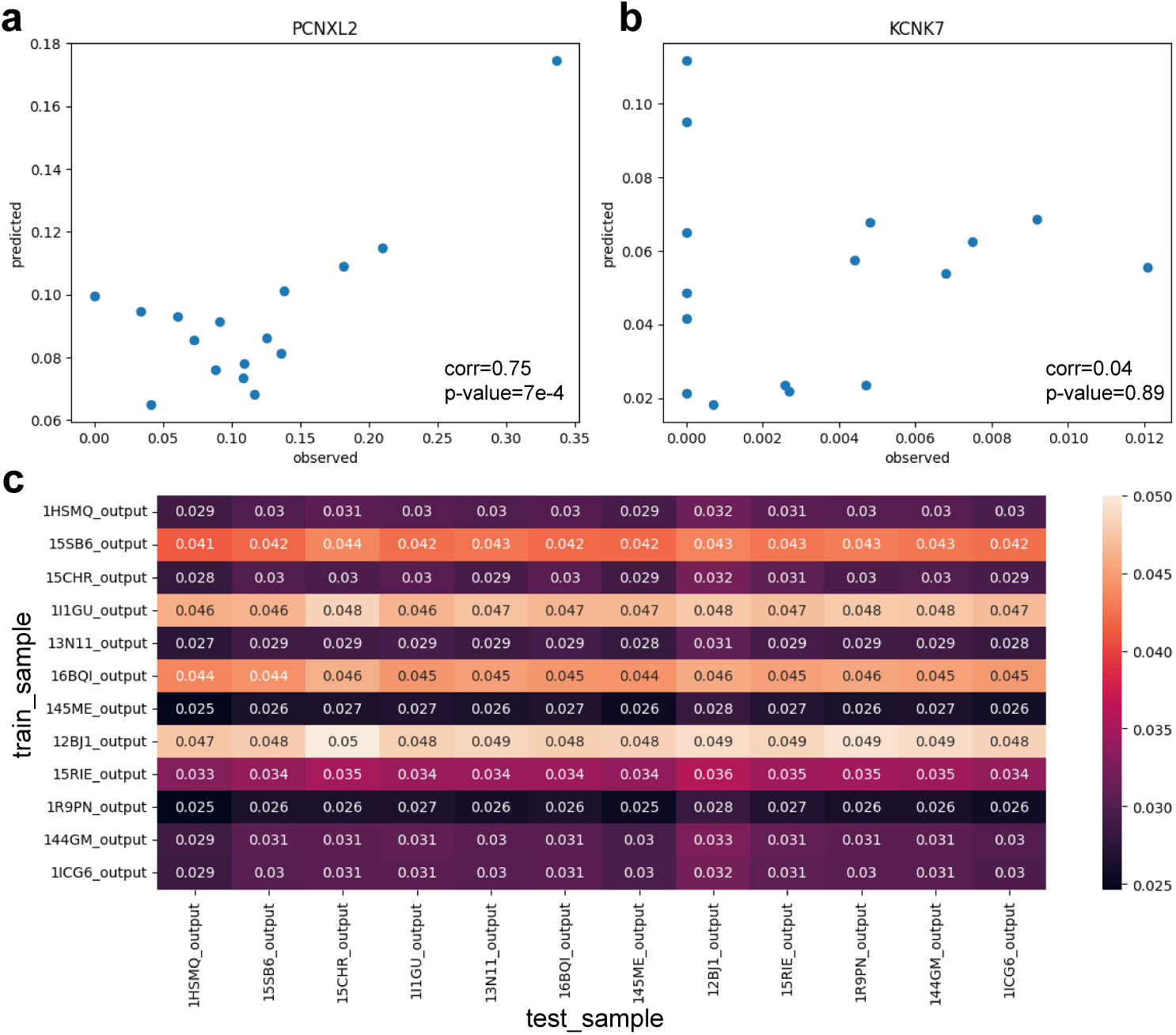
Examples of genes with different prediction performances and the results of cross-individual prediction. (a) The correlation between observed PCNXL2 expression and predicted PCNXL2 expression (good example). (b) The correlation between observed KCNK7 expression and predicted KCNK7 expression (poor example). (c) The MSE value of cross-individual prediction result based on UNICORN. The rows representing training samples and the columns represent the testing samples.

**Supplementary Fig. 18:**
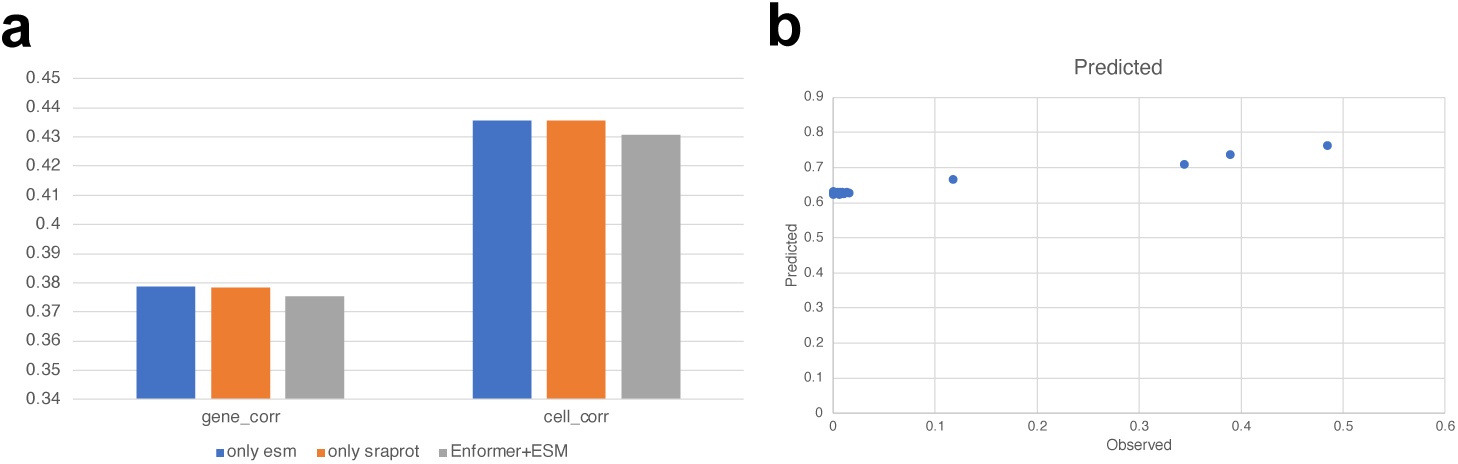
Analysis the factors in affecting multi-omic prediction. (a) The ablation test for different approaches of embedding combination. Here ESM2 [71] and SaProt [97] are two models used to generate protein embeddings. (b) The scatter plot for visualizing the relationship between observed peak information and predicted peak information for different cell types measured in 10X Multi-Omic dataset.

**Supplementary Fig. 19:**
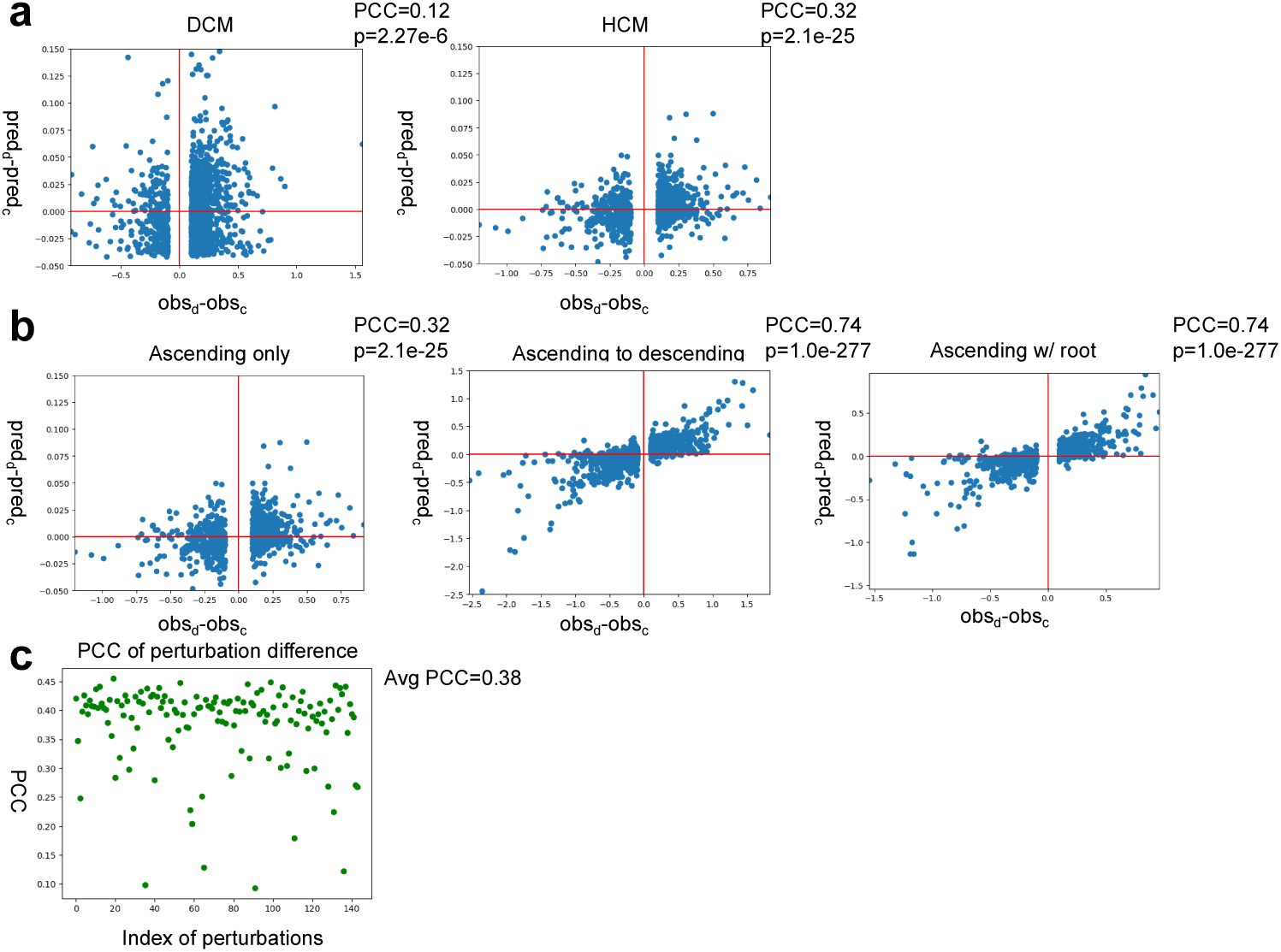
Relationship between the difference computed based on the observed and predicted levels across multiple conditions. (a) Relationship computed based on the Heart dataset. (a) Relationship computed based on the Aorta dataset. (c) Relationship between the perturbation index and PCC between two differences computed based on the perturbation dataset. Necessary statistics such as PCC and p-value (p) are annotated in the corresponding panel.

**Supplementary Fig. 20:**
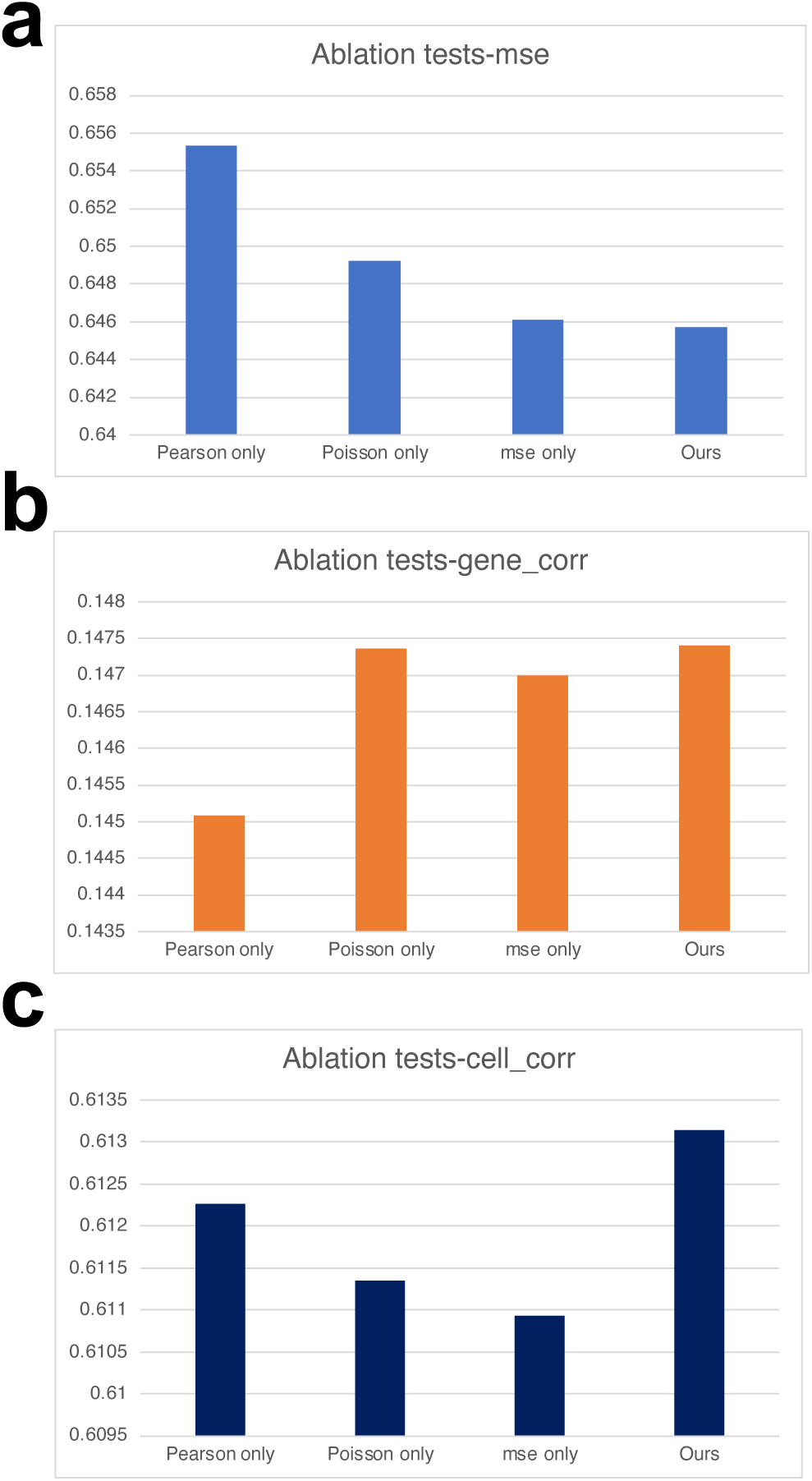
Results of ablation tests for multi-modal prediction. We report the averaged score for each metric. (a) The correlation coefficients of genes under different choices of loss function components. (b) The correlation coefficients of cells under different choices of loss function components. (c) The MSE under different choices of loss function components.

**Supplementary Fig. 21:**
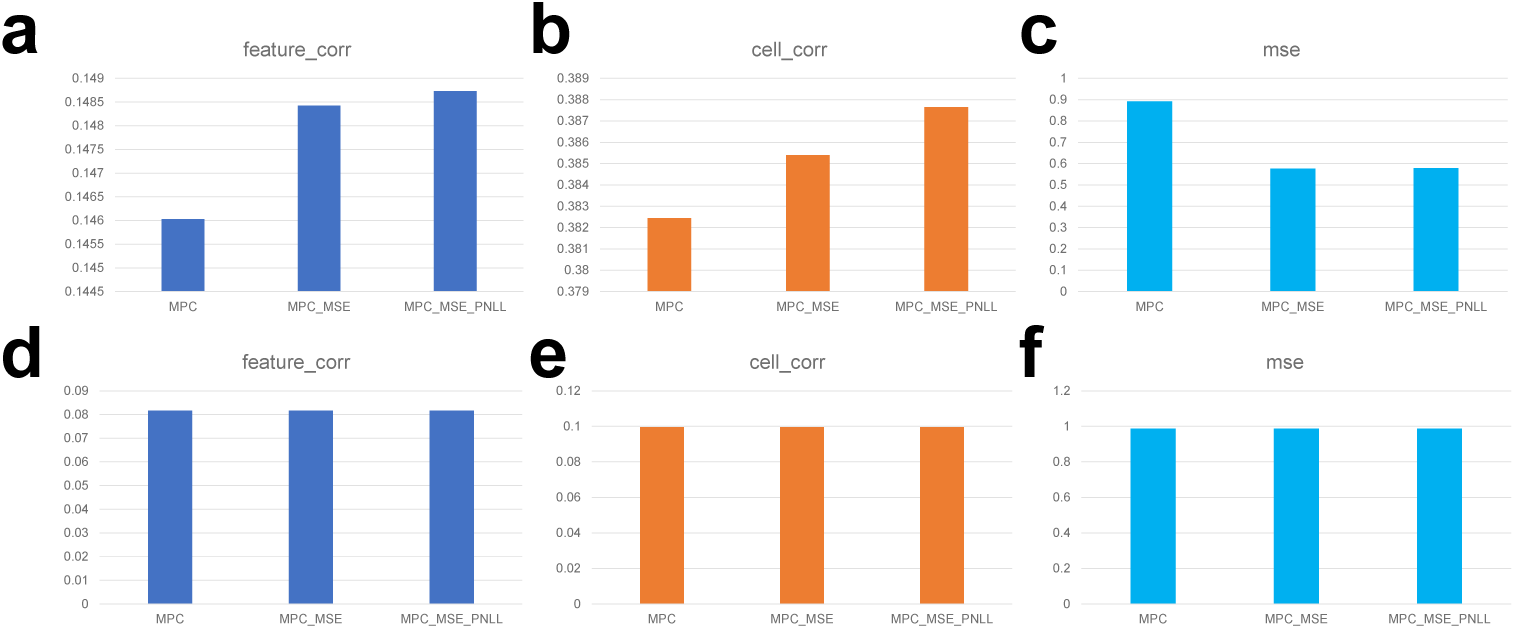
Results of ablation tests for gene expression prediction. We report the averaged score for each metric. (a) The correlation coefficients of features under different choices of loss function components based on 10X Multiome data. (b) The correlation coefficients of cells under different choices of loss function components based on 10X Multiome data. (c) The MSE under different choices of loss function components based on 10X Multiome data. (d) The correlation coefficients of features under different choices of loss function components based on CITE-seq data. (e) The correlation coefficients of cells under different choices of loss function components based on CITE-seq data. (f) The MSE under different choices of loss function components based on CITE-seq data.

**Supplementary Fig. 22:**
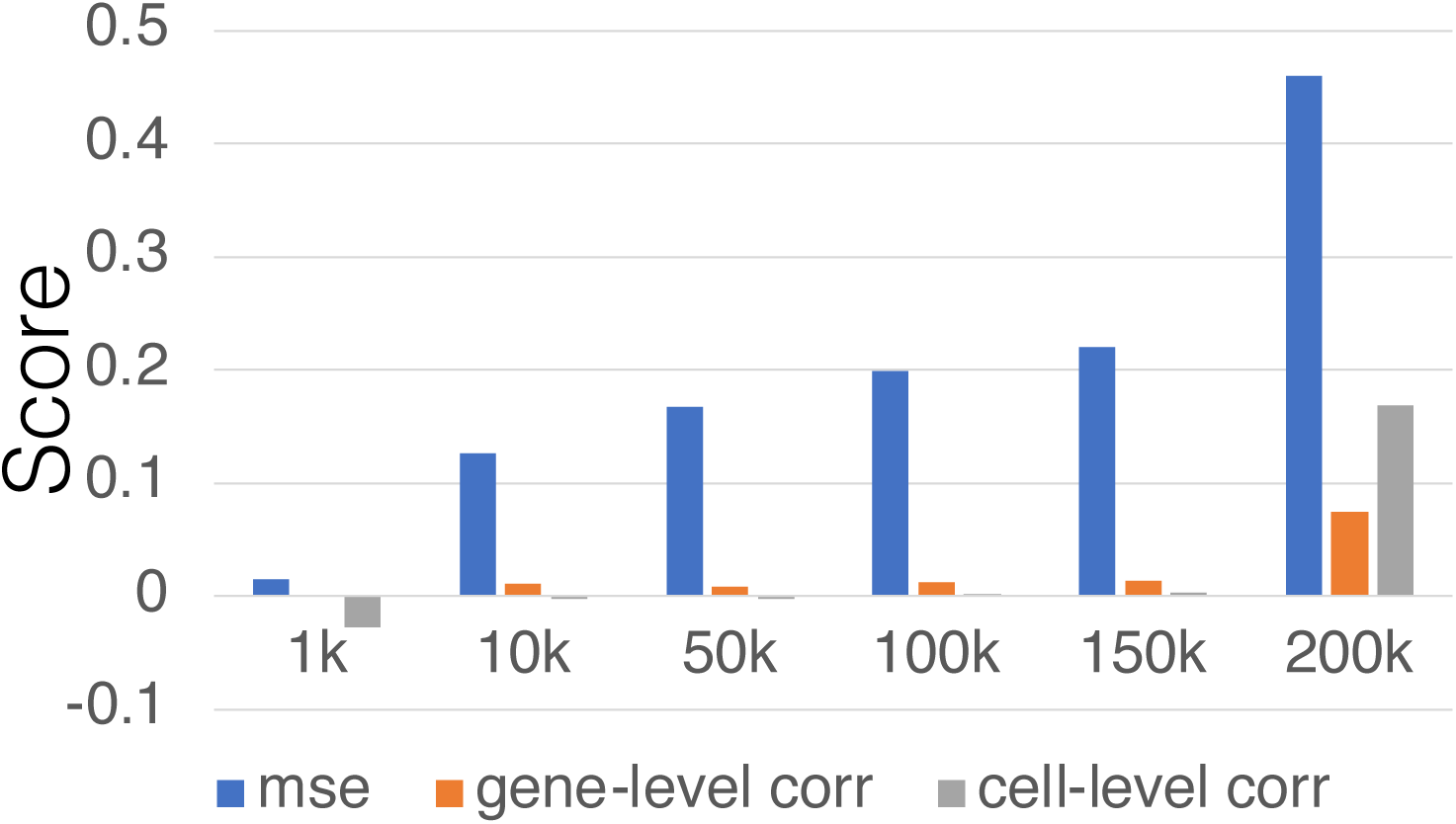
Comparisons for single-cell gene expression prediction under different context lengths. We visualize three metrics (MSE, gene-level correlation score, and cell-level correlation score) by showing their averaged values under different context lengths from 1k to 200k.

**Supplementary Fig. 23:**
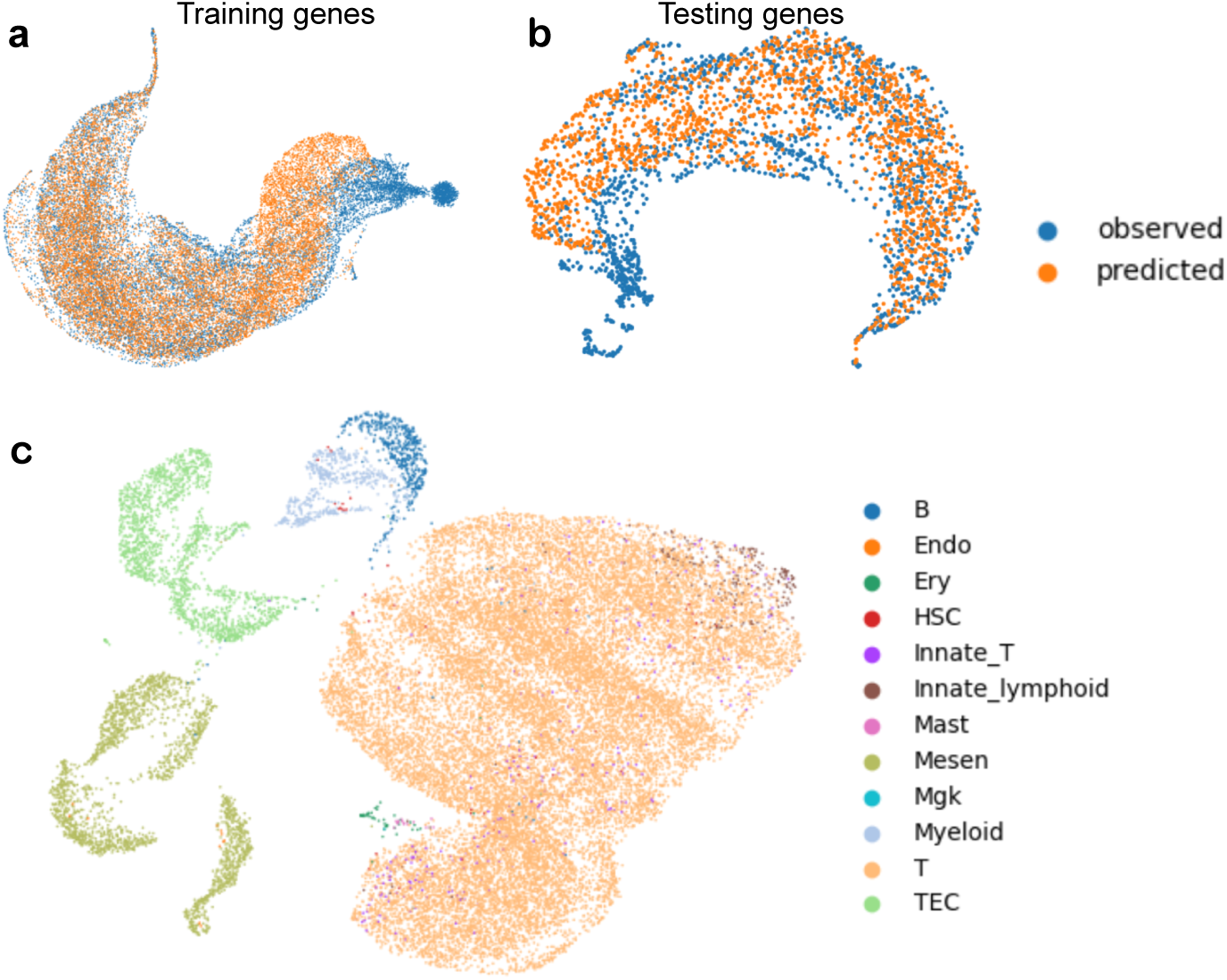
The visualization results of atlas-level data prediction. (a) The UMAP plots for training gene embeddings from observed gene expressions and predicted gene expressions. (b) The UMAP plots for testing gene embeddings from observed gene expressions and predicted gene expressions. (c) The UMAP plots for cells with predicted expression levels from the thymus atlas dataset, colored by cell types.

**Supplementary Fig. 24:**
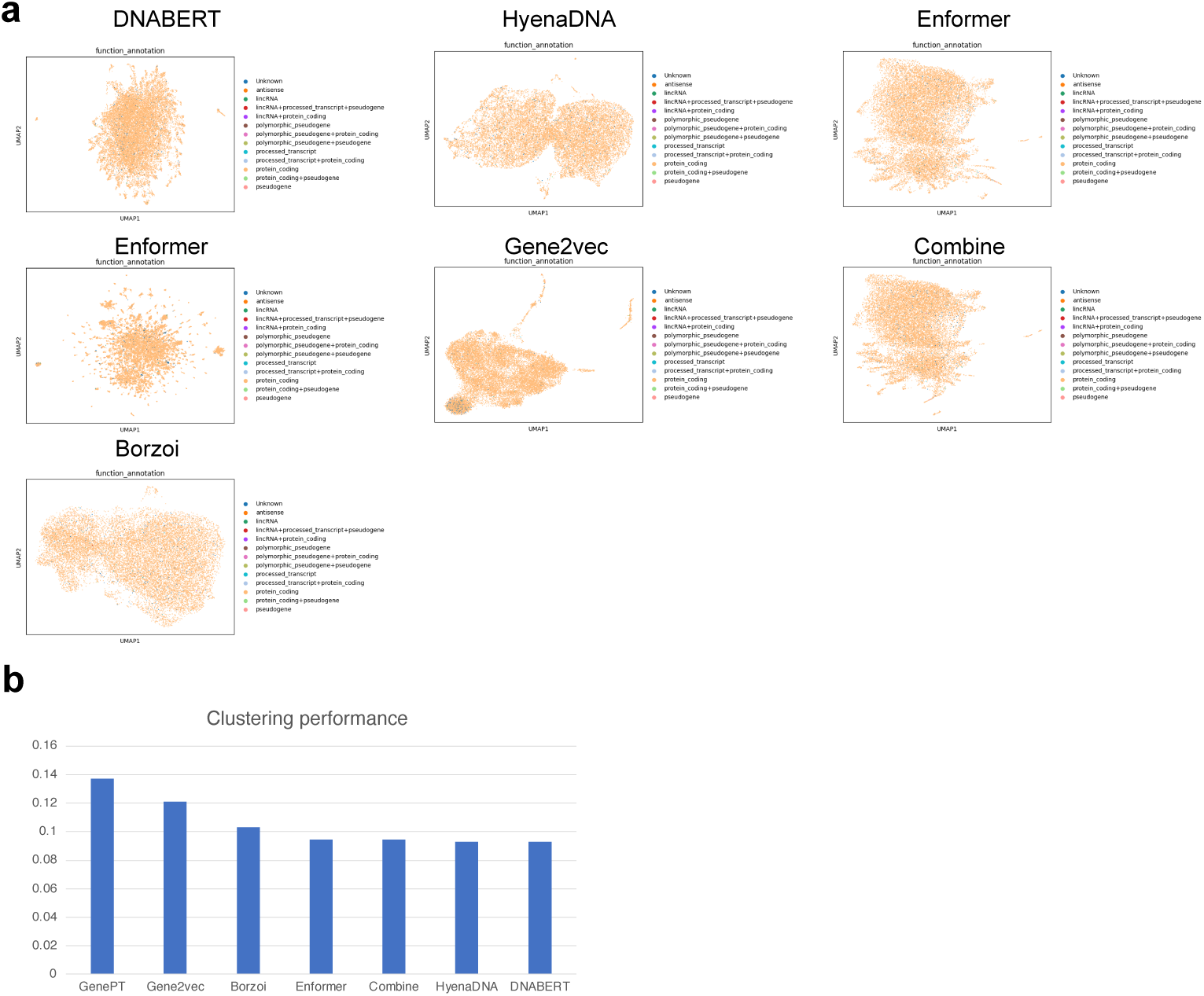
Performances of clustering based on different gene embeddings. (a) The UMAP plots of different gene embeddings colored by functional annotations. (b) The averaged scores of different gene embeddings.

**Supplementary Fig. 25:**
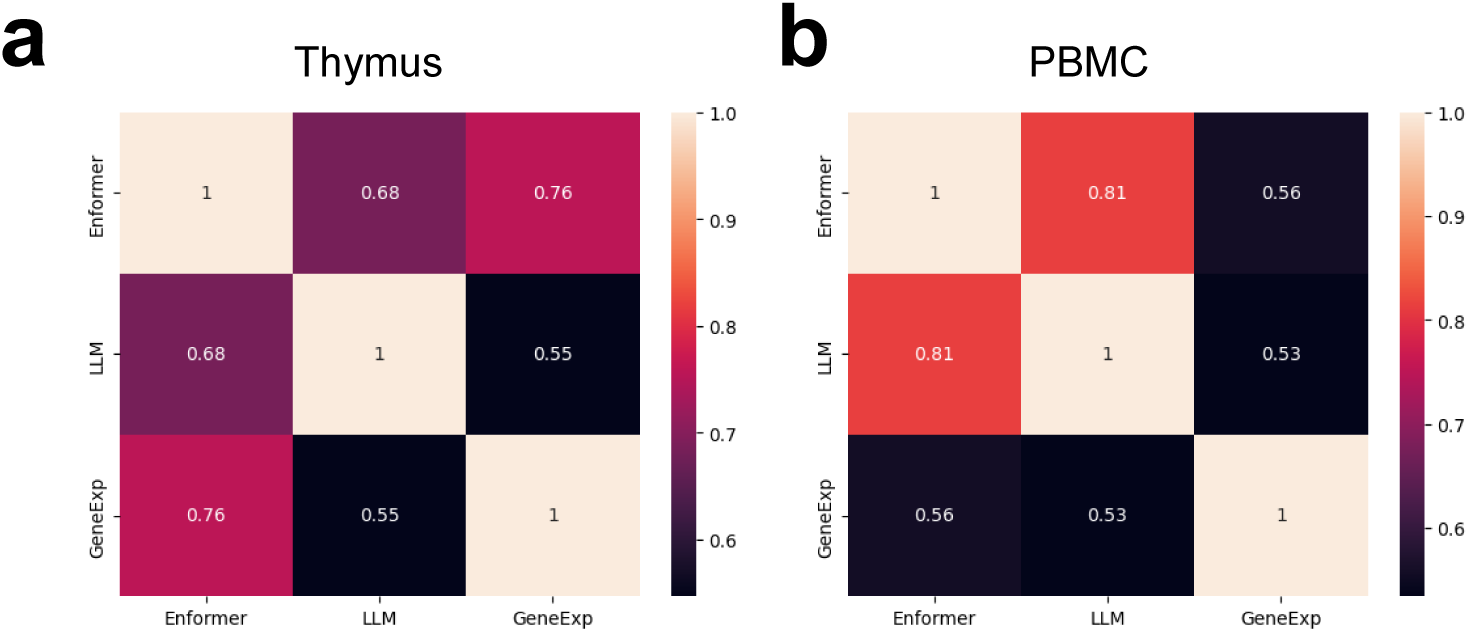
Comparison of the gene-level similarity from Enformer, LLM, and gene expression profiles. (a) represents the results computed based on thymus dataset, and (b) represents the results computed based on PBMC dataset.

**Supplementary Fig. 26:**
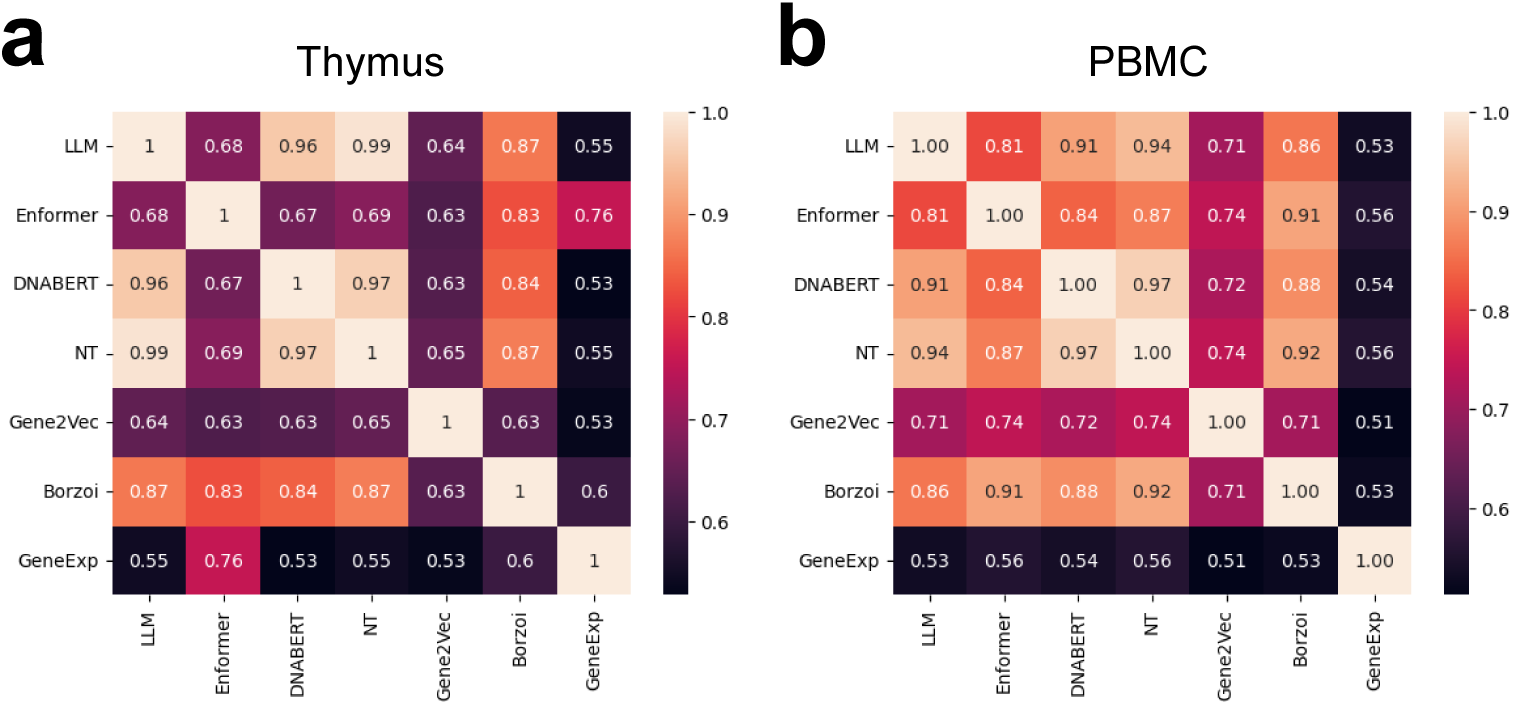
Comparison of the gene-level similarity from all base models and expression profiles used in this manuscript. (a) represents the results computed based on thymus dataset, and (b) represents the results computed based on PBMC dataset.

**Supplementary Fig. 27:**
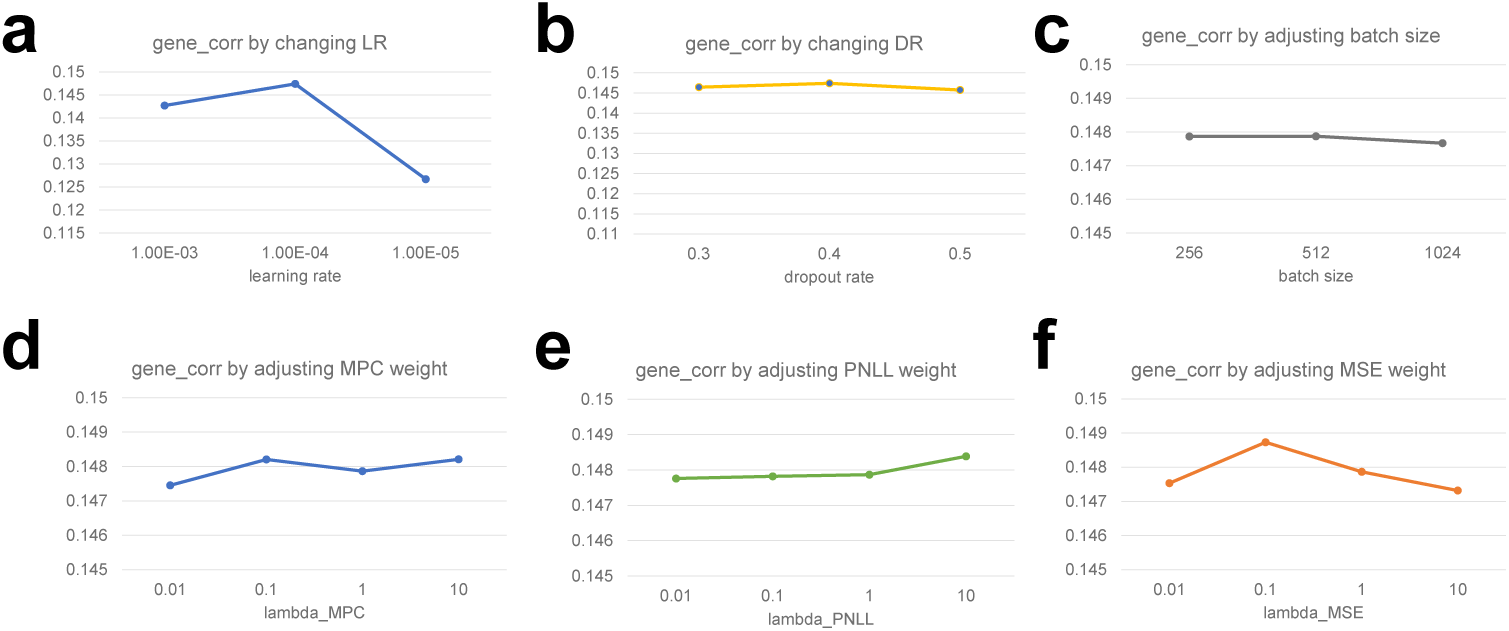
Model performances under different settings of hyperparameters. (a) The effect of changing LR. (b) The effect of changing DR. (c) The effect of changing batch size. (d) The effect of changing the weight of L*_MP C_*. (e) The effect of changing the weight of L*_P NLL_*. (f) The effect of changing the weight of L*_MSE_*.

**Supplementary Fig. 28:**
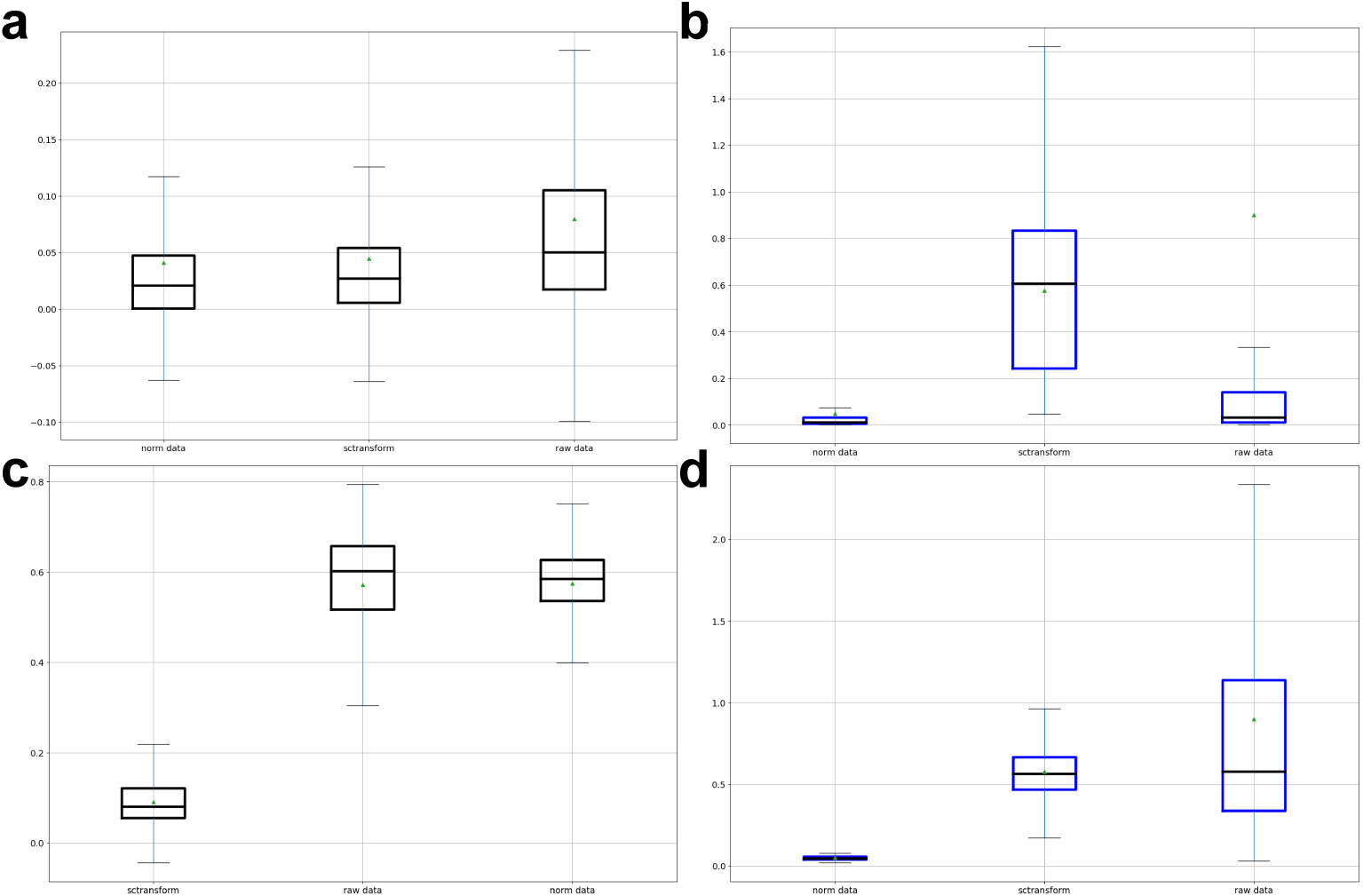
Comparisons between raw data mode (raw data), lognormalized data mode (norm data), and sctransform data mode (sctransform). The triangle shape represents the mean value, and the black dashed line represents the median value. We report the results based on box plots. (a) Results of gene-level correlations across different methods based on the PBMC dataset. (b) Results of gene-level MSE across different methods based on the PBMC dataset. (c) Results of cell-level correlations across different methods based on the PBMC dataset. (d) Results of cell-level MSE across different methods based on the PBMC dataset.

**Supplementary Fig. 29:**
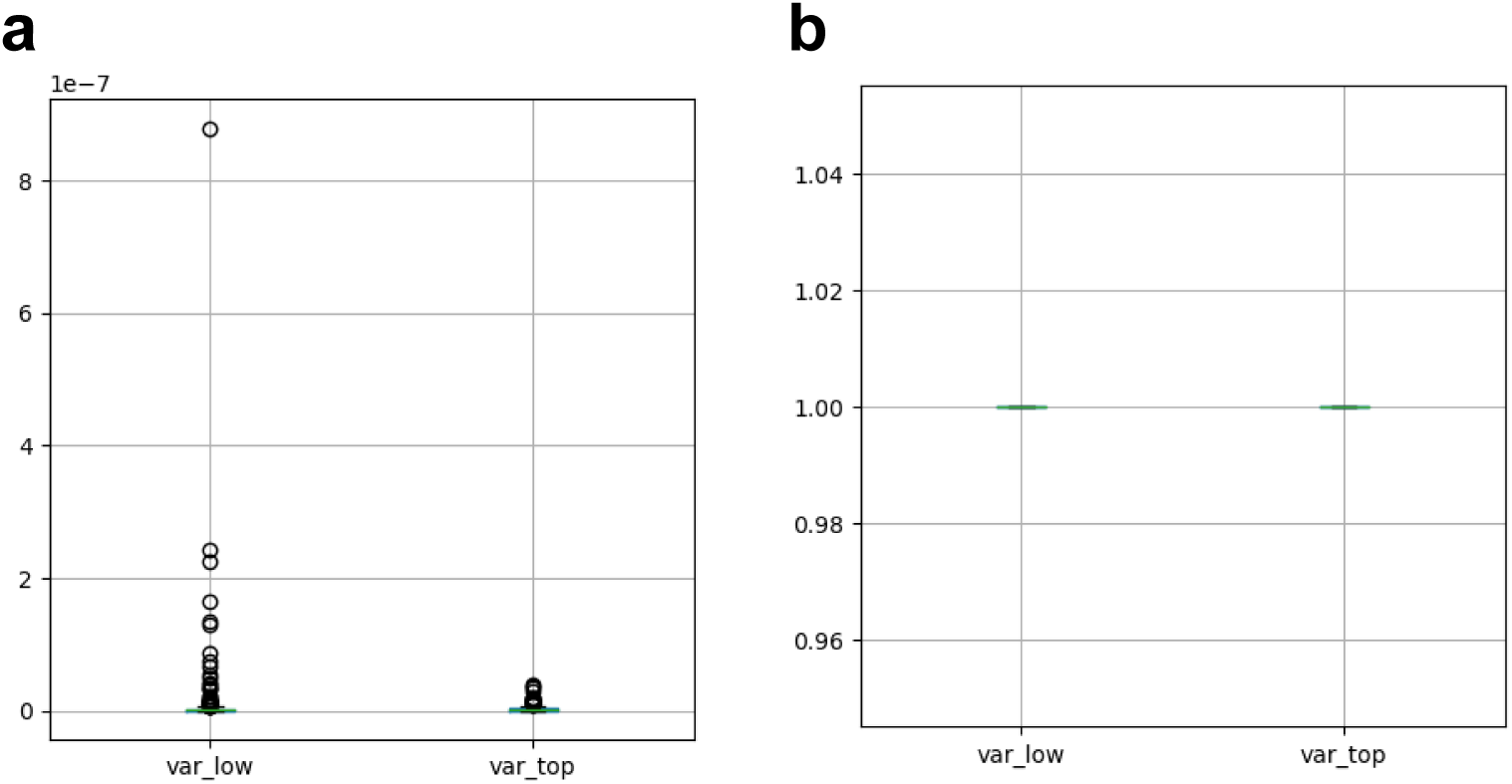
The difference of metrics computed based on effective allele and non-effective allele. Here the legend *var low* represent last 100 alleles and the legend *var top* represents the top 100 alleles. The gene expression is computed at the pseudo-bulk level. (a) The distribution of MSE between the predicted gene expression and the observed gene expression. (b) The distribution of gene-level correlation coefficients between the predicted gene expression and the observed gene expression.

**Supplementary Fig. 30:**
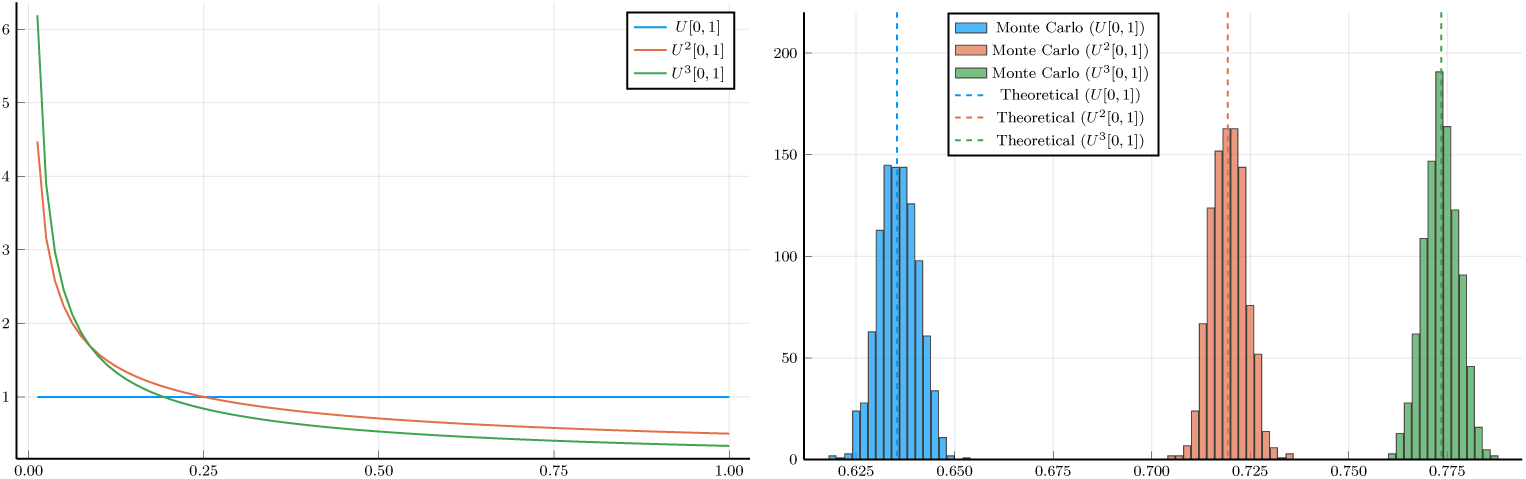
*(Left)* The density function of *U ^k^, k* = 1, 2, 3. *(Right)* Theoretical probability and the approximation by 1000 Monte Carlo experiments on *n* = 1000 samples.

**Supplementary Fig. 31:**
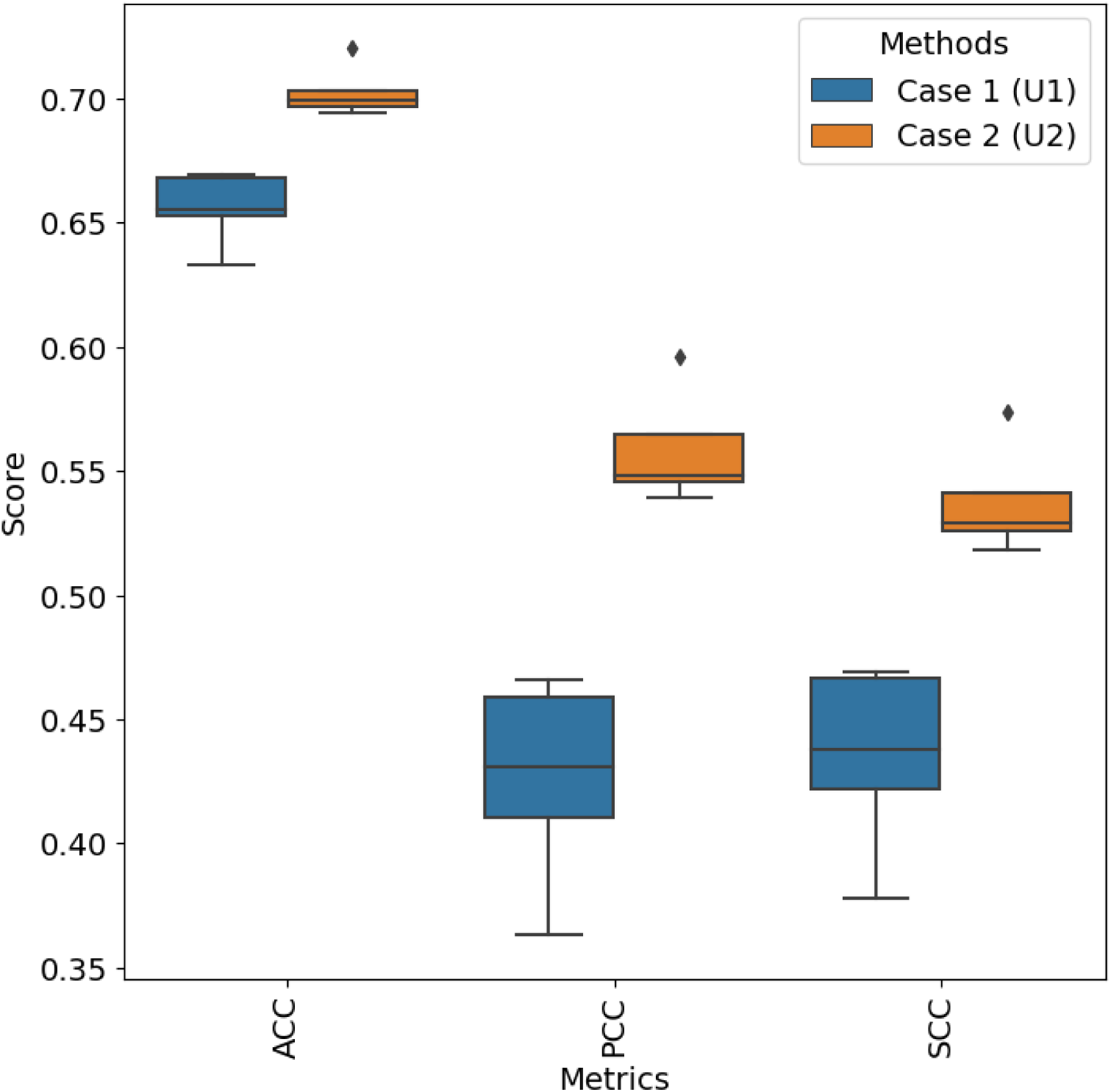
Simulation results for verifying the uncertainty estimation approach. Here we consider two different cases based on three different metrics. Higher scores mean better estimation.

