## appendix for "UNICORN: Towards Universal Cellular Expression Prediction with a Multi-Task Learning Framework"

### G Supplementary figures

| Models | Support different sequence embeddings? | Support multi-omic expression prediction? | Support uncertainty estimation? | Support individual sequence input? | Trainable? | Distribution-aware loss function design? | Conditional cell-type analysis? | Minimal-required GPU Type |
|---|---|---|---|---|---|---|---|---|
| **UNICORN** | Y | Y | Y | Y | Y | Y | Y | GTX 1080Ti (16GB) |
| Seq2cells | N | N | N | Y | Y | N | N | GTX 1080Ti (16GB) |
| Enformer | N | N | N | Y | Y | N | N | A100 (40GB) |
| Borzoi | N | N | N | Y | Y | N | N | A100 (40GB) |
| Decima | N | N | N | N | N | N | N | NA |

**Extended Data Fig. 1** Comparison of different models based on their advantages in design and implementation.

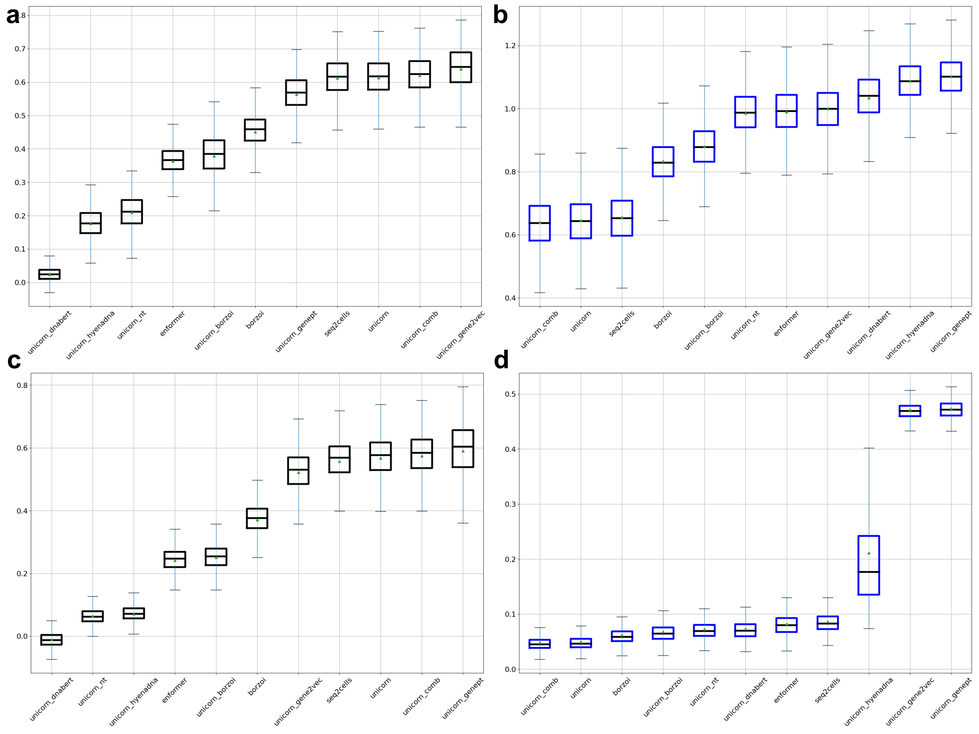


**Extended Data Fig. 2** Comprehensive evaluations for gene expression predictions from sequences. The triangle shape represents the mean value, and the black dashed line represents the median value. We report the results based on box plots. (a) Results of cell-level correlations across different methods based on the thymus dataset. (b) Results of MSE across different methods based on the thymus dataset. (c) Results of cell-level correlations across different methods based on the PBMC dataset. (d) Results of MSE across different methods based on the PBMC dataset.

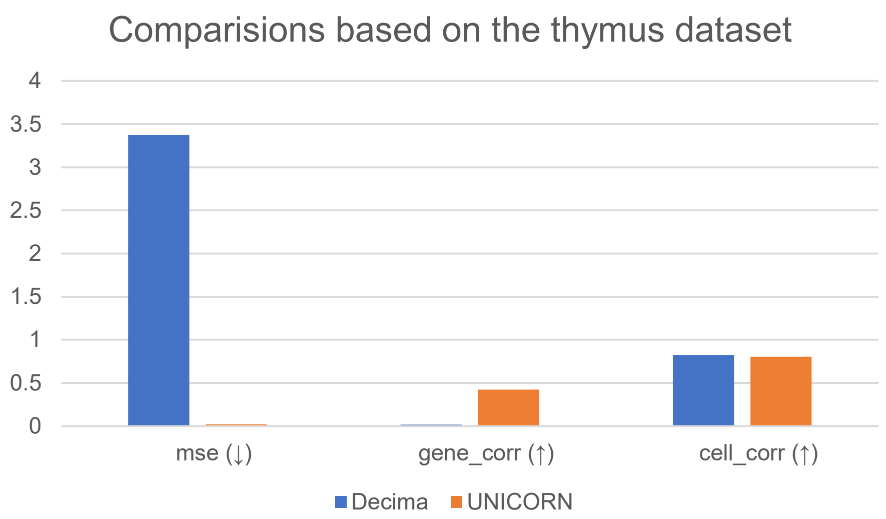


**Extended Data Fig. 3** Comparison between UNICORN and Decima based on the thymus dataset. To make a fair benchmark, we select the overlapped cell types and tissue from Decima's prediction results.

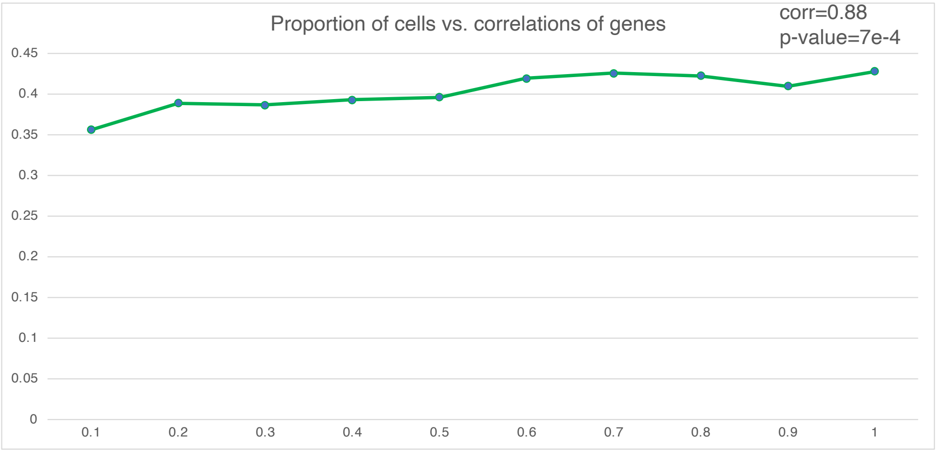


**Extended Data Fig. 4** The relationship between the proportion of cells used for training and the correlation coefficients computed based on genes. The correlation is computed based on Pearson correlation.

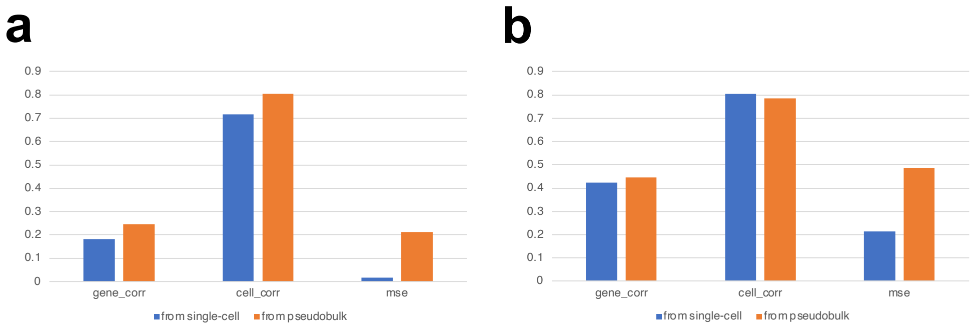


**Extended Data Fig. 5** Comparisons between gene expression prediction based on single-cell data as input and gene expression prediction based on pseudo-bulk data as input. Both results are evaluated based on the pseudo-bulk level. (a): Comparisons of different metrics between two settings based on the thymus dataset. (b): Comparisons of different metrics between two settings based on the PBMC dataset.

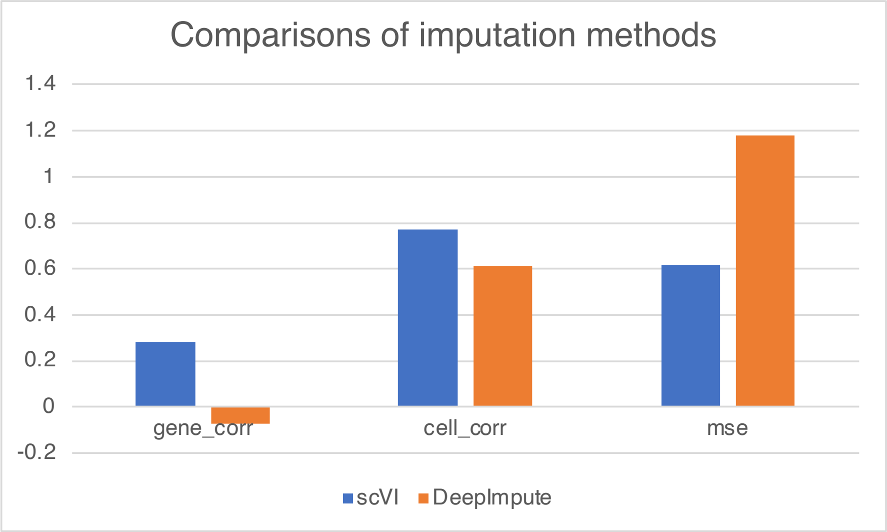


**Extended Data Fig. 6** MSE under different imputation methods.

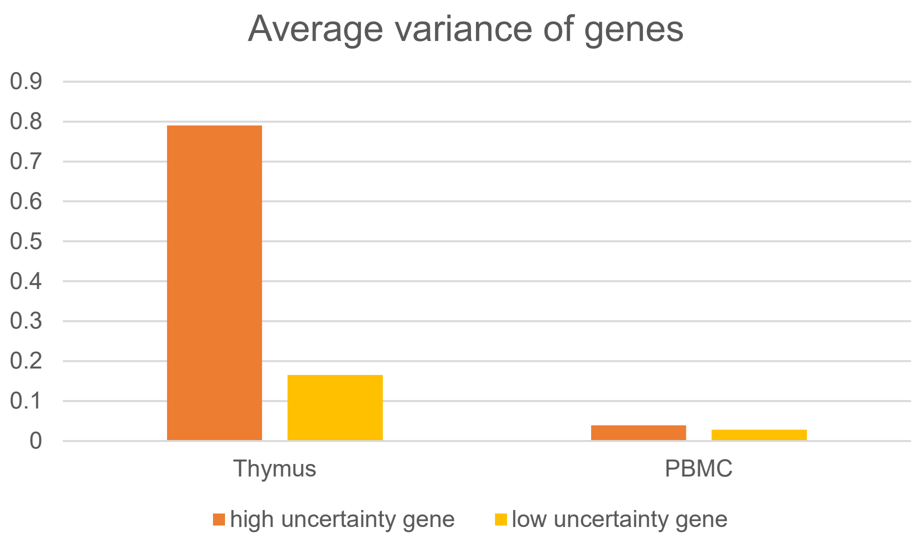


**Extended Data Fig. 7** Average variance of genes for the two different datasets.

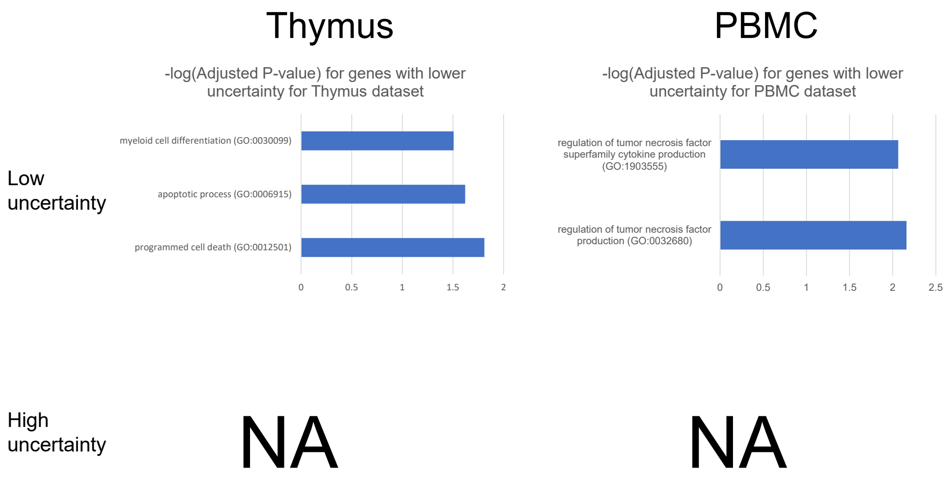


**Extended Data Fig. 8** Tissue-specific GO Enrichment Analysis for genes with different uncertainty levels. NA means we cannot detect clear and research-supported tissue-specific pathways in this gene group.

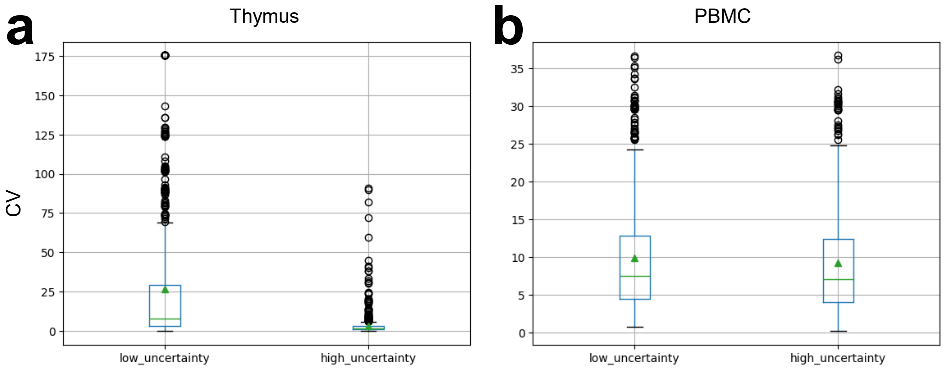


**Extended Data Fig. 9** Comparisons of coefficients of variation (CV) for genes with different uncertainty levels for (a) thymus dataset and (b) PBMC dataset.

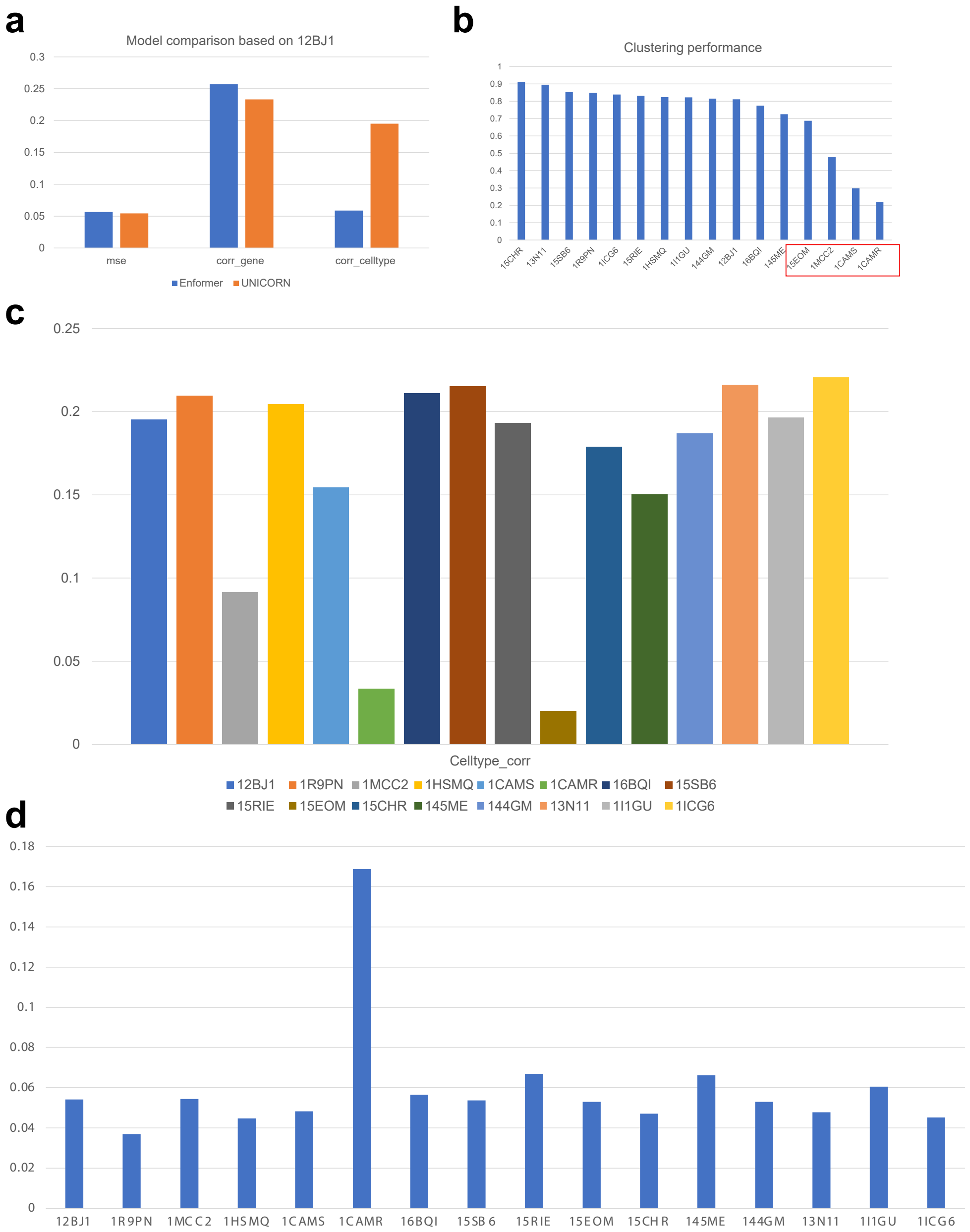


**Extended Data Fig. 10** Analysis of gene expression predictions for individualized scRNA-seq datasets. (a) The performance comparison between Enformer and UNICORN based on the sample 12BJ1. (b) The average clustering scores across different individuals based on the corresponding scRNA-seq datasets. (c) The results of correlation coefficients computed based on cell types across different samples. (d) The results of MSE computed based on genes and cells across different samples.

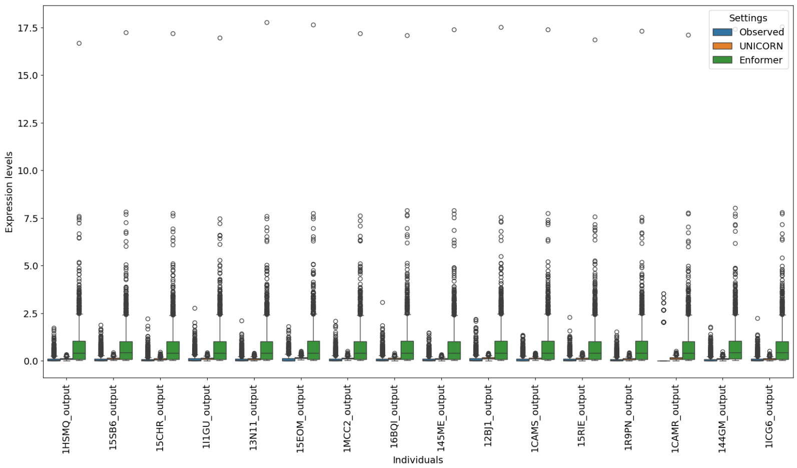


**Extended Data Fig. 11** Comparisons of gene expression levels across individuals. We assign different colors for the observed values and predicted values.

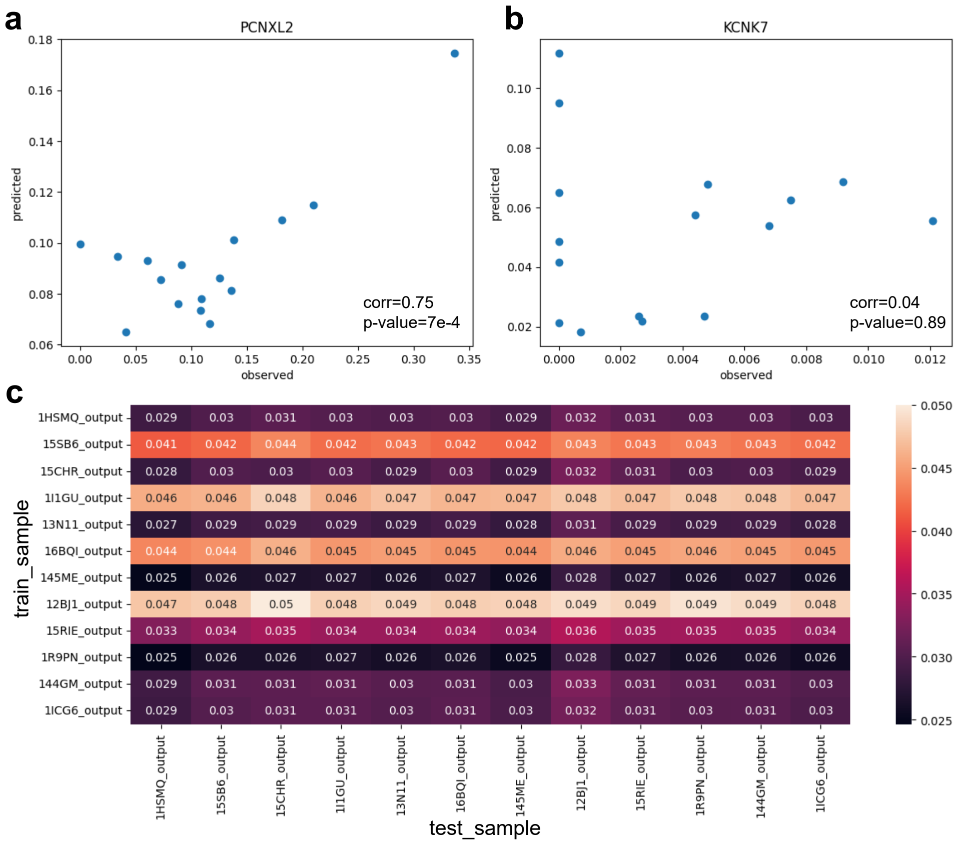


**Extended Data Fig. 12** Examples of genes with different prediction performances and the results of cross-individual prediction. (a) The correlation between observed PCNXL2 expression and predicted PCNXL2 expression (good example). (b) The correlation between observed KCNK7 expression and predicted KCNK7 expression (poor example). (c) The MSE value of cross-individual prediction result based on UNICORN. The rows representing training samples and the columns represent the testing samples.

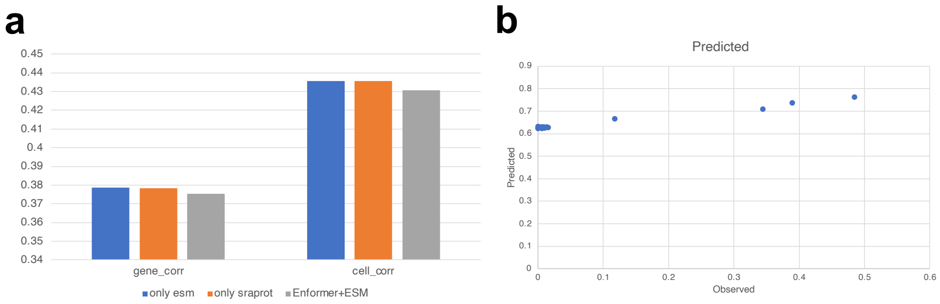


**Extended Data Fig. 13** Analysis the factors in affecting multi-omic prediction. (a) The ablation test for different approaches of embedding combination. Here ESM2 [62] and SaProt [78] are two models used to generate protein embeddings. (b) The scatter plot for visualizing the relationship between observed peak information and predicted peak information for different cell types measured in 10X Multi-Omic dataset.

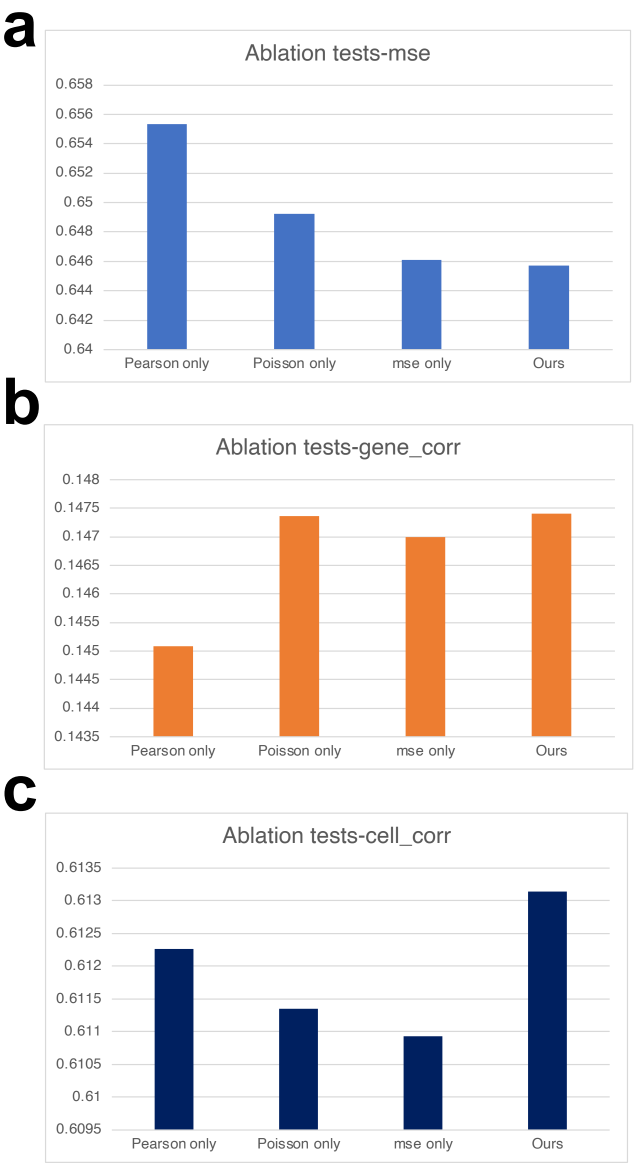


**Extended Data Fig. 14** Results of ablation tests for multi-modal prediction. We report the averaged score for each metric. (a) The correlation coefficients of genes under different choices of loss function components. (b) The correlation coefficients of cells under different choices of loss function components. (c) The MSE under different choices of loss function components.

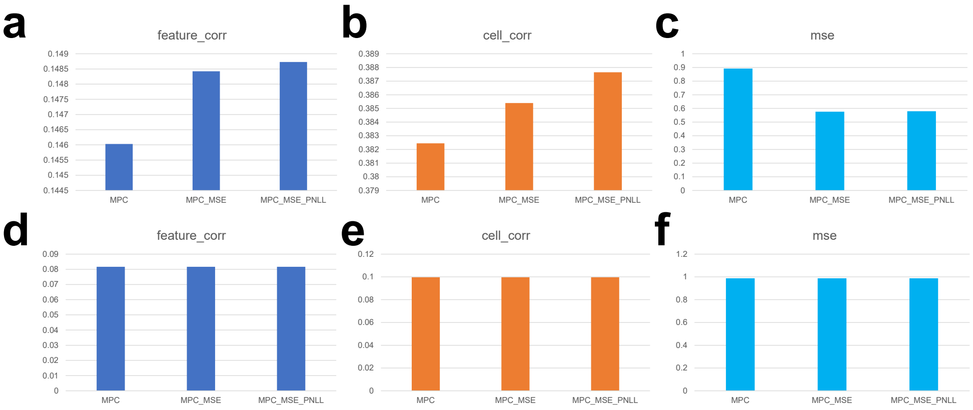


**Extended Data Fig. 15** Results of ablation tests for gene expression prediction. We report the averaged score for each metric. (a) The correlation coefficients of features under different choices of loss function components based on 10X Multiome data. (b) The correlation coefficients of cells under different choices of loss function components based on 10X Multiome data. (c) The MSE under different choices of loss function components based on 10X Multiome data. (d) The correlation coefficients of features under different choices of loss function components based on CITE-seq data. (e) The correlation coefficients of cells under different choices of loss function components based on CITE-seq data. (f) The MSE under different choices of loss function components based on CITE-seq data.

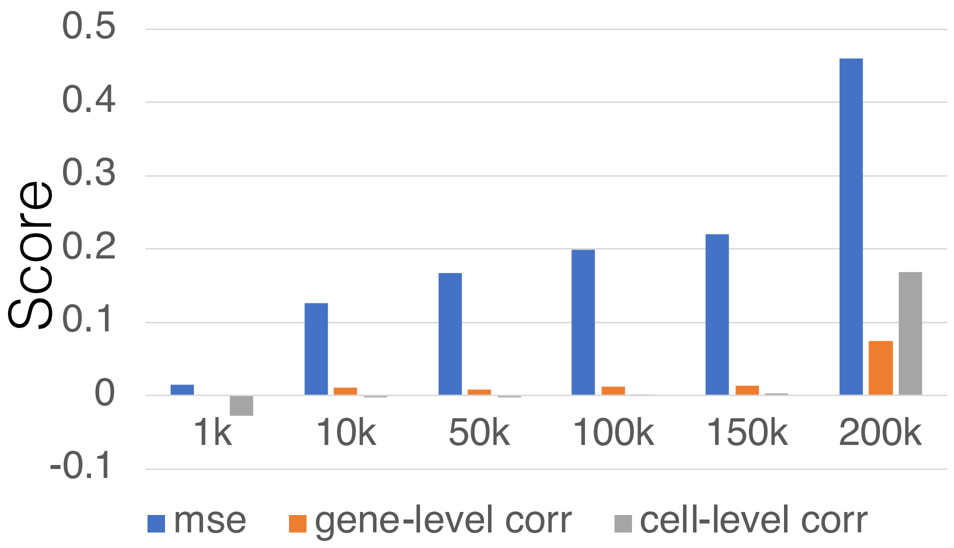


**Extended Data Fig. 16** Comparisons for single-cell gene expression prediction under different context lengths. We visualize three metrics (MSE, gene-level correlation score, and cell-level correlation score) by showing their averaged values under different context lengths from 1k to 200k.

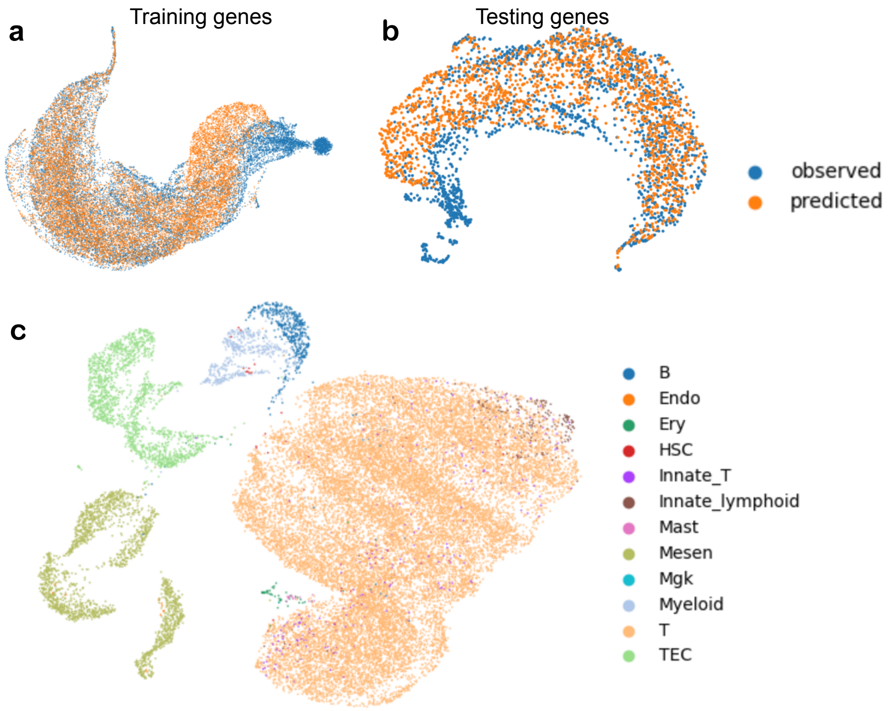


**Extended Data Fig. 17** The visualization results of atlas-level data prediction. (a) The UMAP plots for training gene embeddings from observed gene expressions and predicted gene expressions. (b) The UMAP plots for testing gene embeddings from observed gene expressions and predicted gene expressions. (c) The UMAP plots for cells with predicted expression levels from the thymus atlas dataset, colored by cell types.

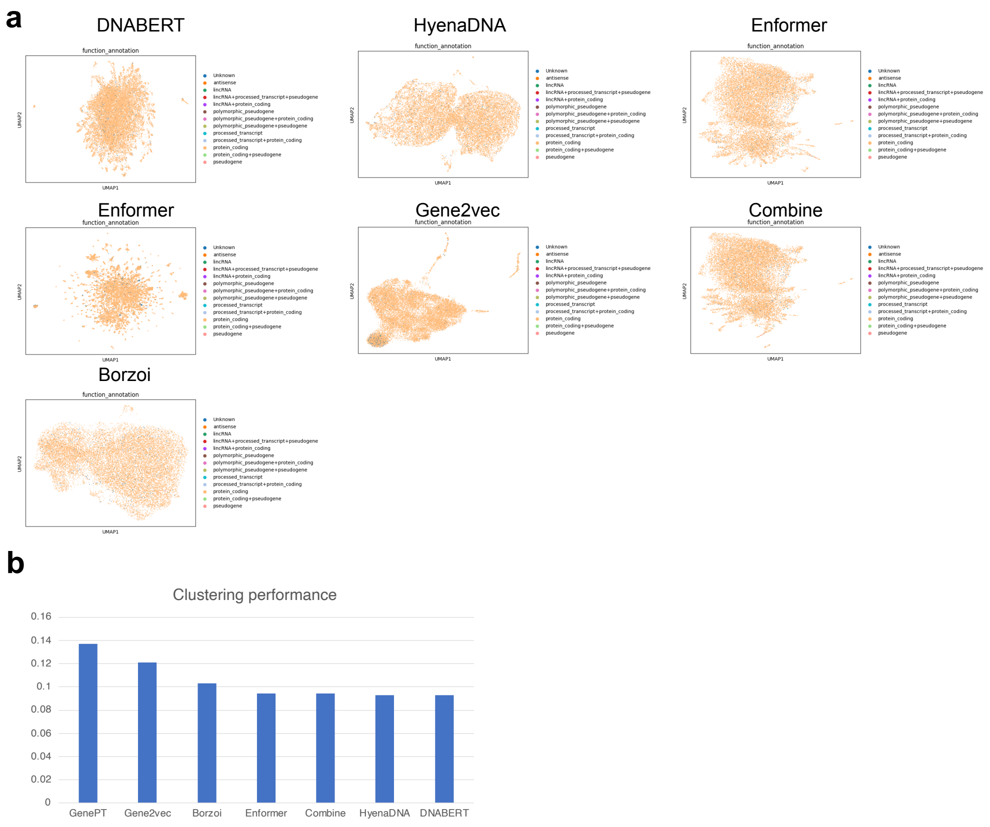


**Extended Data Fig. 18** Performances of clustering based on different gene embeddings. (a) The UMAP plots of different gene embeddings colored by functional annotations. (b) The averaged scores of different gene embeddings.

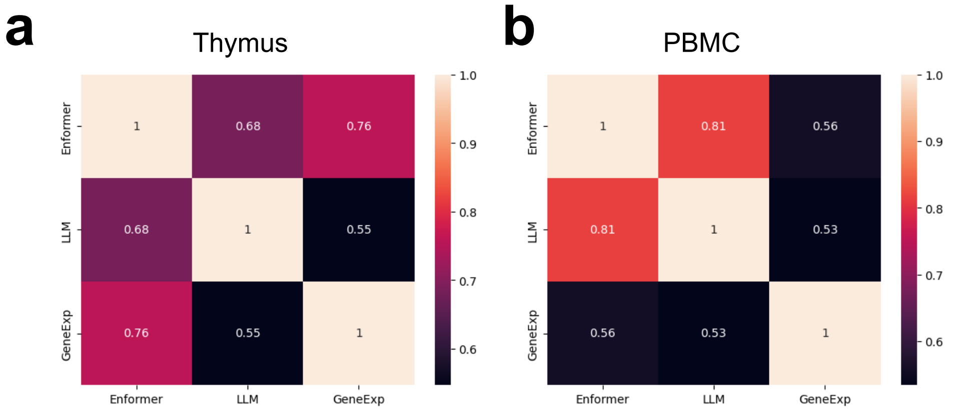


**Extended Data Fig. 19** Comparison of the gene-level similarity from Enformer, LLM, and gene expression profiles. (a) represents the results computed based on thymus dataset, and (b) represents the results computed based on PBMC dataset.

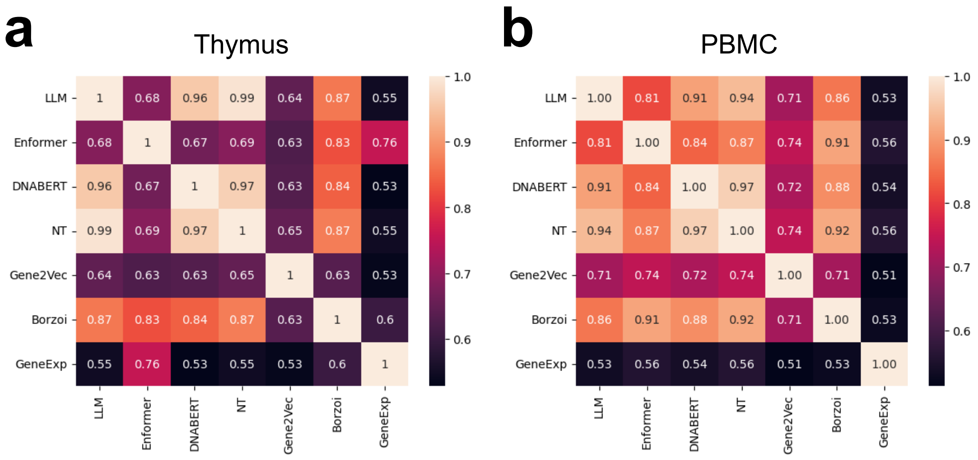


**Extended Data Fig. 20** Comparison of the gene-level similarity from all base models and expression profiles used in this manuscript. (a) represents the results computed based on thymus dataset, and (b) represents the results computed based on PBMC dataset.

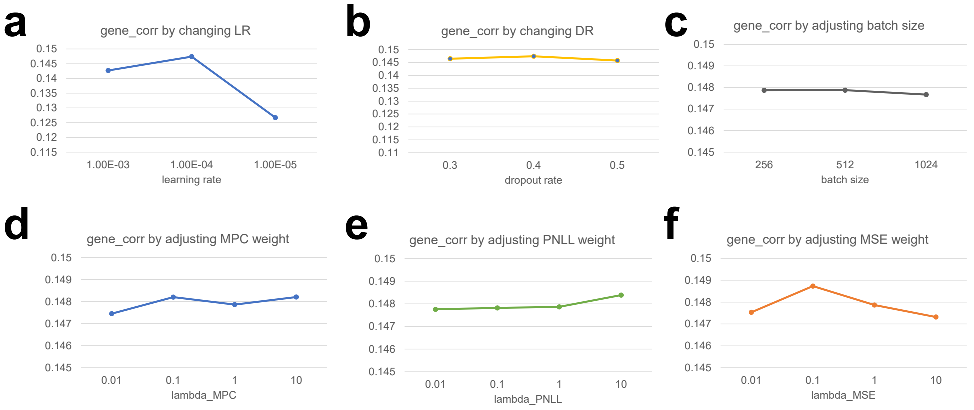


**Extended Data Fig. 21** Model performances under different settings of hyper-parameters. (a) The effect of changing LR. (b) The effect of changing DR. (c) The effect of changing batch size. (d) The effect of changing the weight of $\mathcal{L}_{MPC}$. (e) The effect of changing the weight of $\mathcal{L}_{PNLL}$. (f) The effect of changing the weight of $\mathcal{L}_{MSE}$.

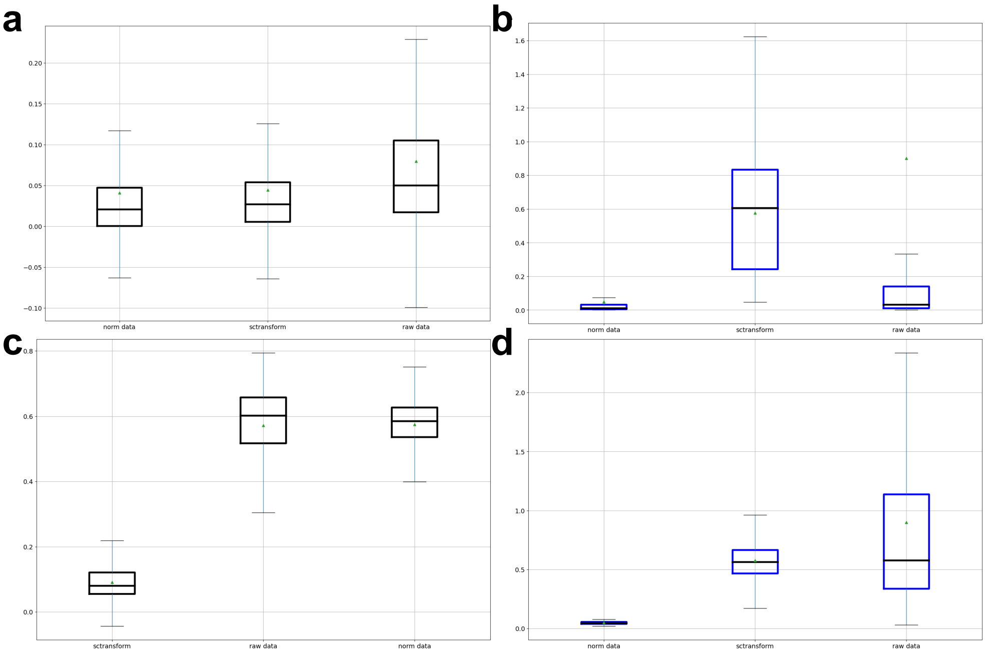


**Extended Data Fig. 22** Comparisons between raw data mode (raw data), log-normalized data mode (norm data), and sctransform data mode (sctransform). The triangle shape represents the mean value, and the black dashed line represents the median value. We report the results based on box plots. (a) Results of gene-level correlations across different methods based on the PBMC dataset. (b) Results of gene-level MSE across different methods based on the PBMC dataset. (c) Results of cell-level correlations across different methods based on the PBMC dataset. (d) Results of cell-level MSE across different methods based on the PBMC dataset.

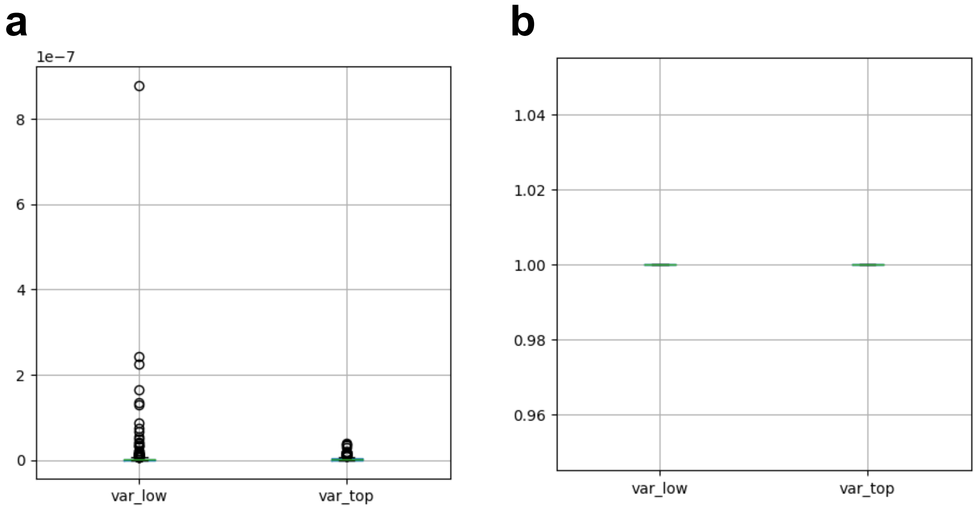


**Extended Data Fig. 23** The difference of metrics computed based on effective allele and non-effective allele. Here the legend *var_low* represent last 100 alleles and the legend *var_top* represents the top 100 alleles. The gene expression is computed at the pseudo-bulk level. (a) The distribution of MSE between the predicted gene expression and the observed gene expression. (b) The distribution of gene-level correlation coefficients between the predicted gene expression and the observed gene expression.
